# Large-scale two-photon calcium imaging in freely moving mice

**DOI:** 10.1101/2021.09.20.461015

**Authors:** Weijian Zong, Horst A. Obenhaus, Emilie R. Skytøen, Hanna Eneqvist, Nienke L. de Jong, Marina R. Jorge, May-Britt Moser, Edvard I. Moser

**Author notes:** Corresponding author. and (editorial correspondence).

## Abstract

We developed a miniaturized two-photon microscope (MINI2P) for fast, high-resolution, multiplane calcium imaging of over 1,000 neurons at a time in freely moving mice. With a microscope weight below 3g and a highly flexible connection cable, MINI2P allowed imaging to proceed with no impediment of behavior in half-hour free-foraging trials compared to untethered, unimplanted animals. The improved cell yield was achieved through a new optical system design featuring an enlarged field of view (FOV) and a new micro-tunable lens with increased *z*-scanning range and speed that allowed for fast and stable imaging of multiple, interleaved planes as well as 3D functional imaging. A novel technique for successive imaging across multiple, adjacent FOVs enabled recordings from more than 10,000 neurons in the same animal. Large-scale proof-of-principle data were obtained from cell populations in visual cortex, medial entorhinal cortex, and hippocampus, revealing spatial tuning of cells in all areas, including visual cortex.

**Highlights:** We developed a lightweight 2-photon miniscope for imaging in freely-foraging mice

Activity can be monitored in volumes of over 1,000 visual or entorhinal-cortex cells

A new *z*-scanning module allows fast imaging across multiple interleaved planes

Successive imaging from adjacent regions enables imaging from more than 10,000 cells

## Introduction

With the recent introduction of large-scale neural recording technologies for freely moving rodents, it has become possible to embark on the search for neural population codes underlying complex mammalian brain functions. New generations of silicon probes for electrophysiological recording (Jun et al., 2017; Steinmetz et al., 2021) and portable microscopes for calcium imaging (Ghosh et al., 2011; Skocek et al., 2018; Yanny et al., 2020; Zong et al., 2017) enable the simultaneous observation of activity in hundreds to thousands of identifiable neurons while animals perform experimental tasks, allowing researchers to identify the population dynamics of neural circuits, a goal that would be out of reach if the cells were recorded one by one or a few at a time.

Calcium imaging with portable microscopes is of particular interest for applications that require high anatomical resolution and the ability to distinguish between recorded cells based on genetic profile, soma location, or axonal projection pattern. Modern imaging protocols use genetically-encoded calcium indicators (GECIs), such as those belonging to the fluorescent GCaMP family (Chen et al., 2013; Shen et al., 2020), to optically monitor changes in intracellular free calcium across wide FOVs. The expression of GCaMP either through viral delivery or in transgenic animals allows neural activity to be monitored simultaneously at near-spike temporal resolution across large populations of individually identifiable neurons. For high-resolution imaging *in vivo*, investigators have mostly relied on stationary benchtop two-photon (2P) microscopes where mice are head-fixed under the objective. In these applications, behavioral testing is either limited to tasks where sensory stimulation and motor output can take place at a single location in the real world, or the head-fixed animal is tested in a virtual-reality environment under conditions that simulate the natural environment (Dombeck et al., 2007; Harvey et al., 2012). Head-fixed protocols are suboptimal, however, for studies that involve self-guided non-stationary behavior such as navigation, and they preclude, to a large extent, the identification of cells that are inherently spatial in their firing patterns, and that depend on proprioceptive-vestibular information, which is compromised by head-fixing (Minderer et al., 2016).

Due to the unsuitability of benchtop imaging for tasks that require unrestrained movement, investigators have tried, for almost two decades, to develop miniature 2P microscopes – 2P miniscopes – that integrate the 2P microscopés optical focusing and point-scanning components and can be carried on the head of freely moving animals. Portable devices that emerged from this work were connected to a remote stationary femtosecond laser source, photodetectors and acquisition equipment through cable assemblies containing optical fibers and electrical wires (Helmchen et al., 2013; Helmchen et al., 2001; Piyawattanametha et al., 2009; Sawinski et al., 2009; Zong et al., 2021; Zong et al., 2017). Early versions of these 2P miniscopes did not catch on, however, because they lacked the capacity to image sensitive calcium indicators such as GCaMP6, their scanning speed was slow, the miniscopes were heavy and difficult to carry for rodents, or optical cables were stiff and inflexible (Helmchen et al., 2001; Piyawattanametha et al., 2009; Sawinski et al., 2009). Calcium imaging in freely-moving animals took off, instead, with the invention of one-photon (1P) miniscopes where the optical sectioning and 3D imaging capability of 2P excitation was sacrificed to reduce size and to obtain a light-weight cable connection (Ghosh et al., 2011). While spatial resolution was compromised, the portable nature of 1P miniscopes vastly expanded the range of behaviors and scale of environments accessible to imaging experiments. With the 1P devices, activity can still be imaged at near-cellular resolution in sparsely active thin-layered structures such as CA1 of the hippocampus (Rubin et al., 2019; Ziv et al., 2013) or in denser structures if the expression of the indicator is kept sparse and thus only a small subset of all neurons is recorded (Glas et al., 2019). However, with 1P miniscopes, reliable single-cell resolution imaging is unattainable for applications that require capturing activity in densely labeled and active networks, due to the susceptibility to background fluorescence, optical aberrations, inability to detect small calcium transients, and lack of information in the *z* plane (Aharoni and Hoogland, 2019; Aharoni et al., 2019).

These constraints motivated attempts to develop a new generation of miniaturized 2P devices with resolution, speed and *z*-scanning capability comparable to that of 2P benchtop microscopes, and with a FOV close to that of 1P miniscopes (Li et al., 2021; Ozbay et al., 2018; Wallace and Kerr, 2019; Zong et al., 2021; Zong et al., 2017). Among these attempts were the 2P fiberscopes, where cable twisting and microscope weight were reduced by delivering fiber pulses and collecting fluorescence in a ‘double-claddinǵ fiber (Li et al., 2021; Rivera et al., 2011; Zhang et al., 2012). However, double-cladding fibers came at the expense of slow speed, low excitation and collection efficiency, and, in consequence, low cell yield. Of particular interest was the development of a new type of 2P miniscope that included (1) a hollow-core photonic-crystal fiber (HC-920) to deliver 920-nm femtosecond laser pulses, (2) a fast microelectromechanical systems (MEMS) scanner (Milanovic et al., 2004) for fast point scanning, and (3) a supple fiber bundle to collect the fluorescence (Zong et al., 2021; Zong et al., 2017). The first version of these miniscopes (Zong et al., 2017) weighed only 2.2 g but was not suitable for neural population imaging due to its small FOV (130×130 µm^2^). This was compensated for in a recent second-version upgrade equipped with a larger-aperture MEMS scanner as well as a custom low-magnification miniature objective and an electrically tunable lens (ETL) (Zong et al., 2021). With these improvements, the FOV increased to over 400×400 µm^2^ and 180 µm *z*-scanning capability could be added. Unfortunately, these changes increased the miniscope weight to ~5 g (4.2 g microscope, 0.9 g scope holder). This weight gain, along with the stiffness of the optical fiber bundle, interfered with the animals’ free movement, compromising the validity of any experiment in which mice are required to navigate through large environments for tens of minutes. Finally, imaging depth was improved with the development of three-photon (3P) miniscopes (Klioutchnikov et al., 2020) but the constrains on behavior remained (5 g microscope weight, stiff collection fiber bundle).

Here we present a new generation of 2P miniscopes, MINI2P, that overcomes the limits of previous versions by both meeting requirements for fatigue-free exploratory behavior during extended recording periods and satisfying demands for further increasing the cell yield by an order of magnitude. The improved thermal stability and speed of the integrated *z*-scanning module of MINI2P enables true 3D functional imaging. We supplement these advances with a novel technique where adjacent FOVs are stitched together over successive recordings, increasing the yield of successively recorded cells to over 10,000 in one animal. MINI2P is open source and can be built in most imaging laboratories with detailed building instructions provided in this paper. The performance and reliability of MINI2P are validated by proof-of-principle recordings of spatially tuned neurons in three brain regions. In an accompanying research article in this issue (Obenhaus et al, accompanying manuscript) we used MINI2P to obtain large scale recordings of dense populations of neurons in the medial entorhinal cortex (MEC) and the neighboring parasubiculum (PAS), which in turn helped us determine the topographical organization of all known, spatially modulated cell types in MEC and PAS.

## Results

### Exploratory behavior is affected by miniscope weight and cable flexibility

To determine how sensitive the behavior of a mouse is to the weight of the head-mounted miniscope and the flexibility of the connection cables, we first investigated the impact of this pair of variables on free-foraging behavior in 10 adult mice (5 wildtype and 5 transgenic; weight, 28.8 ± 2.2 g (mean ± S.D.)). Using dummy miniscopes (with no imaging), we compared the most recently published second version of the 2P miniscope (Zong et al., 2021) to that of a less intrusive pilot microscope body with more flexible cables that we thought could be realistically produced by changes in miniscope and cable design. The most recently published version (Zong et al., 2021) had a weight of ~5 g and was used with a standard 1.5 mm ‘thick’ cable assembly (5g-T; cable assembly refers to an HC-920 photonic-crystal fiber, a 6-core electric wire, and a 1.5-mm-diameter fiber bundle of one to two hundred 0.05-mm-diameter glass fibers). In the pilot model, the scope weight was reduced to 3 g and the diameter of the cable assembly to less than one-half, or 0.7 mm (‘thin’ cable; 3g-t). Animals with 5g-T and 3g-t dummy miniscopes and cables, or no miniscope or tether at all (control), ran for scattered food crumbles in sessions of 30 min length inside an 80 cm wide, square open-field box (Figure 1). The sequence of 5g-T, 3g-t and control experiments was randomized, with one condition per day. A counterweight was used to balance the weight of the cable assembly.

**Figure 1.**
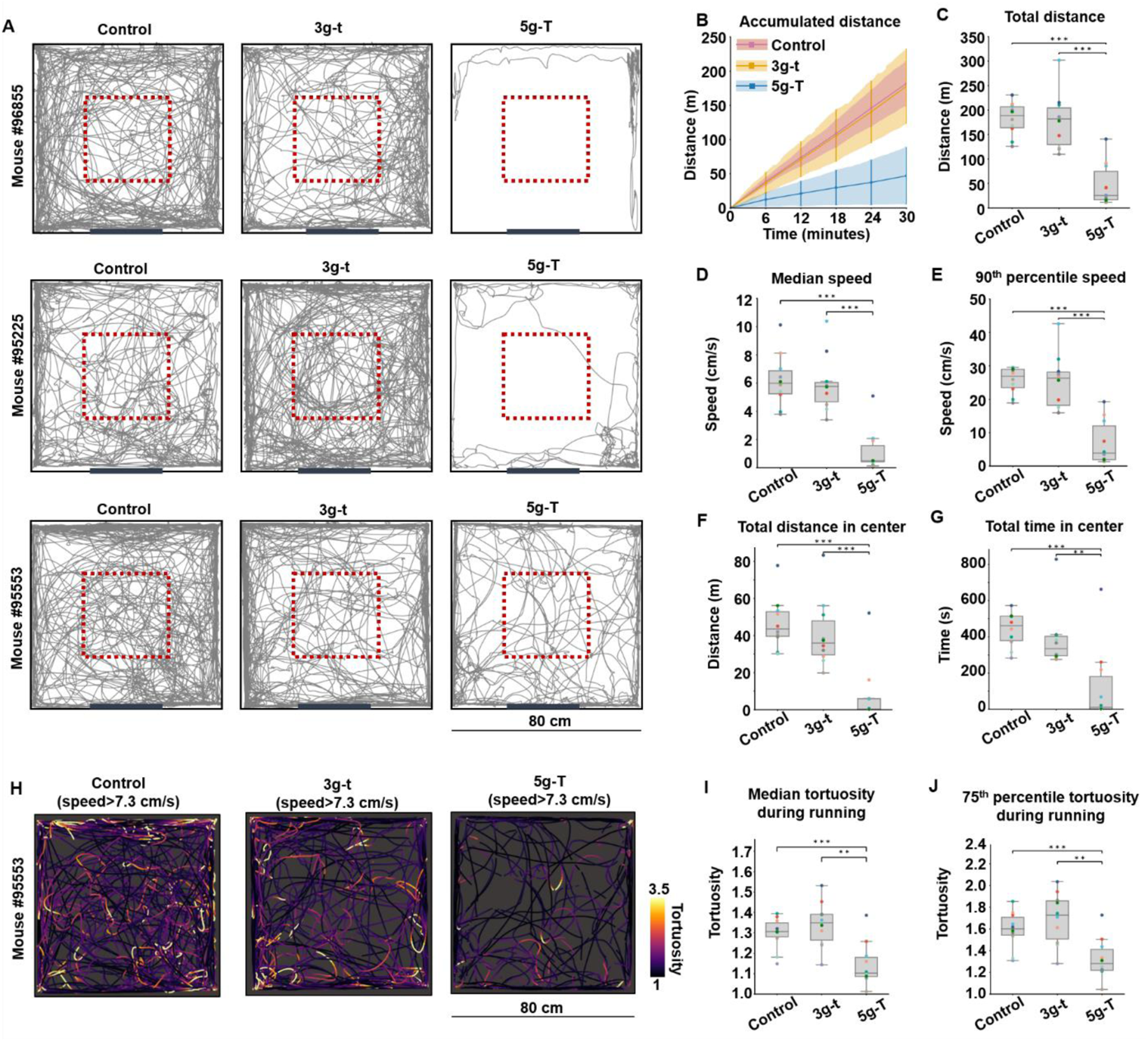
Flexible cable and low weight of MINI2P significantly improves free-foraging behavior of mice. (**A**) Representative trajectories of 3 animals over 3 sessions each, where mice were running in 80 cm wide square open field boxes: Control (no dummy or cable), 3g dummy + thin connection cable assembly (3g-t) and 5g dummy + thick connection cable assembly (5g-T). Animals were ranked based on their total running distance in the 5g-T experiment; trajectories are shown for the 20th, 50th and 80th percentile of all animals on the 5g-T session (top to bottom). Red dashed boxes in the middle show 30 × 30 cm square used in calculations of time and distance in the center ((**F**) and (**G**)). Thick black line at the bottom shows the position of a cue card for orientation. (**B**) Accumulated distance over 30 min of continuous running. Lines denote mean value across 10 animals, vertical bars show S.D. at 6 discrete time points (6th min, 12th min, etc.), and shaded region indicates S.D. across all time points. Note similarity between 3g-t and control group. (**C**) Box plot showing total distance travelled in each condition (30 min each). Horizontal line indicates median, boxes interquartile range, and whiskers 1.5 times interquartile range. Colored dots represent individual animals (colors remain the same between sessions). Conditions are compared using a Friedman test followed by Tukey post hoc correction, *p < 0.05, **p < 0.01, and ***p < 0.001. (**D** and **E**) Differences in median speed (**D**) and 90th percentile speed (**E**) in the three conditions (box plot). Symbols as in C. (**F** and **G**) Differences in distance travelled (**F**) and time spent (**G**) within the central 30 cm square of the open field (red dashed boxes in A). (**H-J**) Reduced turning behavior in 5g-T condition. (**H**) Representative trajectories during high-speed running events (frames with speed lower than 75th percentile (<7.3 cm/s) were removed). Color scale shows momentary tortuosity (curvedness) of animaĺs trajectory during high-speed movement. (**I-J**) Median tortuosity (**I**) and 75th percentile of tortuosity (**J**) during running (speed>7.3cm/s) in each condition. See also Figure S1.

The experiments showed that animals carrying dummy miniscopes mimicking previous-generation versions with a thick cable assembly (5g-T) ran shorter overall distances (Figure 1A-C), at a slower speed (Figure 1D and 1E), than control animals with no dummy or cable. The distance and speed of 3g-t animals matched that of untethered control animals (Friedman test on data from all three experimental conditions, total running distance: χ^2^ = 16.8; median running speed: χ^2^ = 15.8; 90^th^ percentile running speed: χ^2^ = 16.8, all with p < 0.0004; all post-hoc Tukey tests for 5g-T vs control and 5g-T vs 3g-t: p < 0.001; 3g-t vs control: all p >0.90). Differences in distance and time were particularly pronounced in the center of the box (30 x 30 cm; Figure 1F and 1G; Friedman test, distance: χ^2^ = 15.8; time: χ^2^ = 12.2, both with p < 0.0022; all post-hoc Tukey tests for 5g-T vs control and 5g-T vs 3g-t: p < 0.004; all 3g-t vs control: p > 0.691). Weight and cable thickness also affected the animals’ ability to turn during running. For frames with high running speeds (above the 75^th^ percentile of all speed values summarized over all animals), we estimated the momentary tortuosity or curvedness of the animaĺs trajectory by calculating, in sliding segments, the ratio of the length of the trajectory over the straight-line distance between the start and end point in each segment (Figure S1A). The tortuosity of the animal’s trajectory was reduced in the 5g-T group compared to 3g-t and control animals but was not different between 3g-t and control (Figure 1H-J; Friedman test, χ^2^ = 12.6, p = 0.0019; post-hoc Tukey tests, 5g-T vs control: p =0.006, 5g-T vs 3g-t: p = 0.001; 3g-t vs control: p = 0.598). The time spent turning (frames above 75^th^ percentile of all tortuosity values summarized over all animals), as well as running speed during these frames, were similarly reduced in the 5g-T condition compared to the other two conditions, whereas no difference appeared between 3g-t and control (Figure S1B-E). Finally, experiments addressing the impact of weight and cable flexibility separately showed that the animaĺs movement was more affected by the thickness or stiffness of the connection cable assembly than the weight of the miniscope (Figure S1F-Q). Taken together, these pilot observations suggest that a head-mounted miniscope with a weight equal to or less than 3g and with a 0.7 mm cable assembly would be sufficient for maintaining the natural free-foraging behavior of mice in 2P imaging experiments.

### Weight and cable flexibility: MINI2P satisfies criteria for unrestrained movement

Keeping in mind the constraints on weight and cable stiffness, we introduced several changes to the design of the miniscope and cable assembly to avoid impeding the animals’ exploratory behavior. We set out to reduce the miniscope weight to below 3 g and to engineer a cable no thicker than 0.7 mm, to mimic the 3g-t condition of the behavior experiment, which we showed had no detectable impact on the animals’ movement pattern.

The heaviest part in the second-version of the 2P miniscope (Zong et al., 2021) is the *z*-scanning module, consisting of an electrically tunable lens (ETL, ~1.3 g, Figure S2A), which along with relay optics makes up 42% of the total miniscope weight (1.8g of 4.2g; Table 1, Part 1). To decrease the overall weight while maintaining the large *z*-scanning range, we developed a new type of miniature *z*-focusing device based on a nanotech micro-tunable lens (Tlens®, Polight, Horten, Norway) (Farghaly et al., 2016) and named it μTlens (Figure 2C, 2D and Figure S2A-C). We assembled a quartet μTlens (4 flat lenses stacked together, Figure 2C), which weighed only 0.06 g, spanning a volume of 4.5×4.5×2.2 mm^3^ (Figure S2A). The quartet μTlens had a response time of less than 0.4 ms (Figure S2B), an optical power of ~75 diopter (dpt) (Figure S2C), and a *z*-scanning range of 240 μm, which is 60 μm more than that of the previous version of 2P miniscope (Zong et al., 2021) (Figure 2D). The small size of the μTlens allowed it to be mounted near the MEMS scanner without relay optics (Figure 2A and Supplementary Document, Section 1). The static driving technology of the MEMS piezo minimizes the driving current and thereby provides high thermal stability to the μTlens. No temperature increase or focus drift was observed when the focus was switched at 15 Hz or 40 Hz between multiple imaging planes, or set at max optical power (+51 dpt, or 240 μm) for up to 1 hour of continuous recording (see also Supplementary Document, Section 4). In contrast, a temperature increase of over 20 degrees Celsius, accompanied by drift of the focal plane, was noticed after less than 10 min imaging with the previously used ETL when applied at full optical power (±30 dpt) and driving current (±200 mA) (Supplementary Document, Section 4).

**Figure 2.**
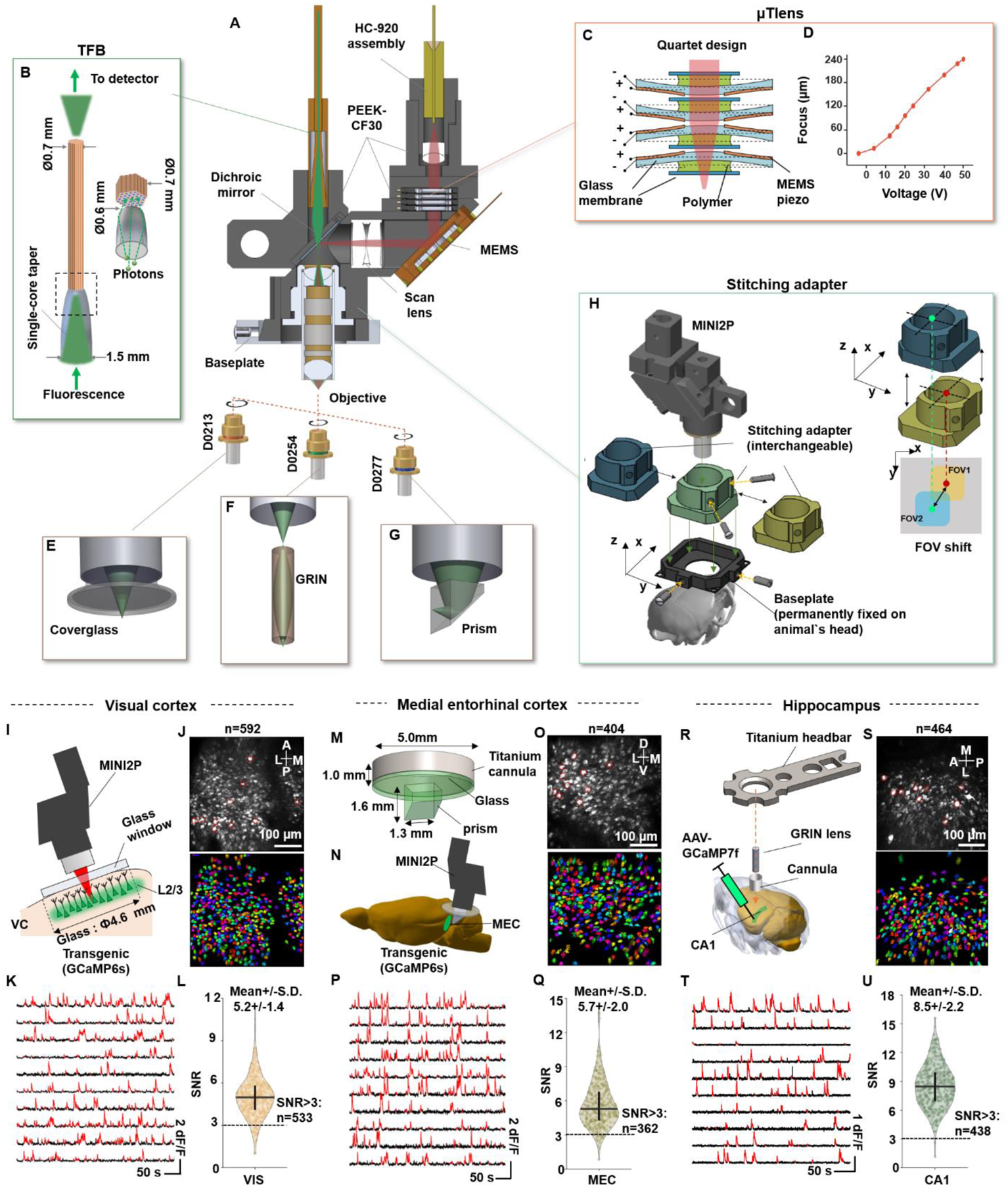
New features of MINI2P open possibilities for large-scale 2P calcium imaging in a variety of brain regions. (**A**) Section-view of a MINI2P with key components. (**B**) Structure of the tapered fiber bundle (TFB). (**C**) Quartet design of µTlens consisting of four stacked piezo-membrane lenses. (**D**) Axial scanning range of the MINI2P with quartet µTlens. (**E-G**) Three customized objectives are available for different imaging applications. (**H**) A series of stitching adapters added between the MINI2P scopebody and the baseplate allow shifting of the FOV even after the baseplate is permanently fixed on the head of the animal. Accurate and repeatable FOV shifts in up to 400-μm steps are achieved by replacing stitching adapters. (**I-L**) Imaging in visual cortices (VC) in a CaMKII-tTA/tetO-GCaMP6s transgenic mouse. The mouse was running in an open enclosure as in Figure 1. (**I**) Schematic showing mounting of MINI2P microscope on top of cover glass to image L2/L3 neurons in VC. (**J**) Example MINI2P imaging in VC where 592 neurons were extracted by Suite2P in a single plane of one FOV. Top: maximum intensity projection of all recorded images (30 min). Bottom: 592 raw extracted neurons randomly colored for visualization. (**K**) Ten example calcium signals from neurons marked by red circles in top panel of (**J**). Red parts of traces identify suprathreshold calcium transients. (**L**) Violin plots showing SNR of extracted cells in the lower panel of (**J**). The outline of the violin plot shows probability density smoothed by a kernel density estimator (bandwidth: 10% of the data range). Interquartile range and median are indicated with vertical and horizontal lines, respectively. Dashed line shows threshold for cell inclusion (SNR=3). The number of cells above threshold is indicated. The symbols apply to all subsequent violin plots. (**M-Q**) Imaging in the medial entorhinal cortex (MEC). Symbols as in (I to L). (**M**) Schematic of prism assembly for MEC imaging in a CaMKII-tTA/tetO-GCaMP6s transgenic mouse. (**N**) Schematic showing position of MINI2P microscope for MEC imaging. The prism was inserted along the dorsoventral surface of MEC, between the cerebrum and the cerebellum, allowing activity in L2/L3 of this brain area to be imaged from the back. (**O**) Example of MINI2P imaging in MEC where 404 neurons were extracted by Suite2P in a single plane of one FOV. (**P**) Ten example calcium signals from the neurons marked by red circles in upper panel of (**O**). (**Q**) SNR of extracted cells in (**O**). Dashed line shows threshold for cell inclusion (SNR=3). The number of cells above threshold is indicated. (**R-U**) Imaging in hippocampal area CA1 after local injection of AAV-GCaMP7f in a wild-type mouse. (**R**) A 1.0-mm GRIN lens (with a 1.8mm guide cannula) was implanted in the left hemisphere to access CA1 from the top of the brain. A tunnel of overlying cortex was removed to enable the GRIN lens to be placed at the surface of CA1. A titanium headbar was added after the GRIN implantation for fixing the head of the animals and mounting the microscope. (**S**) Top: Example of MINI2P imaging in CA1. Bottom: 464 neurons extracted by Suite2P in one FOV. (**T**) 10 example calcium signals from the neurons marked by red circles in the top panel of (**S**). (**U**) SNR of extracted cells in (**S**). Dashed line shows threshold for cell inclusion (SNR=3). The number of cells over the threshold is indicated. See also Table 1 and Figure S2

**Table 1.** Summary of MINI2P features. Related to Figure 2. Part 1: Comparison of quartet µTlens and Optotune EL-3-10. Part 2: Parameters of the two types of MEMS scanners, MEMS-L and MEMS-F. Part 3: Parameters of three types of objective. Part 4: Comparison of MINI2P to the latest published 2P miniscope.

Part 1: μTlens v.s. EL-3-10
| | μTlens (quartet) | EL-3-10<br>(with relay optics) |
| --- | --- | --- |
| Weight: | 0.06 g | 1.8 g |
| Size: | 4.5x4.5x2.2 mm<sup>3</sup> | 10x10x15 mm<sup>3</sup> |
| Response: | <0.4 ms | 1.5 ms |
| Optical power: | -23 to 52 dpt (75 dpt) | -30 to 30 dpt (60 dpt) |
| Heating effect: | Negligible | High |

Part 2: two types of MEMS
|  | Large-angle MEMS<br>(MEMS-L) | Fast MEMS<br>(MEMS-F) |
| --- | --- | --- |
| Type | A3I12.2-1200AL | A7M10.2-1000AL |
| Mirror size | 1.2 mm | 1.0 mm |
| First resonant frequency | 2.6 kHz | 4.8 kHz |
| Max Scanning angle: | $\pm 5.2$ degree | $\pm 4.5$ degree |
| Working frequency | 2 kHz | 5.6 kHz |
| Imaging speed | 15 Hz (256x256) | 40 Hz (256x256) |

Part 3: three objectives
|  | Domilight D0213 | Domilight D0254 | Domilight D0277 |
| --- | --- | --- | --- |
| Max image area | $\Phi 550 \mu\text{m}$ | $\Phi 550 \mu\text{m}$ | $\Phi 550 \mu\text{m}$ |
| N.A. | 0.5 | 0.5 | 0.45 |
| Working distance | 1.00-mm water+0.17mm glass | 0.58 mm air | 0~0.7mm water/air+1.5~2mm glass |
| XY resolution | $1.15 \pm 0.15 \mu\text{m}$ | $1.21 \pm 0.13 \mu\text{m}$ | $1.24 \pm 0.17 \mu\text{m}$ |
| Z resolution | $17.80 \pm 0.85 \mu\text{m}$ | $14.53 \pm 2.12 \mu\text{m}$ | $12.8 \pm 0.28 \mu\text{m}$ |

Part 4: MINI2P v.s. the previous version of 2P miniscope
|  | MINI2P | The previous version of 2P miniscope (Zong et al., 2021) |
| --- | --- | --- |
| MAX FOV | $\sim 0.25 \text{ mm}^2$ ($\sim 0.5 \text{ mm} \times 0.5 \text{ mm}$, MINI2P-L, single FOV)<br>$\sim 0.17 \text{ mm}^2$ ($\sim 0.42 \text{ mm} \times 0.42 \text{ mm}$, MINI2P-F, single FOV)<br>$> 4 \text{ mm}^2$ ($5 \times 5 \times 0.16 \text{ mm}^2$, field stitching) | $\sim 0.17 \text{ mm}^2$ ($\sim 0.42 \text{ mm} \times 0.42 \text{ mm}$) |
| Speed | 15Hz (MINI2P-L, 256 line)<br>40Hz (MINI2P-F, 256 line) | 20Hz (256 line) |
| Weight | 2.4 g | 4.2 g |
| Collection fiber bundle | Diameter: 0.7 mm | Diameter: 1.5 mm |
| Z-scanning module | Range: 240 $\mu\text{m}$,<br>Response: <0.4 ms,<br>Weight: 0.06 g | Range: 180 $\mu\text{m}$,<br>Response: <1.5 ms,<br>Weight: 1.8 g |
| Objective | Dry, water and glass | Only water |
| Building complexity | Easy | Complex |
| System size | Portable | Stationary |
| Accessibility | Fully open-source | Needs customized components |

The second-heaviest part in the previous 2P miniscope is the microscope shell (~1.5g or 36%).Instead of aluminum, we used carbon-filled polyether ether ketone (PEEK-CF30), which brought the weight of the whole shell down to 0.8 g, or one-half of the weight of the aluminum shell of the previous version (Zong et al., 2021), without loss of mechanical strength or machining accuracy (Figure 2A). With the new miniature *z*-focusing device and the new shell material, MINI2P reached a total weight of only 2.4 g. With the addition of a stitching adapter and baseplate (0.6 g, see below), the total weight was equal to the dummy miniscope in the 3g-t condition of the behavioral pilot experiment.

In addition to reducing weight, we designed a thinner, lighter, and more flexible optical cable to prevent disruption of free-exploration behavior. In order to maintain the efficiency of fluorescence collection, the collection fiber bundle needs a minimum diameter. Imaging in a uniform fluorescence sample showed that, with a 0.7 mm-diameter supple fiber bundle (SFB), less than about 20% of the signal could be collected compared to a 1.5 mm SFB of similar material used in the two previous 2P miniscope versions (Zong et al., 2021; Zong et al., 2017) (Figure S2D and S2E). For *in vivo* imaging, the collection efficiency of such a 0.7 mm fiber bundle would be even lower due to extensive light scattering in brain tissue. To improve the collection efficiency, we thus designed and fabricated a new type of tapered fiber bundle (TFB) consisting of a 7-mm long tapered glass rod followed by a fiber bundle with a diameter of 0.7 mm (Figure 2B and Figure S2F). In the TFB, scattered fluorescence photons are collected by the wider end of the tapered glass (diameter 1.5 mm) and, after multiple total internal reflections in the glass rod, the photons are coupled into the fiber bundle from the tapered end (diameter 0.6 mm). Because the tapered glass also has a focusing effect, no collecting lens was required. The reflection index and the outline curvature of the tapered glass rod were carefully designed to match the numerical aperture (NA) of the objective (NA 0.15 at imaging plane) and the fiber bundle (NA 0.5), bringing the collection efficiency of the TFB to a level nearly identical to that of the 1.5 mm SFB and far beyond that of the 0.7 mm SFB (Figure S2D middle and Figure S2E). The TFB matched the characteristics of the light and flexible cable assembly used in the behavioral pilot experiments (Figure 1 and Figure S1) and should therefore be able to support extended recordings during unconstrained movement.

### MINI2P enables large-scale high-resolution imaging in diverse brain structures

In order to support different experiments requiring either faster imaging speed or a larger FOV, we developed two versions of the new 2P miniscope, MINI2P-L and MINI2P-F. MINI2P-L uses the same MEMS scanner (MEMS-L in Figure S2G and Table 1, Part 2) as the previous 2P miniscope (Zong et al., 2021) but with a new scan lens (focal length 5 mm). The larger scan lens allowed for imaging of a larger FOV with a 15 Hz frame rate at 256 × 256 pixels (3 miniscopes were used for tests reported with this version of MINI2P; maximum FOV sizes ranged from 500×500 µm^2^ to 510×510 µm^2^). MINI2P-F, in contrast, has a new type of low Q-factor MEMS scanner (MEMS-F in Figure S2G and Table 1, Part 2) and underwent further optimizations to scan frequency (Figure S2G) and angle (Figure S2H) (see also STAR Methods). This provided faster speed (40 Hz frame rate at 256 × 256 pixels) at the cost of a somewhat smaller FOV (2 miniscopes were used for the tests reported with this version of MINI2P; maximum FOV size ranged from 410×410 µm^2^ to 420×420 µm^2^).

Because scattering of fluorescent signals in the brain limits the imaging depth, different optical access methods are required for different brain regions, with choices depending on the depth and orientation of the target tissue (Bocarsly et al., 2015; Holtmaat et al., 2009; Low et al., 2014; Yang and Yuste, 2017; Ziv et al., 2013). Thus, we tailored a series of objectives to MINI2P, designed, respectively, for imaging through a thin glass window, a GRIN lens and a thick prism (Figure 2E-G). The three objectives are interchangeable with the same threads so that different applications can be implemented in one MINI2P system. Resolution and maximum FOV of the three objectives are similar (Table 1, Part 3; Supplementary Document, Section 2).

To validate the recording quality of MINI2P with different access methods, we monitored calcium activity in freely-moving mice in which the miniscope was used with three different objectives to target visual cortex (VC, via a glass window; Figure 2I-L; Video S1, Part I), medial entorhinal cortex (MEC, via a prism; Figure 2M-Q; Video S1, Part II), and hippocampus (CA1, via a GRIN lens; Figure 2R-U; Video S1, Part III). Using *Suite2p*, an open-source cell detection and signal extraction pipeline (Pachitariu et al., 2017), we were able to detect hundreds of neurons in a single FOV in all three regions (592 cells in VC, 404 in MEC, and 464 in CA1). Signal-to-noise ratios (SNR) were high both in transgenic mice expressing GCaMP6s broadly in excitatory neurons (mouse line: CamKII-tTA/tetO-GCaMP6s for all VC and MEC data, mean SNRs ± S.D: 5.2±1.4 for VC, and 5.7±2.0 for MEC) and in wild-type mice pre-injected with AAV1-syn-GCaMP7f (CA1 data, mean SNRs ± S.D: 8.5±2.2). A total of 90% of the cells in VC (533 of 592), 90% in MEC (362 of 404), and 94% in CA1 (438 of 464) were above a SNR threshold of 3 (Figure 2L, 2Q and 2U). After more than 9 weeks of recording in VC and MEC, we observed a moderate decrease in cell yield (39% and 21%, respectively) but no decline in SNRs (Figure S2I-L). Individual neurons in VC could be followed reliably across months (Figure S2M). Taken together, the recordings demonstrate the versatility and applicability of MINI2P to a wide range of commonly used implantation methods for *in vivo* imaging.

### Increasing the cell yield to thousands

We explored three major strategies to increase the number of recorded cells (Figure 3A). In the first strategy, we enlarged the FOV. When the FOV was kept at 420×420 µm^2^ with MINI2P-L, equivalent to that of the previous second generation of 2P miniscope (Zong et al., 2021), we obtained a cell yield of 225±115 per FOV (mean ± S.D.; Figure 3B left and Table S1, Part 1; n=6 mice, 6 FOVs, 3 different brain regions). Then, by expanding the FOV by a factor of 1.4, to over 500×500 µm^2^ with the same MINI2P-L, we were able to record 335±62 cells per FOV (Figure 3B right and Table S1, Part 1; n=3 mice, 4 FOVs, 3 different brain regions).

**Figure 3.**
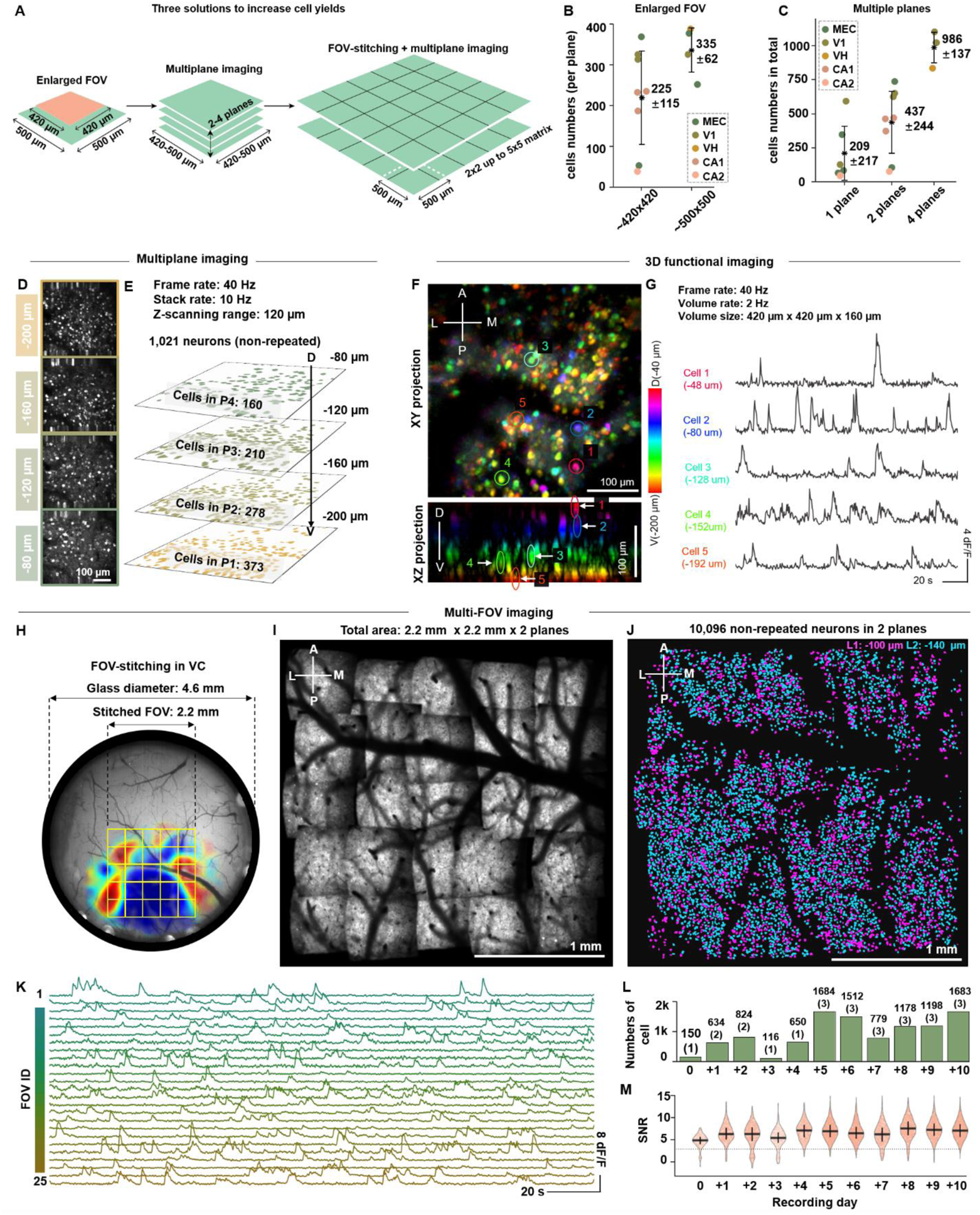
Increasing the cell yield of MINI2P to thousands. (**A**) Three steps to extend the FOV and increase the cell yield: enlargement of FOV, multiplane imaging, and stitching of neighboring FOVs with multiplane imaging. (**B**) Compared to the previous generation of miniscope (Zong et al., 2021), cell yields increased 150% when the FOV was enlarged from 420×420 μm2 to 500×500 μm2 (mean cell numbers ± S.D are indicated; 420×420 μm2: n=7 mice, 7 FOVs, 4 different brain regions; 500×500 μm2: n=3 mice, 4 FOVs, 3 brain regions, recorded with MINI2P-L, see able S1 for further details). Black stars indicate mean value. Color dots denote single recordings; color indicates brain region of each recording. VH represents higher-order visual cortex including LM, AL, RL, AM, and PM. Same definition applies in C. See also Table S1 part 1. (**C**) Single plane imaging gave an average yield of 209±217 cells (mean ± S.D; n=6 mice, 6 recordings, 3 different brain regions, MINI2P-L was used, but the FOV was set to 420×420 μm2). Two-plane imaging increased cell yields by 210 %, (n=7 mice, 8 FOVs, 5 brain regions, recorded with MINI2P-L but the FOV was set to 420×420 μm2). Four-plane imaging increased yields by 440% (n=2 mice, 3 FOVs, 2 brain regions, one recorded with MINI2P-L, the rest with MINI2P-F) compared to single-plane imaging. See also Table S1 part 1. (**D** and **E**) 4-plane imaging increased cell yield to over one thousand. (**D**) Maximum intensity projection of near-simultaneous 4-plane imaging in VC (planes scanned sequentially; 25 ms per plane). Plane interval 40 µm, frame rate 40 Hz, stack rate 10 Hz, recorded with MINI2P-F. (**E**) Hundreds of neurons could be extracted from each plane in (**D**). (**F** and **G**) 3D imaging of calcium activity in hundreds of VC neurons in freely-moving mice. Volume size: 420×420×160 µm3, volume rate: 2 Hz, recorded with MINI2P-F. (**F**) xy- and xz- max projection of 3D stack data (20 planes). A 4.8x bilinear interpolation was applied in the z-direction, resulting in 96 planes, to achieve isotropic voxel size. Color code indicates imaging depth measured from the surface of cortex. (**G**) Calcium signals of 5 neurons randomly selected from different imaging depths, marked by colored circles in (**F**). The imaging planes that were chosen to extract the calcium signals are indicated with arrowheads in the bottom panel in (**F**) (**H-M**) FOV stitching increased the cumulative cell yield across successive sessions to over 10,000 in VC. (**H**) Schematic of 4.6 mm diameter chronic window and matrix of 5×5 stitched FOVs on the left hemisphere. The outline of stitched FOVs (yellow squares) is superimposed on a retinotopic map extracted by wide-field calcium imaging, using an automated and quantitative method to demarcate visual cortical areas based on retinotopic organization (Zhuang et al., 2017). Color indicates the sine value of the angle between vertical (**V**) and horizontal (**H**) retinotopic mapping gradients (red: 1; blue: −1). Visual areas can be identified by similar color (red or blue) in the sign map. (**I**) Average image (from plane 2) showing stitched matrix of 5×5 FOVs covering a total of 2.2 × 2.2 mm2. (**J**) 2D projection of the 10,096 neurons in (**I**) shows separation of neurons in different planes (blue: −140 µm; magenta: −100 µm). (**K**) Calcium signals of 25 neurons randomly selected from each FOV. Color indicates FOV from which the neurons were selected. (**L**) Numbers of cells and FOVs (in brackets) recorded on 11 consecutive days. (**M**) SNR of all cells recorded on each day. Definition of each violin plot is the same as in Figure 2L. See also Figure S3.

While the size of the objective and the scanning angle of the MEMS scanner currently prevent additional increase of the FOV, we were able, with the second strategy, to further elevate the cell yield by imaging across multiple overlying planes with the new *z*-scanning function (Figure 3C-E, Video S2). Due to the relatively low *z*-resolution, as well as the typical 10-20 μmsize of neuron somata in layers 2 and 3 of the mouse cortex (Gilman et al., 2017), bleedthrough in the form of cells recorded in more than one plane (“repeated cells”) would be expected if the spacing between imaging planes is not large enough. In order to determine the minimum plane interval at which such bleedthrough could be largely avoided, within the constraints of the *z*-focusing module, we first quantified in VC the number of repeated cells in two-plane recordings with different plane intervals, using as criteria an *xy* center distance smaller than 15 µm, an anatomical overlap ratio larger than 50%, and a signal correlation larger than 0.50 (see STAR Methods; Figure S3A and S3B, miniscope type: MINI2P-F, frame rate: 40 Hz n= 2 mice, 6 FOVs and 5,602 cells). With an interval of 20 µm, a total of 22.9±5.3% of the cells (mean±S.D.) were identified as repeated cells. The fraction dropped to 4.4±1.8% at 40 µm, 0.1±0.2% at 60 µm, and 0% at 80 µm. Based on these data, we set the minimum plane interval to 40 µm in all VC recordings. A similar ratio of repeated cells was obtained with a 40 µm plane interval in MEC (Figure S3C, 5.0±4.3%). As a result of these investigations, multiplane imaging could be successfully applied across multiple brain areas (MEC, V1, higher-order visual cortices, CA1 and CA2; see Table S1, Part 1). With two imaging planes at each location, we were able to record 437±244 non-repeated neurons (maximum 737, in MEC). With four planes, we recorded 986±137 non-repeated neurons (maximum 1,103, in V1). Compared to single-FOV imaging (209±217 cells), the cell yield increased by 210 % and 440%, respectively (Figure 3C-E, Table S1, Part 1, and Video S2). The large scanning range, fast scanning speed and long-term *z*-scanning stability of MINI2P-F also enabled single-cell-resolution 3D functional imaging within a 420×420×160 µm^3^ volume, at a volume rate of 2 Hz, in one freely-moving mouse (Figure 3F, Figure 3G and Video S3, plane interval: 8 µm, 20 continuous planes). Individual cells were clearly identifiable at different locations of the volume (Figure 3F). High-SNR calcium signals could be extracted from each cell (Figure 3G). This first successful demonstration of large-population, high-resolution 3D functional imaging with 2P miniscopes opens the door to studies of 3D neuronal calcium dynamics in freely-moving animals (Video S3).

Finally, in the third strategy, we developed a FOV-stitching method to further increase cell yield by successive imaging of neighboring FOVs (Figure 3A, and Figure 3H-M). Instead of directly mounting the miniscope on the baseplate, we added an additional component (a “stitching adapter”) for connecting these two parts (Figure 2H). A series of interchangeable stitching adapters were designed to yield a certain amount of shift, in steps of 200 μm or 400 µm in *x* and *y*, between the center of the miniscopés FOV and the baseplate. With this procedure, the FOV of MINI2P could be shifted just by replacing the stitching adapter without removing the permanently fixed baseplate. A maximum of 5×5 400 μm shift steps were possible with the current baseplate design, covering more than 2×2 mm^2^ of imaging surface (Figure 3H). In an example two-plane recording using MINI2P-L from VC (Figure 3H-J, Video S4, Part I), we were able to extract in total 10,096 non-repeated layer 2/3 excitatory neurons via *Suite2p*, yielding an average of 404 cells per 500 × 500 μm^2^ FOV across the two planes (Figure 3J). After compensation of field distortion (Figure S3D and STAR Methods), highly reliable alignment of the FOVs was achieved (Video S4, Part II;). Precise alignment was achieved through a three-step procedure (Supplementary Document, Section 6): (1) primary alignment of FOVs by matching landmarks like blood vessels or labelled neurons in overlapping areas; (2) refinement of the alignment by monitoring correlation of neighboring FOVs; and (3) using manually identified, repeated cells in the overlap region to confirm the accuracy of the alignment (STAR Methods, and Video S4, Part II). If a wide-field image had been obtained as well (Figure 3H), the FOVs were also aligned to the wide-field image in step (1) to further increase the accuracy of alignment. Repeated neurons from adjacent FOVs were removed based on ratios of spatial overlap (common pixels) between neurons (Figure S3E and S3F). Due to the tight tolerance of baseplates and stitching adapters, the same cells could be re-visited for more than a week (Figure S3G).

With these three strategies for increasing cell yield, MINI2P could more than double the number of simultaneously recorded cells compared to previous 2P miniscope versions, and the number could be increased more than tenfold over successive recordings. In the following sections, we will demonstrate, for recordings in VC, MEC and hippocampal area CA1, how MINI2P can be used to monitor large-scale neural activity while mice explore large arenas over extended time periods.

### Using MINI2P for large-scale spatial tuning analysis in visual cortex

The visual cortices (VCs), including primary visual cortex (V1) and higher-level visual cortices (LM, AL, RL, AM, PM, etc.), cover a large area of the posterior cortical region in rodents (Hawrylycz et al., 2014; Wang et al., 2020). Growing evidence has shown that VC neurons are involved in spatial coding (Diamanti et al., 2021; Fiser et al., 2016; Haggerty and Ji, 2015; Ji and Wilson, 2007; Saleem et al., 2018), but studies of spatial coding that utilize 2P calcium imaging have been held back by the fact that animals were head-fixed and had to run in virtual environments (Diamanti et al., 2021; Fiser et al., 2016; Saleem et al., 2018). Here, as an illustration of how 2P miniscope imaging can be used to analyze the spatial tuning of large populations of neurons in VC during unrestrained behavior, we examined the data obtained in our stitched, two-plane recordings (10,096 neurons in VC, Figure 3H-M). FOVs were registered to the wide-field image. Regions belonging to primary and higher visual cortices (V1, LM, AL, RL, AM, and PM), were identified by retinotopic mapping (Zhuang et al., 2017) (Figure 3H, Video S4, Part II; Supplementary Document, Section 7; STAR Methods), and each recorded cell was assigned to one of these regions (Figure 4, Video S4, Part II). Before cells were analyzed for spatial tuning, key behavioral characteristics (running distance, arena coverage, speed and head direction distribution) were checked for each recorded session to ensure that data from all FOVs were stable and comparable.

**Figure 4.**
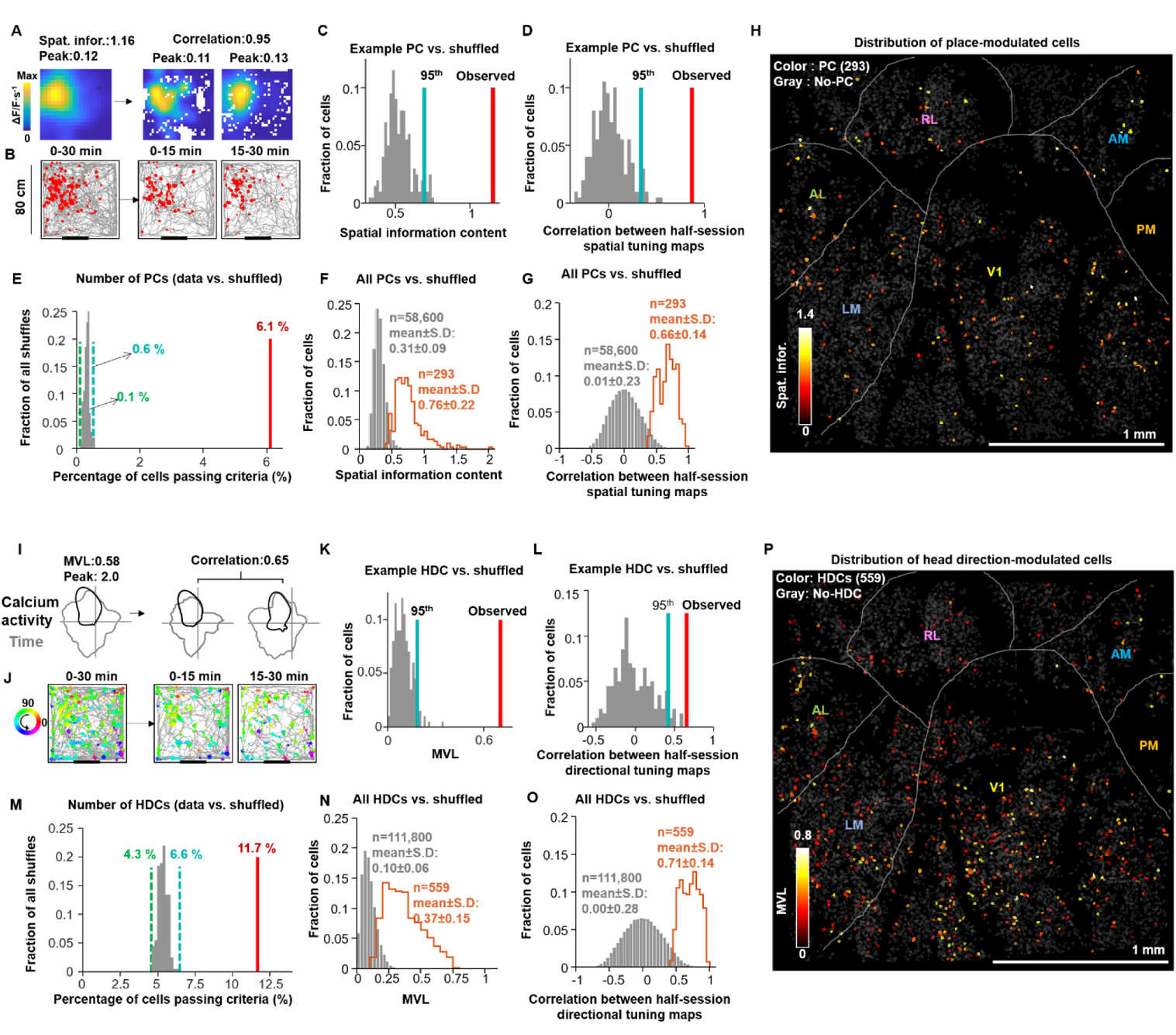
Spatial tuning features of 4,786 visual cortex neurons. (**A-H**) Quantitative analysis of place-modulated cells (PCs) in visual cortex (VC). (**A**-**D**) Example PC. (**A**) Spatial tuning maps showing localized calcium activity in an example PC (left: complete 30 min recording; middle: first half (0-15 min); right: second half (15-30 min)). Scale bar shows intensity of calcium activity (ΔF/F per second). Spatial bins: 2.5×2.5 cm^2^. The map was smoothed with a 3 cm Gaussian smoothing kernel. Unvisited bins were left white. (**B**) Animal trajectory (in gray) with calcium events superimposed (in red) for the whole 30 min recording (left), first half (middle) and second half (right). Same cell as in A. Size of red spots indicates the amplitude of deconvolved calcium events. (**C**) Spatial information content of the example PC (red bar) is higher than the 95th percentile (blue bar) of the distribution of the shuffled data (grey bars), n = 200 iterations. (**D**) Half-session spatial tuning map correlation values (first vs. second half) of the example PC (red bar) is higher than the 95th percentile value (blue bar) of the data from 200 shuffled iterations (grey bars). (**E**) The percentage of PCs in the recorded data (red bar, number of cells n=293, 6.1 %) is higher than the percentage range of PCs in the shuffled data for all cells. Dashed lines show min (0.1%, green) and max (0.6%, blue) percentage in the shuffled data. (**F**) PCs had higher spatial information content (n=293, mean±S.D, 0.76±0.22) than in the shuffled data. Orange curve indicates distribution of values for spatial information content in recorded data from cells that satisfied criteria for PCs. Gray bars indicate distribution of spatial information for the shuffled data. (**G**) PC spatial tuning maps had higher half-session spatial correlation values than in the shuffled distribution. Orange curve indicates distribution of correlation values from PCs that satisfied criteria in the recorded data. Gray bars indicate distribution in shuffled data. (**H**) Map showing distribution of PCs (dots) across the recorded matrix of FOVs. Bright yellow indicates high and dark red indicates low spatial information content (color bar). Gray indicates cells that did not meet place cell cutoff criteria (No-PCs). (**I-P**) Quantitative analysis of head direction-modulated cells (HDCs) in VC. (**I**-**L**) Example HDC. (**I**) Directional tuning map showing localized calcium activity as a function of head direction (HD) (left: complete 30 min recording; middle: first half; right: second half). Faint gray lines show head direction occupancy across each recording block. Black lines show the intensity of calcium activity as a function of head direction in polar coordinates. Each HDC’s tuning curve was normalized by the peak calcium activity, shown on the top of the plots, and each occupancy curve was normalized by peak occupancy time. (**J**) Animal trajectory (in gray) with calcium events superimposed for the example HDC during the whole 30 min recording (left), first half (middle) and second half (right). Events are color-coded by head direction, size of spots indicates the amplitude of deconvolved calcium events. (**K**) MVL of the example HDC (red bar) is higher than the 95th percentile value (blue bar) of a distribution of shuffled data from the same cell. Colors as in C and D. (**L**) Half-session directional correlations for the example HDC is higher than the 95th percentile value of shuffled data from the same cell. Colors as in K. (**M**) Percentage of HDCs in the recorded data (red bar, n=559, 11.7 %) is higher than the percentage range of HDCs in shuffled data from all of the HDCs; dashed lines show min (green) to max (blue): 4.3 % and 6.6 %, respectively. (**N**) HDCs (n=559) had higher mean vector length (MVL) than in the shuffled data. Orange curve indicates distribution of MVL values from all HDC in the recorded data. Gray bars indicate the distribution of MVLs in all permutations of the shuffled data. (**O**) HDCs had higher half-session directional correlations than in the shuffled data). Orange curve indicates distribution of correlation values from all HDCs in the recorded data. Gray bars indicate distribution of values in the shuffled data. (**P**) Map showing distribution of HDCs (dots) across the recorded matrix of FOVs. Bright yellow indicates high and dark red indicates low mean vector length (MVL) of the directional tuning curve. Gray indicates cells that did not pass criteria for HDCs (No-HDC). See also Figure S4.

We examined the spatial tuning of 4,786 non-repeated neurons in this VC sample with SNRs above 3 and event counts (non-zero incidences of deconvolved calcium activity) of 100 or more (Figure 4). Place-modulated cells (PCs) and head direction-modulated cells (HDCs) were identified by a combination of criteria. PCs were defined as cells that passed combined criteria of (1) spatial information (SI), (2) spatial stability (correlation between successive half-session spatial tuning maps), (3) size of firing fields, and (4) peak calcium activity in the firing fields. For each criterion, a cutoff was defined by the 95^th^ percentile of the distribution of values in 200 shuffled iterations of the event times of the same cell (STAR Methods). A total of 293 cells, or 6.1%, of the 4,786 cells passed all four criteria (see Figure 4A-D for an example cell; spatial tuning maps of 100 additional PCs are shown in Figure S4B). This number was compared to the count of cells that passed all four criteria in a shuffling control in which the deconvolved events of the same 293 cells were shuffled repeatedly in time (Figure S4A; STAR Methods). The percentage of PCs per iteration of shuffling ranged from 0.1 to 0.6%, which is well below the percentage in the recorded data (Figure 4E). The 293 PCs that passed criteria in the recorded data had sharply focused and stable spatial tuning fields, as expressed by high spatial information content (0.76±0.22; mean ± S.D.; Figure 4F) and high spatial correlation values (0.66±0.14; Figure 4G). There was also a notable number of HDCs in the VC data (n=559 or 11.7%, Figure 4I-P, an example is shown in Figure 4I-L; directional tuning maps of 100 additional HDCs are shown in Figure S4C). HDCs were defined as cells that passed criteria of (1) directional tuning (mean vector length, MVL) and (2) directional stability (correlation between half-session directional tuning curves). The percentage of HDCs significantly exceeded chance levels determined by shuffling of the 559 HDCs following the same procedure as for PCs. The range of cells per iteration of shuffling was from 4.3 to 6.6 %, again well below that of the recorded data (Figure 4M). The HDCs expressed sharp and stable directional tuning, assessed by high values for MVL (0.37±0.15, Figure 4N) and high directional correlation (0.71±0.14, Figure 4O).

The two cell types were distributed across all VC regions, with no observable clustering within or between regions in the single mouse that we imaged (Figure 4H and 4P). Overall, we have shown with this example recording that with MINI2P it is possible to record more than 10,000 neurons at high resolution across a matrix of 5×5 two-plane FOVs in VC. Thus, MINI2P enables anatomically and functionally precise quantification of spatial tuning properties in densely active cortical populations during unconstrained exploratory behavior.

### Using MINI2P to analyze spatial tuning in MEC and PAS

Among the most difficult cells to characterize with calcium imaging in freely moving mice are the grid cells of the MEC (Hafting et al., 2005) not only because of the vertical orientation of the cell layer, next to the transverse sinus, but also because the mouse must cover the spatial environment homogeneously for the repeating pattern to be extracted. Samples of tens to hundreds of spike events may be required to detect the hexagonal structure of the firing fields. Here we demonstrate that hundreds of identifiable grid cells could be recorded simultaneously from a single freely-moving mouse with MINI2P (Figure 5, Figure S5, Video S5 to Video S7). The recording was performed using MINI2P-L through a 1.3×1.3×1.6 mm^3^ prism over a total area of 0.9×0.9 mm^2^, with stitching of five neighboring 500×500 μm^2^ FOVs (4 FOVs with a 400 μm shift step, 1 FOV with a 200 μm shift step, Figure 5A and 5C). Imaging was centered on the dorsomedial MEC but included parts of the adjacent PAS (Table S1, Part 4). By pre-injecting into hippocampal regions CA3 and dentate gyrus (DG) a retrogradely transported virus (Tervo et al., 2016) that expressed a red fluorophore (retro-tdTomato) (Figure 5B; see also Obenhaus et al, accompanying manuscript), we were able to identify MEC from red-channel expression in the FOVs, due to the strong projections to CA3 and DG from MEC but not from PAS (Witter et al., 2017) (Figure 5D).

**Figure 5.**
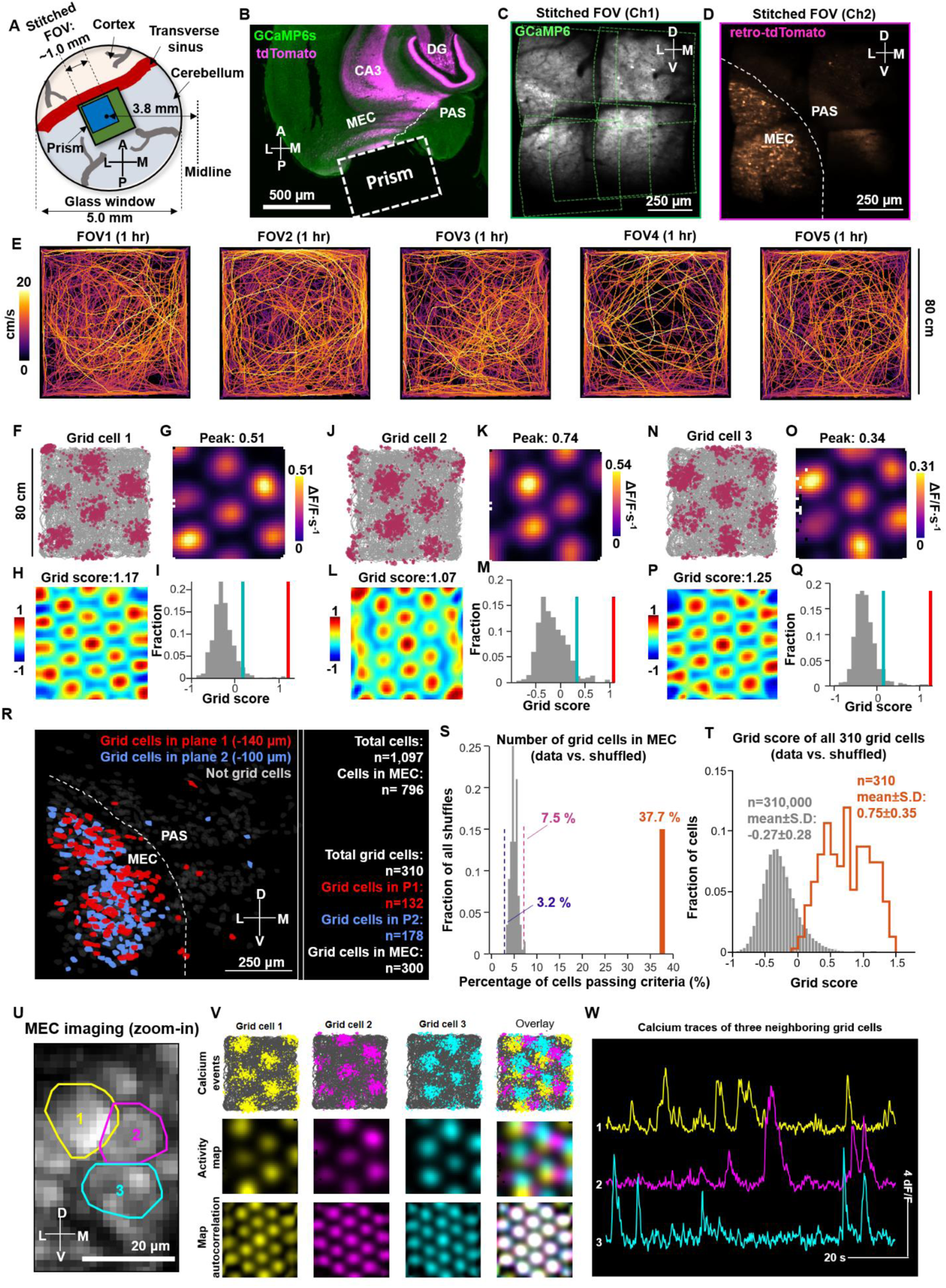
MINI2P recordings in MEC: 310 grid cells among a total of 1,097 neurons. (**A**) Illustration of stitched FOV positions relative to prism and cover glass. (**B**) Horizontal brain section with fluorescence showing strong expression of tdTomato (purple channel) in CA3, DG and layer 2/3 of MEC, indicating ML position of the prism with reference to the medial border of MEC. PAS parasubiculum. (**C-D**) Average projections from 5 stitched FOVs. For each FOV, cells were recorded in two planes, at 40 µm separation. The projections of plane 2 (−100 µm) are shown. (**C**) GCaMP6 channel (Ch1). Green stippled lines show the borders of each FOV. (**D**) TdTomato channel (Ch2). MEC and PAS were distinguished based on tdTomato expression. (**E**) Animal trajectories from five recordings (1 hour each). Color indicates momentary running speed (color bar). (**F-I**) Example grid cell. (**F**) Animal trajectory (in gray) with calcium events superimposed (in purple). Size of purple spots indicates the amplitude of deconvolved calcium events. (**G**) Spatial tuning map showing localized calcium activity. Scale bar to the right indicates activity in ΔF/F per second. Spatial bins 2.5 cm. (**H**) Color-coded autocorrelation of spatial tuning map from panel G. Scale bar to the left (correlation from −1 to 1). The cell’s grid score is indicated above each autocorrelation map. (**I**) Grid scores compared to shuffled data from the same cell where the calcium events were shifted individually and randomly in time across the whole session. Red bar indicates grid score in recorded data. Blue bar indicates 95th percentile of grid scores in 1,000 iterations of shuffling in the same cell (gray bars). (**J-Q**) Two additional example grid cells (J and N as in F, K and O as in G, L and P as in H, M and Q as in I). (**R**) Distribution of grid cells across the MEC and PAS (dark red: grid cells at −140 µm from the surface, dark blue: grid cells at −100 µm; light gray cells did not pass criteria for grid cells). Stippled line indicates border between MEC and PAS. (**S**) The percentage of grid cells within MEC (orange bar, n=300, 37.7%) significantly exceeds the percentage of grid cells in 1,000 iterations of shuffling of deconvolved events from the same 300 grid cells. Dashed lines show min (purple) and max (pink) percentage in the shuffled data from all 300 grid cells (min 3.2 % and max 7.5 %, respectively). (**T**) Grid cells had higher grid scores than in the shuffled data. Orange curve indicates distribution of grid scores from grid cells that satisfied the cutoff criteria in the recorded data. Gray bars indicate distribution in shuffled data. (**U-W**) Examples of neighboring grid cells in the same plane with similar grid spacing and grid orientation but mixed grid phase. (**U**) Zoomed-in image from one MEC recording shows three neighboring grid cells. The colored polygons show the outline of each grid cell. The color code in U is maintained in V and W. The image shows a projection of 5,000 frames from the motion-corrected time-lapse recording. (**V**) Calcium events superimposed on the trajectory (top), spatial tuning map (middle), and autocorrelation of the spatial tuning map. Columns show individual grid cells (1st to 3rd), with their deconvolved calcium events overlaid in the 4th column. Brightness of spatial tuning maps and autocorrelation indicates intensity scaled to the maximum. Note overlap of 6 inner fields in autocorrelation map indicating that the four grid cells share orientation and scale. Note also the separation of clusters (fields) of different cells in the spatial tuning maps. (**W**) Calcium traces for each of the three grid cells. The uncorrelated signal of the calcium transients is consistent with minimal contamination from adjacent grid cells. See also Figure S5, and Table S1 part 4 and part 5.

During imaging, the animal explored the arena uniformly (five 1 hour recordings; Figure 5E). The total number of recorded cells, after removing repeated cells, was 1,097 (5 FOVs; Figure 5R, Table S1, Part 5). The imaging speed (stack rate: 7.5 Hz) was sufficiently fast to capture the spatial grid pattern (with running speeds often exceeding 20 cm/s, slow imaging could have blurred spatial firing patterns). Using shuffled data for reference, we identified 310 grid cells in total (three examples in Figure 5F-Q, all 310 grid cells are shown in Figure S5O-Q), 300 of which were identified within the tdTomato-expressing region (MEC), near its dorsomedial tip. A few scattered grid cells were recorded in the adjacent PAS (Figure 5R), in agreement with earlier electrophysiological recordings (Boccara et al., 2010). Grid cells were identified in similar numbers in both imaging planes (132 grid cells at −140 µm depth and 178 separate grid cells at −100 µm, 4 examples in Video S5; depth measured from the surface of MEC). The recording depths correspond to layer 2 and superficial layer 3 of MEC. The percentage of grid cells in MEC was 37.7 %, a lot more than the 3.2 to 7.5% range obtained in shuffled versions of data from the same 300 MEC grid cells (Figure 5S, 1,000 iterations of shuffling, same procedure as for VC data above). Recorded grid cells had sharply tuned hexagonally patterned firing fields with grid scores (means±S.D.) of 0.75±0.35 (Figure 5T). Grid spacing ranged from 30 cm near the dorsal end of MEC to about 60 cm near the ventral end of the FOVs (Figure S5A; for further examples see Obenhaus et al, accompanying manuscript) and grid axes were generally aligned with the walls of the recording box (Figure S5B), in agreement with previous tetrode data (Brun et al., 2008; Fyhn et al., 2004; Stensola et al., 2012; Stensola et al., 2015). We also detected, in each plane, a fraction of conjunctive grid × head direction cells, i.e. cells that passed criteria for both grid and head direction-modulated cells (two examples in Figure S5C-N). Finally, the resolution of MINI2P allowed us to demonstrate differences in grid phases among neighboring grid cells, with little visible contamination from adjacent cells or neuropil (Figure 5U-W, and Video S6). We observed, as expected from early tetrode recordings (Hafting et al., 2005), that neighboring grid cells had mixed grid phases (Figure 5V, top and middle) but shared the same spacing and orientation (Figure 5V, bottom).

The large-scale recordings of hundreds of grid cells, in conjunction with other spatially modulated cells at high spatial resolution, allowed us to analyze in-depth how these cell types are organized anatomically. In the accompanying research article, we used this unprecedented opportunity to determine the topographic arrangement of spatially modulated neurons in MEC and adjacent PAS (Obenhaus et al, accompanying manuscript).

### By improving coverage of the environment, MINI2P enhances detection of grid cells

Given the abundance of sharply tuned grid cells that we were able to record with MINI2P in MEC, we next sought to examine the impact that weight and cable flexibility had on the identification of grid cells. We tested this in an example animal (Figure 6, Figure S6, and Table S1, Part 6). First, we recorded cells in the open field for 30 min (block 1) with the 0.7-mm TFB (3g-t, as in Figure 1). The animal was then put back into its home cage for 10 min, after which we added a 2g dummy weight on the microscope (bringing the total weight to ~5.0 g, similar to 5g-T in Figure 1), replaced the thin cable with a 1.5-mm supple fiber bundle (SFB, as for 5g-T in Figure 1), and then recorded for another 30 min (block 2). A total of 129 grid cells were identified in the combined data (block 1 + block 2; 1,000 permutations, 95^th^ percentile cutoff, imaging parameters as in Figure 5; see also Table S1, Part 6). We then compared the grid score of each cell obtained from data in block 1 and block 2 separately to the shuffling cutoff obtained in the combined data (block 1 + block 2). The results showed a clear change from the first to the second block not only in the animal’s free-foraging behavior (Figure 6A) but also in the ability to detect grid cells (Figure 6B and 6C). Grid scores of all grid cells identified in the combined recording decreased significantly when after changing weight and cable, from 0.44 ± 0.43 in the first block to −0.14 ± 0.37 in the second block (Figure 6B, paired t-test, n=129, p<0.001), a change that likely reflected incomplete coverage of positions and directions during the second block. While 52% of the grid cells (67 of 129) passed the criterion during the first 30 min block, only 6% (8 of 129) passed it during the second block (Figure 6C). The change was not merely caused by the length of the recording session as a similar effect was not apparent on two consecutive 30 min 3g-t recording blocks (Figure S6, grid scores and the percentages of grid cells were stable at 0.51±0.44 and 0.52±0.49, and at 74 % (77 of 104) and 73 % (76 of 104), for the first and second block, respectively). From these observations we conclude that the light weight of the miniscope and the flexibility of the cable assembly is critical for accurate detection of grid cells in freely moving animals.

**Figure 6.**
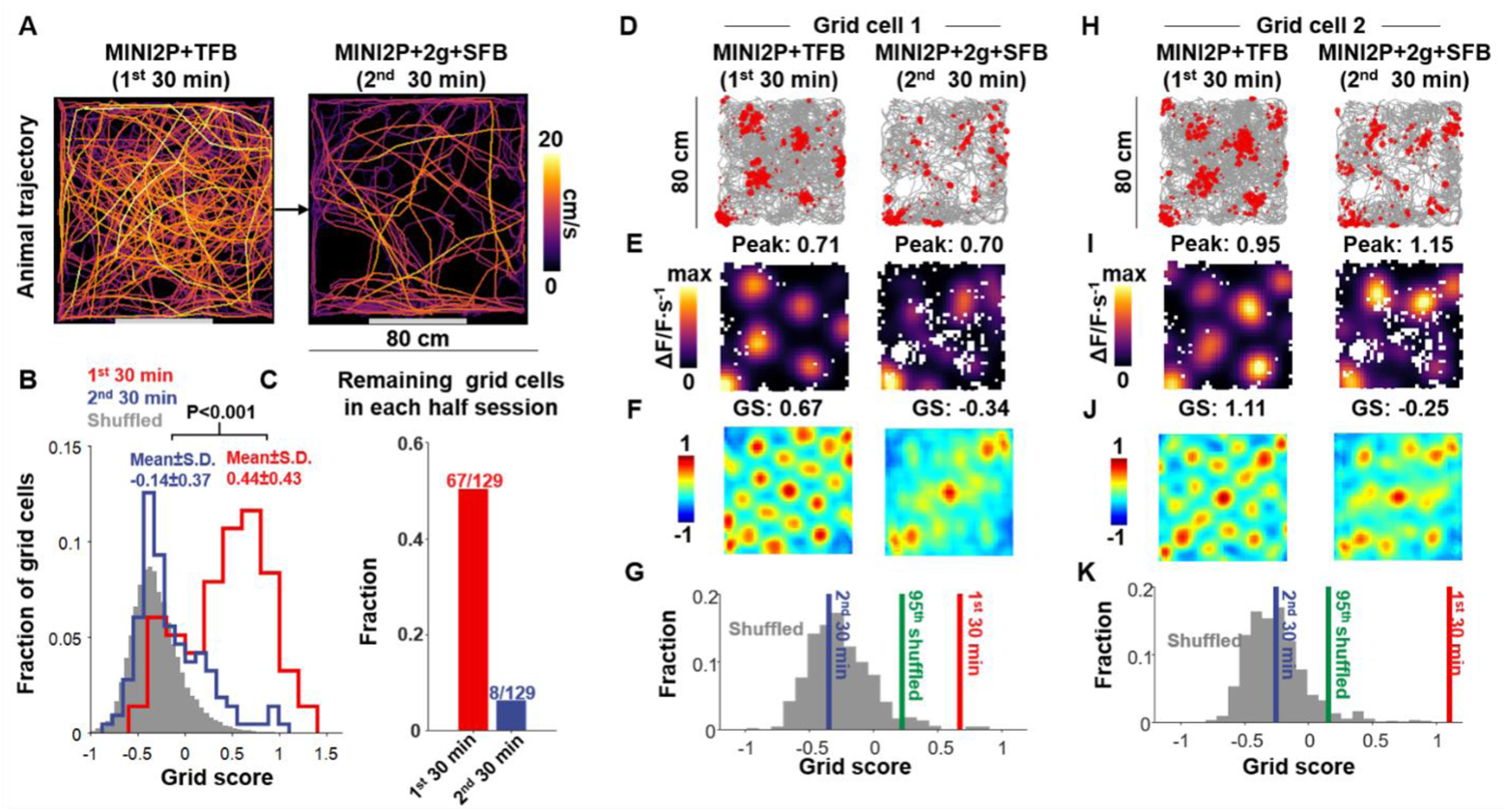
Flexible cable connection and low weight of MINI2P substantially improve detection of grid cells. (**A**) Top: trajectory of a mouse with 3g MINI2P microscope and the tapered fiber bundle (TFB) during 30 min of recording compared to the trajectory of the same mouse carrying a 5g miniscope (MINI2P with 2g additional weight block) and the supple fiber bundle (SFB) on an immediately succeeding 30 min trial. Color indicates momentary running speed (color bar). Note reduced running and coverage with added weight and SFB cable, as in Figure 1. (**B**) Distribution of grid scores (a measure of grid symmetry) during the first (red) and second (blue) half-session for grid cells identified from the full session (two blocks of 30 min). The distribution of shuffled data is shown for comparison (gray). Grid scores decreased significantly towards shuffled levels with the 5g microscope and the SFB (P value indicates result of paired t-test). (**C**) Ratio of grid cells from the full session that passed the threshold for grid cells in the first (red) and second (blue) half-session. (**D** and **H**) Calcium event distributions for two example grid cells in the two conditions. Calcium events were superimposed (in red) on the animal’s trajectory (in gray). Size of red spots indicate the amplitude of deconvolved of calcium events. (**E** and **I**) Spatial tuning maps. Scale bar to the left. Spatial bins 2.5×2.5 cm2. (**F** and **J**) Color-coded autocorrelation of the spatial tuning maps from panel E and I. Scale bar to the left. The cell’s grid score is indicated above each autocorrelation map. (**G** and **K**) Grid scores in the first half-session (red) and second half-session (blue) for the two example cells, compared to shuffled data for the same cell obtained from the full session. Green bar indicates 95th percentile of grid scores for 1,000 iterations of shuffling of the same data (gray bars). See also Figure S6 and Table S1, Part 6.

### Using MINI2P to analyze place cells in hippocampal area CA1

For a final test of MINI2P’s versatility for imaging in different brain regions, we recorded another mouse from hippocampal area CA1, the brain region where 1P miniscope imaging has been applied most successfully (Rubin et al., 2019; Sheintuch et al., 2017; Ziv et al., 2013). In this area, 1P imaging has been possible because neurons are distributed in a thin, sparsely active cell layer. Similar to previous studies employing 1P imaging, we used a GRIN lens to image CA1 activity (Figure 2R, Figure 7A). We successfully recorded 340 non-repeating cells across two imaging planes (miniscope type: MINI2P-L, plane interval 60 µm), 254 of which had SNRs above 3 and event counts over 100 (Figure 7B). Among these cells, we identified a total of 122 place-modulated cells (PCs, Figure 7B), using the same criteria as in VC (two examples in Figure 7C-F). PCs could have a single place field (Figure 7C and D) or occasionally more (Figure 7E and F). The percentage of PCs in the recorded data (Figure 7G, red bar, n=122, 48.0 %) was significantly higher than the range of percentages (0.0 to 6.7 %) meeting criteria for PCs in shuffled versions of data from the same PCs (1,000 iterations, shuffling procedure as for VC and MEC). PCs in CA1 had higher spatial information content (0.97±0.33; mean ± S.D.) and higher half-session spatial correlation (0.83±0.09) than in shuffled data from the same cells (Figure 7H and 7I).

**Figure 7.**
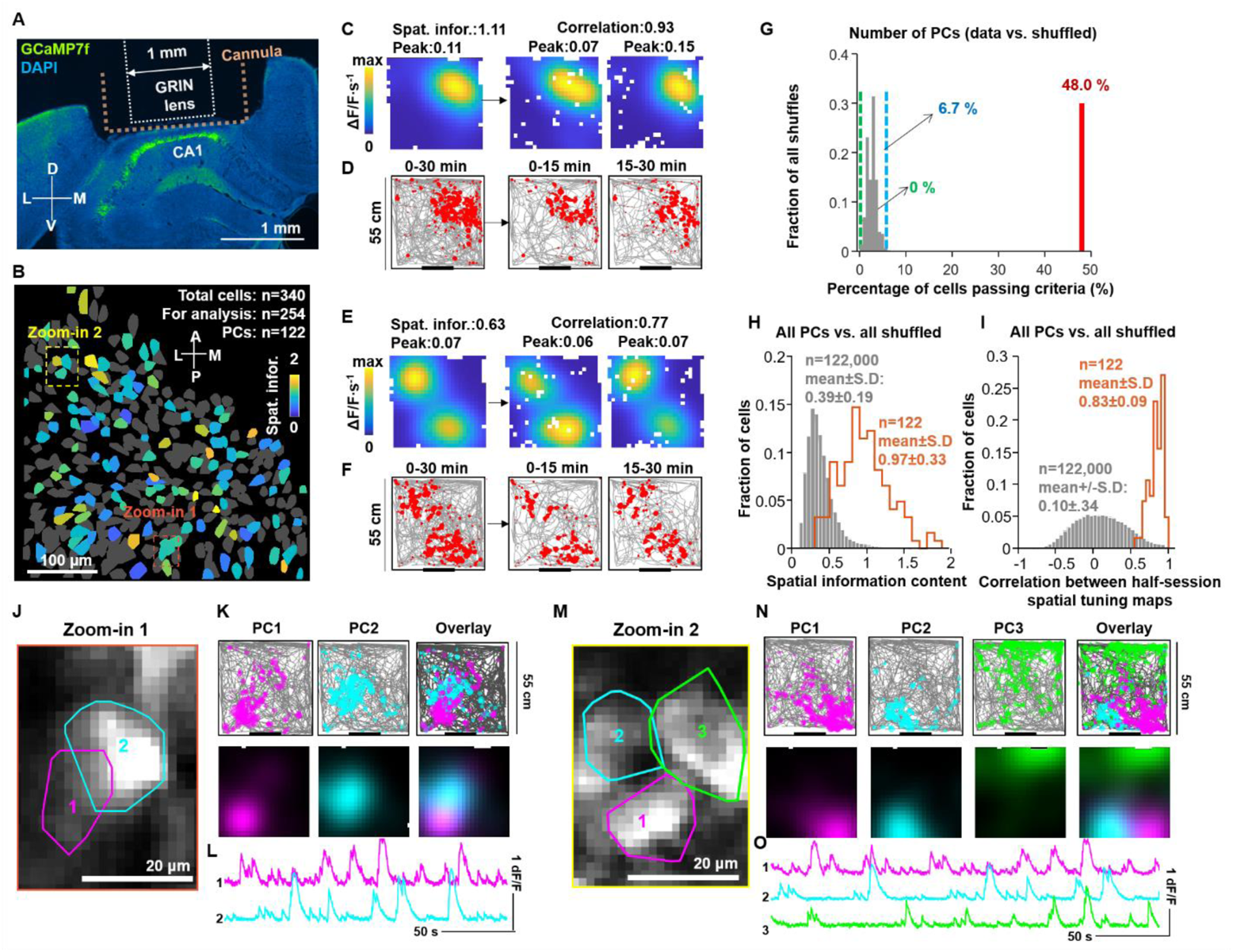
Spatial tuning of 254 cells recorded with MINI2P in hippocampal area CA1. (**A**) Coronal brain section with fluorescence showing strong expression of GCaMP7f (light green) in the pyramidal layer of CA1. The tissue above CA1 was aspirated and a cannula was implanted to keep space for subsequent GRIN lens insertion. (**B**) Map showing distribution of place-modulated cells (PCs) across a single FOV. Bright yellow indicates high and dark blue indicates low spatial information content. Gray indicates cells that did not meet place-cell cutoff criteria. Two dashed boxes (red and yellow) correspond to the zoomed-in regions in J and M. (**C**-**F**) Two example PCs in CA1. (**C** and **D**) PC with a single place field. (**C**) Spatial tuning maps showing localized calcium activity (left: complete 30 min recording; middle: first half (0-15 min); right: second half (15-30 min)). Scale bar shows intensity of calcium activity (ΔF/F per second). Spatial bins: 2.5×2.5 cm^2^. Map was smoothed with a 3 cm Gaussian smoothing kernel. Unvisited bins were left white. (**D**) Animal trajectory (in gray) with calcium events superimposed (in red) in the recording box for the whole 30 min recording (left), first half (middle) and second half (right). Size of red spots indicate deconvolved amplitude of calcium events. (**E** and **F**) PC with two place fields. E as in C, F as in D. (**G**) The percentage of PCs in the recorded data (red bar, n=122, 48.0 %) is higher than the percentage of cells meeting criteria for PCs in the shuffled data for all cells (1,000 iterations). Dashed lines show min (green) and max (blue) percentage in shuffled data (min 0 % and max 6.7 %, respectively). Criteria for place cells were the same as for place-modulated cells in VC (Figure 4) except for the use of a different threshold of min peak activity for the fields. (**H**) PCs had higher spatial information content than in the shuffled data. Orange curve indicates distribution of values for spatial information content in recorded data from cells that satisfied criteria for PCs. Gray bars indicate distribution of spatial information for the shuffled data. (**I**) Spatial tuning maps of PCs showed higher correlation between first and second half of a session than in the shuffled data (all cells). Orange curve indicates distribution of correlation values from PCs that satisfied criteria in the recorded data. Gray bars indicate distribution in the shuffled data. (**J-O**) Examples of neighboring PCs in the same plane with different place field locations. (**J**) Zoomed-in image from the CA1 recording shows two neighboring PCs. Colored polygons indicate outline of each PC. Color code is maintained in K and L. The image is a projection of 5,000 frames from the motion-corrected time-lapse recording. (**K**) Top panel shows calcium events superimposed on the trajectory whereas bottom shows brightness-coded spatial tuning maps scaled to maximum (individual cells in 1^st^ and 2^nd^ column; overlap in 3rd column). Note separation of the place fields of the two neighboring PCs. (**L**) Calcium traces for each PC in K. The uncorrelated signal of the calcium transients indicates minimal contamination from adjacent PCs. (**M-O**) Similar to (J to L) but for three neighboring PCs. M as in J. N as in K, and O as in L.

The total number of cells recorded from CA1 with MINI2P was comparable to numbers reported in studies using 1P imaging, with larger FOVs for this brain area (Rubin et al., 2019; Ziv et al., 2013). However, 2P imaging avoids the signal blurring of 1P imaging, caused by tissue scattering, which could result in signal spillover between neighboring active cells. In the present study, we show, with 2P miniscopes, that neighboring PCs can be distinguished reliably (Figure 7J-O), with little evidence of signal crosstalk between adjacent neurons (Figure 7L and 7O). The data demonstrate how MINI2P can be used for precise studies of the complete microcircuit organization of CA1.

### MINI2P is fully open-source

MINI2P can be constructed in biology labs with basic optics and electronics experience. Key components such as interchangeable objectives, μTlens, hollow-core photonic crystal fiber, and TFB, are commercially available (see Supplementary Document, Section 11, for a full list). Ordering numbers of key components, 3D models and drawings of the customized components, building instructions (Figure S7J), customized recording software, data-processing, as well as spatial tuning analysis code are available (see Key Resources Table). The entire imaging setup can be integrated into a mobile cart smaller than 1 m^3^ and can be installed in a standard recording room without an air conditioner or dust filter (Figure S7A-I). MINI2P is fully controlled by open-source 2P acquisition software (ScanImage) (Pologruto et al., 2003).

## Discussion

We have designed a 2P miniscope (MINI2P) that enables the study of neural activity, at high sampling rates and with high spatial resolution, in thousands of individually identifiable cells in brain regions with different architectures and locations, while mice move freely in the environment. By retaining the optical sectioning capability afforded by hardware-demanding 2P excitation, MINI2P can be used for high-signal-to-noise calcium imaging, irrespective of somatic packing density, density of neuropil, or neuronal activity level, in brain regions where imaging is often precluded with current 1P miniscopes. Furthermore, the ability to simultaneously record from a second structural channel using a red-shifted fluorophore allows identification of projection-labeled or genetically-identified subpopulations within the recorded cell sample (Video S7). With the present reduction in headpiece weight and the increase in flexibility caused by tapering of the collection fiber bundle, MINI2P allows animals to behave with the same level of flexibility as with 1P miniscopes, despite the added 2P hardware. Mice can carry MINI2P microscopes on their head for half an hour or more without interruptions, while they roam freely in open environments at speeds, directions and rotation rates comparable to those of untethered animals without miniscopes. Successive recording sessions can be carried out without bleaching of imaged neurons or changes in the quality of signal or the animal’s behavior. Taken together, these features of MINI2P allow for high-resolution imaging of unprecedented numbers of neurons in widespread brain circuits during spatially dispersed behaviors.

MINI2P outperforms previous versions of 2P miniscopes not only in total imaging volume and number of cells that can be monitored but also in that neural activity can be imaged for days or weeks in tasks with naturalistic exploratory or goal-directed behavior. With previous versions of 2P or 3P miniscopes, which weigh 4.2 g (Zong et al., 2021) or 5.0 g (Klioutchnikov et al., 2020) and are connected via fiber bundles of 1.5-mm in diameter, behavior remains comparable to that of unimplanted control mice only as long as mice occupy small housing cages (20×30 cm^2^) (Zong et al., 2021). Even in these environments, the mice spent the majority of time in the corners of the box, regardless of whether they were with miniscopes or not, and total running distances were very short. By systematically comparing the behavior of mice carrying different types of miniscopes, we found in the present study that while running is obviously affected by the weight of the headpiece, the strongest impediments on behavior were related to stiffness and diameter of the connection cable assembly, which is dominated by the optical recording fiber bundle. Inflexible cables substantially compromise turning behavior and coverage of the environment and thereby strongly reduce the ability to detect spatially patterned cells such as grid cells. With the new light-weight design of MINI2P and the tapering of the optical fiber bundle, these differences in behavior could no longer be observed compared to control mice, not even after more than 30 min of running in an enlarged environment. To make full use of the added potential of 2P compared 1P imaging, namely the ability to record from multiple, adjacent planes without bleedthrough, miniature *z* scanning mechanisms with high thermal stability are required. While previous 2P miniscope versions (Ozbay et al., 2018; Piyawattanametha et al., 2009; Zong et al., 2021) did integrate *z* scanning modules, their applicability to long-term multiplane imaging in freely-moving animals was strongly compromised by slow focusing speed (Piyawattanametha et al., 2009), low resolution (Ozbay et al., 2018), or heavy weight and strong, thermally induced *z* drift over even short recordings (Zong et al., 2021). Here, by applying the μTlens, a novel *z*-scanning device with decreased weight and increased focusing speed, we extend multiplane imaging further to demonstrate, for the first-time, high-resolution 3D functional imaging in freely behaving animals with stability across tens of minutes, paving the way for studies of 3D neuronal calcium dynamics in freely-moving animals.

We illustrated the applicability of MINI2P in three brain regions with vastly different anatomical locations requiring various optical access methods: glass covers were used to reach horizontally oriented regions of visual cortex (VC), prisms for vertically oriented regions such as MEC, and GRIN lenses for deep regions such as CA1. By imaging near-simultaneously from multiple parallel planes, the cell yield could be extended to over a thousand in a 420×420×120 µm^3^ volume in VC, and by using a novel procedure for successive imaging of neighboring FOVs, the yield could be increased beyond 10,000 neurons. Recordings in MEC could reach more than 1,000 cells and in CA1 several hundreds. These cell yields are an order of magnitude larger than those of previous 2P miniscopes, which reached maxima of a few hundred in easily accessible surface regions of the brain.

While using MINI2P for a proof-of-principle large-scale recording in VC, we were able to demonstrate the presence of multiple spatially tuned cell classes in areas of the brain that are often not thought to play a primary role in position coding. In V1 as well as neighboring visual regions, we identified cells with properties similar to those of the place cells and head direction cells of the hippocampus and entorhinal cortices. Head direction-modulated cells were abundant in all subregions of VC, in agreement with earlier electrophysiological recordings (Guitchounts et al., 2020; Velez-Fort et al., 2018). A low but significant number of cells that were tuned to location exhibited tuning properties similar to those of hippocampal place cells. These place-modulated neurons fired reliably whenever the mouse ran through a discrete location in the environment. The observation of extrahippocampal place-tuned neurons in the cortex is consistent with electrophysiology recordings from VC (Haggerty and Ji, 2015; Saleem et al., 2018) and the adjacent retrosplenial cortex (Mao et al., 2017; Mao et al., 2018) as well as 2P imaging studies in VC of head-fixed mice running on a linear track in virtual environments (Diamanti et al., 2021; Fiser et al., 2016; Saleem et al., 2018). The present findings provide a large-scale mapping of place modulation in VC cells in one animal and add the observation that these cells are scattered across visual regions and different imaging depths. These findings show that with MINI2P, responses of hundreds to thousands of spatially and directionally tuned neurons, that are phenotypically similar to cells recorded in hippocampal and parahippocampal regions, can be explored in VC during unrestrained behavior in the real world. Taken together, these observations raise the possibility that VC plays an active role in spatial coding during navigation, in support of previous studies in head-fixed animals (Diamanti et al., 2021; Fiser et al., 2016; Saleem et al., 2018).

We also demonstrated for the first time the feasibility of 2P miniscope imaging for large-scale recordings through a prism implanted along the elongated, dorsoventral surface of MEC. At the border between MEC and PAS, we were able to record from more than a thousand neurons, among which we identified a cluster of several hundred grid cells. Many of the recorded grid cells were labelled by tdTomato expression, confirming that grid cells project to the CA3 and dentate gyrus regions of the hippocampus (Video S7) (Zhang et al., 2013). The sharp tuning of firing fields in the presence of running speeds of over 20 cm/s suggests that sampling is fast enough to avoid distortion of the cellś firing fields. Within the anatomical cluster of grid cells, the most dorsally located cells had smaller grid spacing, in the order of 30-40 cm, whereas more ventrally located cells exhibited wider spacing, in the order of 40-60 cm (see also Obenhaus et al, accompanying manuscript). Neighboring grid cells largely had distinct grid phases, with non-overlapping grid field locations, as expected from earlier data (Gu et al., 2018; Hafting et al., 2005). The clear separation of grid phases confirms the lack of bleedthrough of signals between neighboring neurons. The ability to record from over a hundred densely packed and cleanly isolated cells in MEC and neighboring PAS simultaneously and during unrestrained behavior offers the unique opportunity to study the functional organization of cells in these higher association cortices. The analysis of the topographic arrangement of all spatially modulated cell types in these regions is out of reach for both electrophysiological methods (because of insufficient tissue coverage and lack of anatomical precision) and for 2P benchtop recordings (because of the necessary head fixation). In the accompanying research article (Obenhaus et al, accompanying manuscript) we use a variant of MINI2P to systematically determine the anatomical arrangement of all known spatially tuned cell types in MEC and adjacent PAS, including grid, object vector, HD, and border cells.

We finally showed that MINI2P can be used in combination with GRIN lenses for population imaging of deep structures such as CA1, where we were able to record over one hundred active place cells in CA1. Signal bleedthrough was also not observed in CA1, mirroring the analysis of neighboring cells in MEC and pointing to 2P miniscopes as a useful tool for precise studies of the complete assembly of neurons in local circuits of place cells. In their current form, for CA1 specifically, cell yields with 2P miniscopes do not outcompete cell numbers obtained with 1P miniscopes (Rubin et al., 2019; Ziv et al., 2013), but 2P miniscopes add the possibility to more confidently separate the activity of densely active neighboring cells. In the future, improved optical access methods (Velasco and Levene, 2014), that circumvent the use of aberration-prone GRIN lenses, will expand the accessible surface area, which, in combination with FOV-stitching techniques introduced here, will allow for further increase of cell yield in CA1.

We expect the development of head-mounted 2P miniscopes to continue during the years to come. While the MINI2P microscope has now been miniaturized sufficiently for use in a broad range of behavioral tasks that afford unrestrained behavior, miniature optics such as metalenses will lead to further decreases in weight and size of the microscope (Arbabi et al., 2018), and miniaturized photodetectors will allow in-situ signal detection so that the connection cable can be yet more flexible (Bortfeldt et al., 2020). Moreover, while the current miniscope enables recording from more than a thousand neurons at a time, pulse-multiplexing technology (Cheng et al., 2011) and adaptive optics (Ji, 2017; Wang et al., 2014) will undoubtedly increase the cell yield further. New types of hollow-core photonic crystal fibers will allow multi-wavelength excitation, increasing the options for choices of indicator (Amrani et al., 2021), and 2P miniscope imaging will be fused with optogenetic stimulation technology (Emiliani et al., 2015; Marshel et al., 2019), enabling large-scale mapping of connectivity between functionally characterized neurons not only in virtual-reality setups but also during unrestrained behavior. By open-sourcing protocols and software for assembly and use of the new miniscope, we hope that a growing number of investigators will be involved in the further development of large-scale high-resolution imaging in freely behaving rodents.

## Acknowledgments

We thank A.M. Amundsgård, K. Haugen, K.J. Jenssen, E. Kråkvik, and H. Waade for technical assistance, I. Ulsaker-Janke, R.I. Jacobsen, T. Rose, J. Sugar, A.Z. Vollan, T. Waaga, and M.M. Bolton for discussion of analyses and data, T. Slettmoen and F. Donato for suggestions on MEC implantation. R.I. Jacobsen and T. Rose for suggestions on visual cortex imaging, M.P. Witter for suggestions on anatomy and help with identification of recording positions, R. N. Raveendran and the Viral Vector Core Facility of the Kavli Institute for Systems Neuroscience (NTNU) for viral reagents, H.-K. Yddal for support with development of the Tlens, and S. Eggen and I. Ulsaker-Janke for help with animal surgery. The work was supported by a Synergy Grant from the European Research Council to E.I.M. (‘KILONEURONS’, Grant Agreement N° 951319), a Research Council of Norway (RCN) FRIPRO grant to E.I.M. (grant number 286225), a Research Council of Norway (RCN) FRIPRO grant to M.-B.M. (grant number 300394), an RCN Centre of Excellence scheme grant and an RCN National Infrastructure grant to E.I.M. and M.-B.M. (Centre of Neural Computation, grant number 223262; NORBRAIN, grant number 295721), a direct contribution to M.-B.M. and E.I.M. from the Ministry of Education and Research of Norway, the Kavli Foundation, and funding from the European Union’s Horizon 2020 Research and Innovation Programme under the Marie Skłodowska-Curie grant agreement to W.Z. (ANAT-MEC NO 842006).

## Author contributions

Conceptualization: W.Z., E.I.M, M.-B.M, H.A.O.

Supervision: E.I.M, M.-B.M

System design: W.Z.

Hardware development: W.Z, N.J.

System control: W.Z., H.A.O.

Software development: W.Z.

Data collection: W.Z, H.E, N.J., E.R.S.

AAV injection methodology: W.Z, H.A.O., E.R.S.

Behavior experiment: H.E., W.Z.

VC and MEC surgery and recording: W.Z

MEC surgery protocol: H.A.O., W.Z.,

CA1 surgery and recording: E.R.S.

Pilot studies on previous versions of microscope and data processing: H.A.O., W.Z.

Statistical Analysis: W.Z., H.E., E.I.M.

Figures: W.Z, H.E., N.J., M.R.J.

Videos: W.Z.

Materials lists and protocol: W.Z. and M.R.J.

Writing – original draft: W.Z., E.I.M.

Writing – review & editing: E.I.M., M.-B.M., W.Z., H.A.O.

Discussion and comments on paper draft: all authors

---

**Figure S1.**
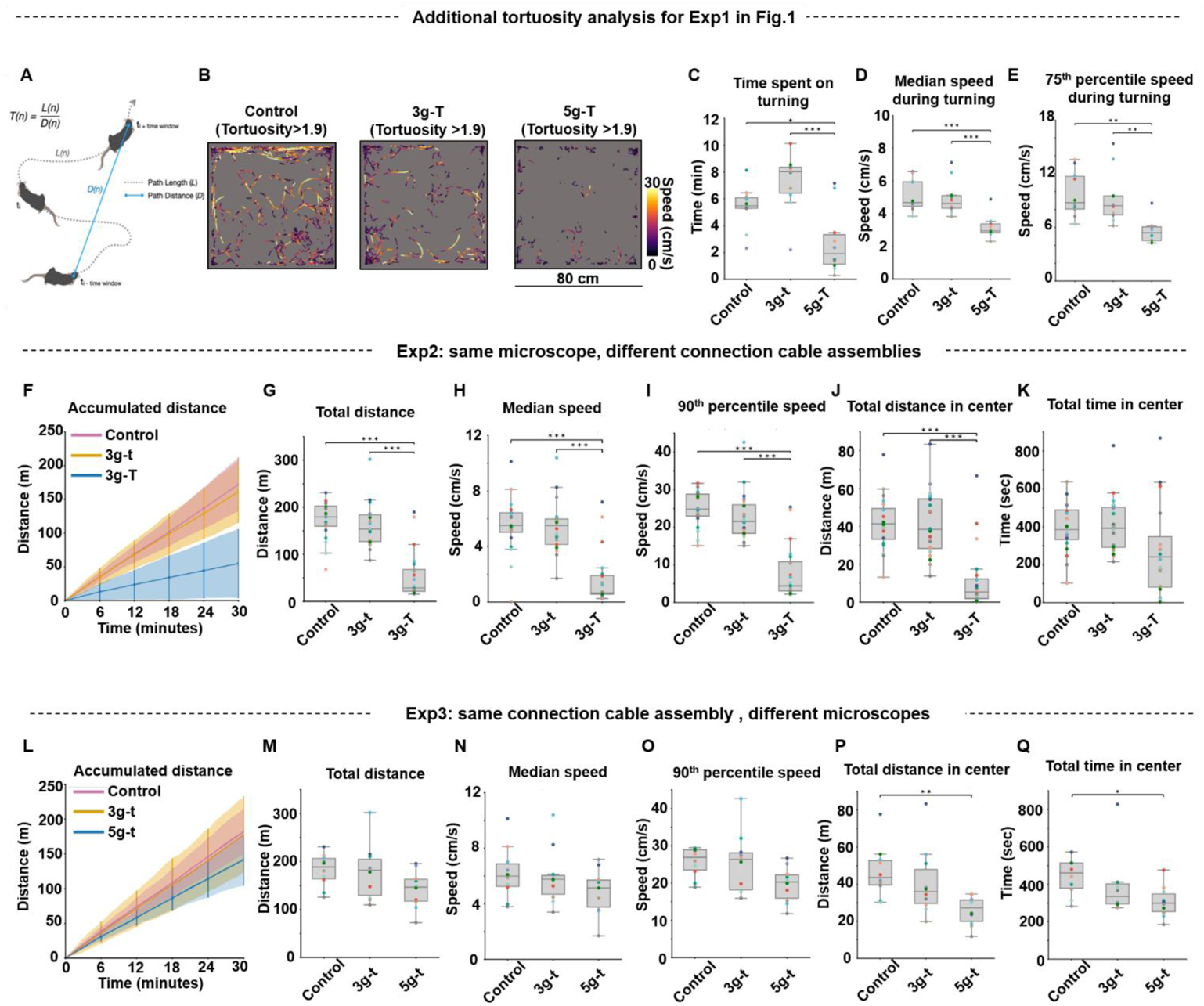
Further analysis of behavioral impact of cable thickness and microscope weight. Related to Figure 1. (**A**) Schematic of tortuosity analysis. Total path length within a time segment was compared to the Euclidean distance between the start and end point of the window. (**B-E**) The time spent on turning was defined as the total time of frames with tortuosity above the 75th percentile tortuosity value computed from all frames in control recordings with no miniscope and no cable (a value of 1.9). This time was reduced in the 5g-T group but not in the 3g-t group (n=10 in each condition; Friedman test on data from all three experimental conditions: χ2 =6.2, p =0.045; post-hoc Tukey *t* test for 5g-T vs. 3g-t, p<0.001, 5g-T vs. control: 0.025; 3g-t vs. control: 0.136). Running speed during turning moments was lower in the 5g-T experiment than in 3g-t and control (n=10 in each condition; Friedman test: χ2 > 12.6, p<0.0018; post-hoc Tukey *t* test for 5g-T vs each of the other groups: p<0.018; 3g-t vs. control: p>0.493). (**B**) Representative turning trajectories in three experiments in which frames with tortuosity lower than the 75th percentile over all values for tortuosity were removed. Color scale shows value of momentary speed during turning. (**C**) Total time spent on turning (tortuosity >75^th^ percentile) across the three experiments. (**D**-**E**) Median speed (**D**) and 75th percentile speed (**E**) during turning (tortuosity >75^th^ percentile). (**F-K**) Isolating the effect of fiber thickness (flexibility) on behavior (n = 20 animals). Control: no dummy or cable, 3g-t: 3g microscope and thin connection cable assembly, and 3g-T: 3g microscope and thick connection cable assembly. Friedman test on data from all three experimental conditions, total running distance: χ2 = 30.1, p<0.0001; median running speed: χ2 = 24.3, p <0.0001; 90th percentile running speed: χ2 = 31.6, p <0.0001; total running distance in center: χ2 = 24.4, p <0.0001; total running time in center: χ2 = 6.3, p = 0.043; post-hoc Tukey tests for 3g-T vs control: p < 0.001 (total running distance, median running speed, 90th percentile running speed, and total running distance in center) or p = 0.130 (total running time in center); post-hoc Tukey tests for 3g-T vs 3g-t: p < 0.001 (total running distance, median running speed, 90th percentile running speed, and total running distance in center) and p = 0.085 (total running time in center); post-hoc Tukey tests for 3g-t vs control: p = 0.726 (total running distance); p >0.90 (median running speed), p = 0.490 (90th percentile running speed), p >0.90 (total running distance in center), p >0.90 (total running time in center). Differences in running time in the center did not reach statistical significance because several mice with thick cables stopped and rested in the middle of the box (where the strain of the cable was smaller than in the periphery). (**F**) as in Figure 1B, (**G**) as in Figure 1C, (**H**) as in Figure 1D, (**I**) as in Figure 1E, (**J**) as in Figure 1F, and (**K**) as in Figure 1G. (**L-Q**) Isolating the effect of microscope weight on behavior (n=10). Control: no dummy or cable, 3g-t: 3g miniscope and thin connection cable assembly, and 5g-t: 5g miniscope and thin connection cable assembly. Friedman test on data from all three experimental conditions, total running distance: χ2 = 9.8, p = 0.0074; median running speed: χ2 = 6.2, p = 0.045; 90th percentile running speed: χ2 = 11.4, p = 0.003; total running distance in center: χ2 = 7.2, p = 0.027; total running time in center: χ2 = 5.6, p = 0.061; post-hoc Tukey tests for 5g-t vs control, p = 0.128 (total running distance); p = 0.269 (median running speed); p = 0.078 (90th percentile running speed)), p = 0.007 (total running distance in center), and p =0.048 (total running time in center); Tukey tests for 5g-t vs 3g-t: p = 0.187 (total running distance); p =0.421 (median running speed); p =0.107 (90th percentile running speed), p = 0.067 (total running distance in center), and p = 0.319 (total running time in center); Tukey tests for 3g-t vs control: p >0.90 (total running distance); p>0.90 (median running speed); p = 0.618 (90th percentile running speed); p = 0.618 (total running distance in center); and p=0.563 (total running time in center);. (**L**) as in Figure 1B, (**M**) as in Figure 1C, (**N**) as in Figure 1D, (**O**) as in Figure 1E, (**P**) as in Figure 1F, and (**Q**) as in Figure 1G. Definitions in box plots are the same as in Figure 1.

**Figure S2.**
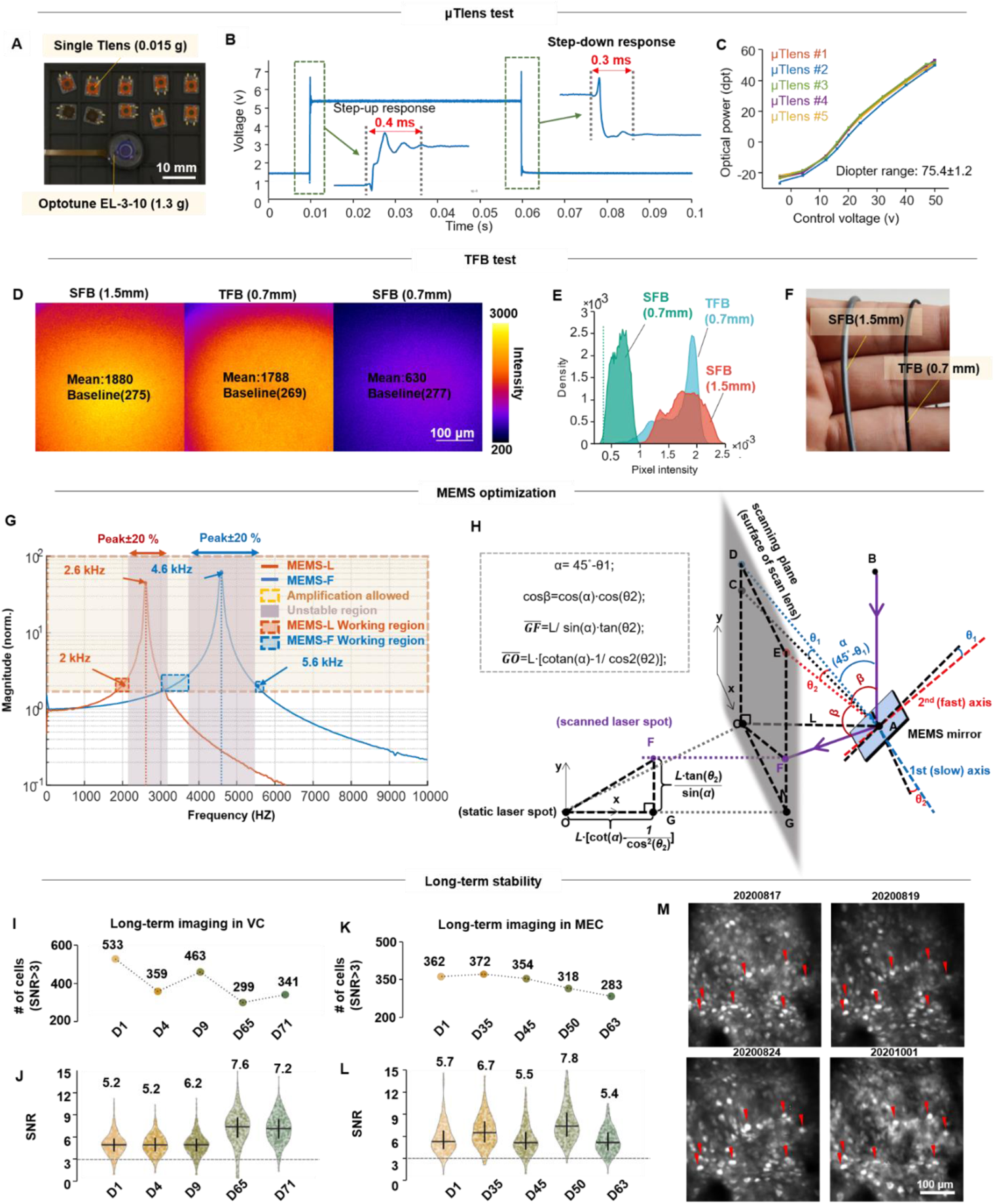
Supplementary features and performance of MINI2P. Related to Figure 2. (**A-C**) µTlens test. (**A**) Multiple single µTlenses (top) and an Optotune EL-3-10 (ETL, used in the previous generation of miniscope (Zong, et al., 2021), bottom). (**B**) Representative step response of the quartet µTlens. (**C**) Optical power (diopter) vs. control voltage for five quartet µTlens samples. (**D-F**) TFB test. (**D-E**) Detection efficiency comparison between 1.5 mm SFB, 0.7 mm TFB and 0.7 mm SFB. (**D**) Imaging of a uniform fluorescent slide (FSK2, Thorlabs, NJ, USA) with the 1.5 mm SFB (left), 0.7 mm TFB (middle) and 0.7 mm SFB (right). Pixel intensity is color-coded; bright yellow indicates higher intensity; dark purple indicates low intensity. Laser power, PMT sensitivity and pixel dwell time were identical in the three recordings. (**E**) Histograms showing pixel intensity of images in (**D**). Dashed line indicates baseline of the image (average intensity of the image without exposure). (**F**) Picture of SFB (left) and TFB (right). (**G** and **H**) MEMS optimization. (**G**) Working frequency analysis for two types of MEMS scanners, one with a larger scanning angle but slower speed (MEMS-L), and the other with smaller scanning angle but faster speed (MEMS-F). (**H**) Geometry and mathematics of MEMS scanning in MINI2P (see also STAR Methods for details). (**I-M**) Imaging stability of MINI2P. (**I** and **J**) Hundreds of neurons can be recorded for more than 2 months in the same FOV of VC in the mouse shown in Figure 2I to 2L. (**I**) Number of cells across days. Recording day (D1, D4, etc.) is color-coded (from D1 to D71). (**J**) SNRs are maintained throughout the recording period. The outline of the violin plot shows probability density smoothed by a kernel density estimator (bandwidth: 10% of the data range). Interquartile range and median are indicated with vertical and horizontal lines, respectively. (**K** and **L**) Hundreds of neurons can be recorded for more than 2 months in the same FOV of MEC in the mouse shown in Figure 2M-Q(D1-D63). (**K**) Number of cells across recording days. Recording day is color-coded (from D1 to D63). (**L**) SNRs are maintained throughout the recording period. (**M**) The same neurons can be followed across 1.5 months of imaging in the same FOV (from data shown in I, no stitching, red arrows point to 7 example cells appearing in all FOVs across recording days).

**Figure S3.**
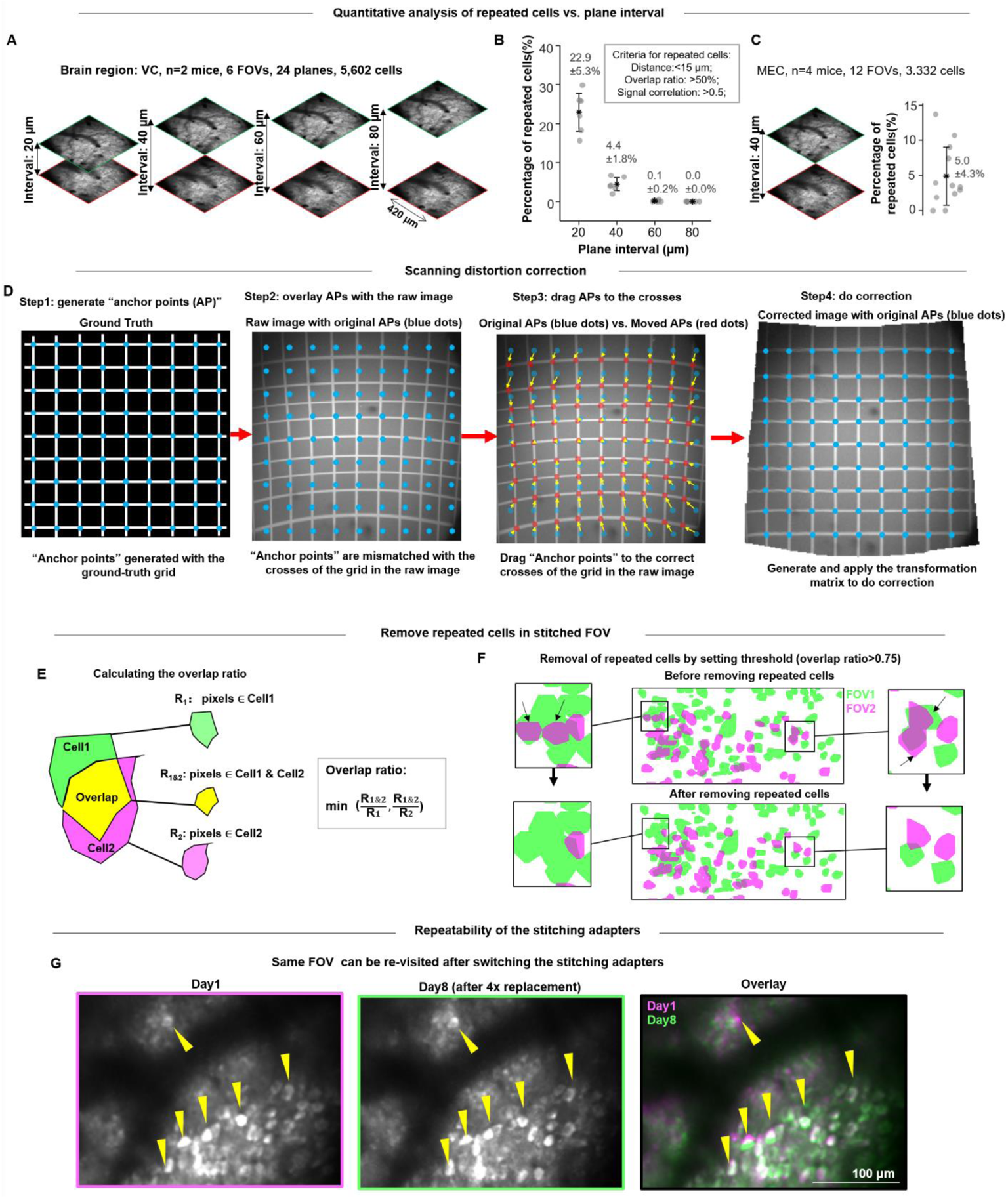
Multiplane imaging, distortion correction and FOV stitching. Related to Figure 3. (**A-C**) Quantitative analysis of the fraction of repeated cells as a function of plane interval in VC and MEC recordings. See also Table S1 part 2 and part 3. (**A**) Double-plane calcium imaging in VC with different plane intervals (from 0 µm to 80 µm in 20 µm steps). Recording duration: 5 min. Frame rate: 20 Hz. Stack rate: 10 Hz. 2P miniscope: MINI2P-F, Imaging area: VC. Indicator: GCaMP6s. All imaging planes were within 200 µm from the cortical surface. (**B**) Relationship between plane interval and the percentage of repeated neurons. See also Table S1, Part 2. (**C**) 2-plane imaging in MEC with plane interval fixed at 40 µm (left). The fraction of repeated cells is comparable to imaging with a 40-μm interval in VC (right). 2P miniscope: MINI2P-L, See also Table S1, Part 3. (**D**) Step-by-step illustration of the distortion correction procedure. (**E** and **F**) Removal of repeated cells after accurate FOV alignment. (**E**) Definition of overlap ratio for a cell pair. (**F**) Repeated cells were successfully removed by thresholding the data at an overlap ratio of 0.75. (**G**) The same FOV can be re-visited after switching the stitching adapters. In this example data, the same neurons (yellow arrow tips) could be revisited after 8 days and after replacing the stitching adapter a total of 4 times.

**Figure S4.**
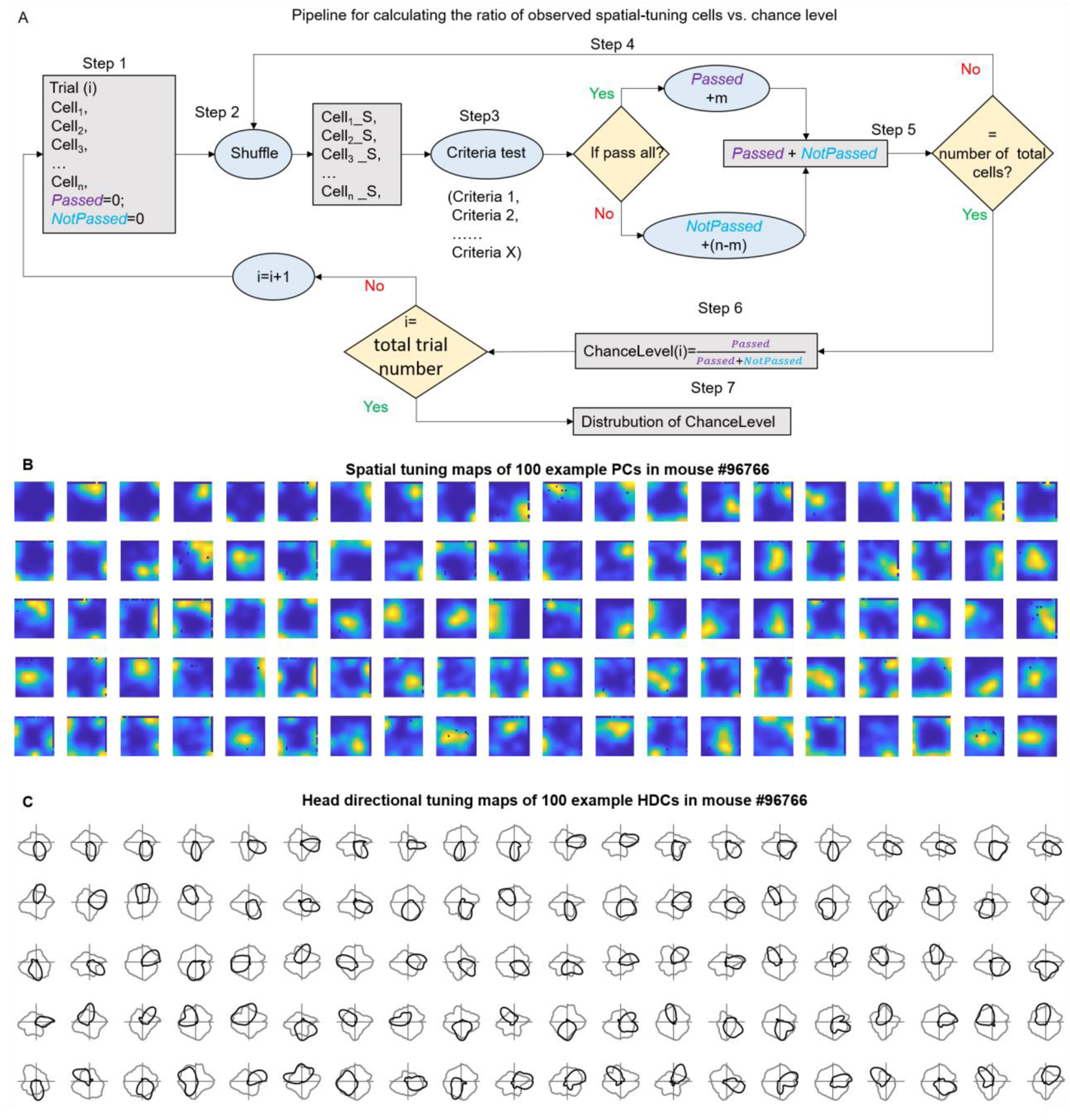
Supplementary for spatial tuning analysis in VC. Related to Figure 4. (**A**) Pipeline for calculating the ratio of observed spatially-tuned cells with reference to statistical chance level. In each trial, Step1: Select recorded spatially tuned cells that passed all criteria (Cell1, Cell2,…., Celln), and initiate the counts of Passed and NotPassed; n is the total number of spatially tuned cells for testing. Step2: Shuffle all spatially tuned cells, get Cell1_S, Cell2_S,…., Celln_S). Step3: Check if these shuffled cells passed all criteria. Step4: If m shuffled cells passed all criteria, Passed=Passed+m; and NotPassed = NotPassed+(n-m). Step5: If the total number of Passed+NotPassed cells reaches the same number as the total number of neurons (spatially tuned and not spatially tuned cells) or more, stop shuffling, calculate the ChanceLevel in trial i: Passed/(Passed + NotPassed). Step6: Repeat Step1 through Step6 until the trial number reaches the desired number of trials, calculate the range of ratios of shuffled cells passing criteria, and compare to the observed ratio (red bar in step7). (**B**) Spatial tuning maps of n=100 example place-modulated cells (PC) for mouse #96766. PCs are ranked by spatial information (SI) and PCs with higher SI are on top. The colormap definition is the same as in Figure 4A. (**C**) Head directional tuning maps of n=100 example head direction-modulated cells (HDCs) for mouse #96766. HDCs are ranked by MVL and HDCs with higher MVL are on top.

**Figure S5.**
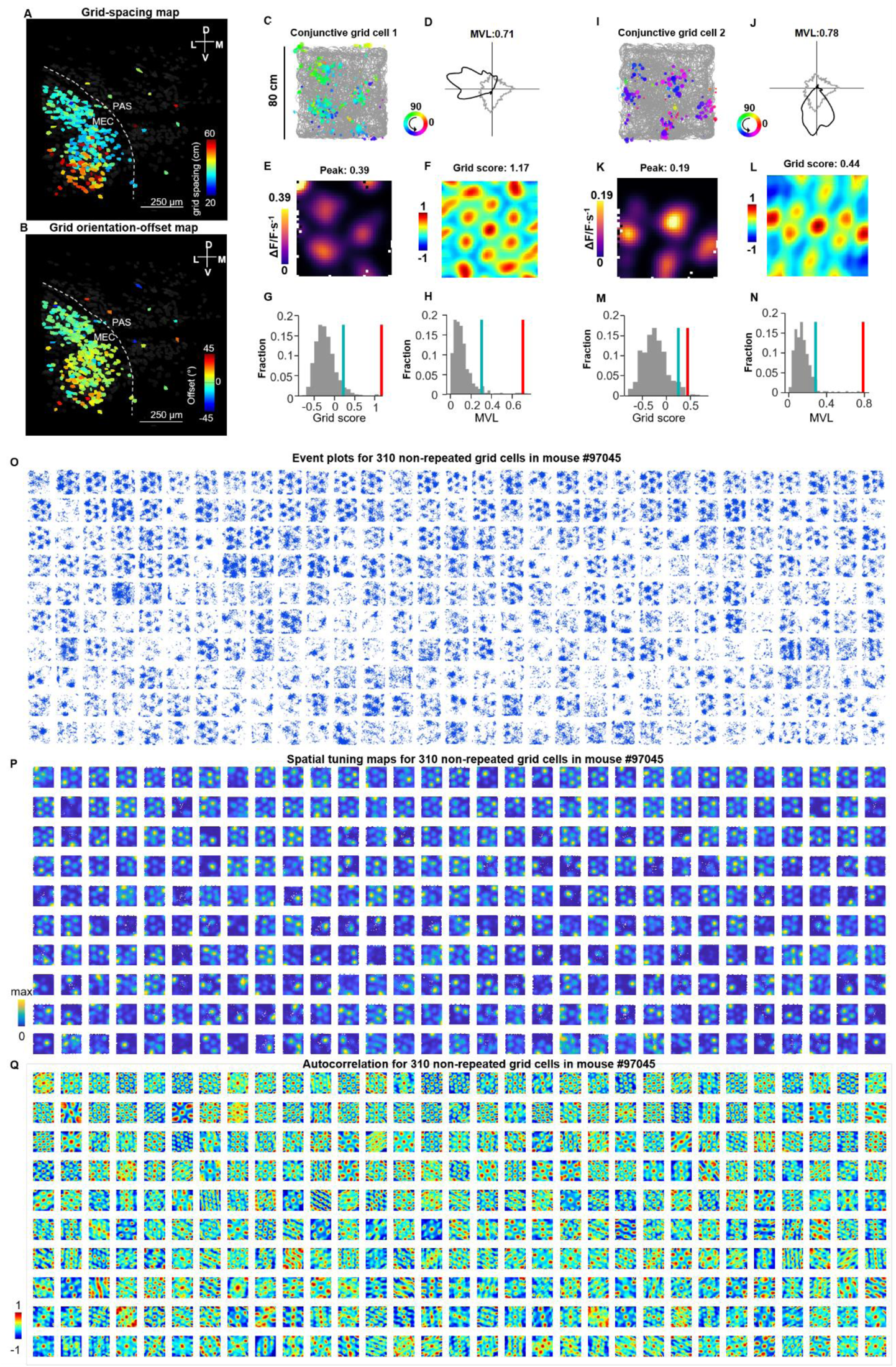
Features of grid cells in mouse #97045. Related to Figure 5. (**A**) Grid spacing map (same data as in Figure 5). Grid cells were color-coded by grid spacing (distance between the center field and the mean of six inner fields on the autocorrelation map). (**B**) Grid orientation map (same data as in Figure 5). Grid orientation was expressed as the arithmetic angular mean of the three main axes of the inner fields (Axes 1-3) on autocorrelation maps. The grid orientation offset is the smallest angle of Axes 1-3 to the north-south wall of the box. See STAR Methods for definition of orientation offset. (**C-H**) Example conjunctive grid × head direction-modulated cell. (**C**) Animal trajectory (in gray) with calcium events superimposed (in color). Events are color-coded by head direction, size of spots indicates the amplitude of deconvolved calcium events. (**D**) Directional tuning map showing localized calcium activity as a function of head direction (HD). Faint gray lines show head direction occupancy across each recording block. Black lines show the intensity of calcium activity as a function of head direction in polar coordinates. Tuning curve was normalized by the peak calcium activity, and each occupancy curve was normalized by peak occupancy time (**E**) Spatial tuning map showing localized calcium activity. Scale bar to the right indicates activity in ΔF/F per second. Spatial bins 2.5 cm. (**F**) Color-coded autocorrelation of spatial tuning map from panel E. Scale bar to the left (correlation from −1 to 1). The cell’s grid score is indicated above each autocorrelation map. (**G**) Grid scores compared to shuffled data from the same cell where the calcium events were shifted individually and randomly in time across the whole session. Red bar indicates grid score in recorded data. Blue bar indicates 95th percentile of grid scores in 1,000 shuffling iterations (gray bars). (**H**) MVL of the example (red bar) is higher than the 95th percentile value (blue bar) in shuffled data from the same cell (1,000 iterations). Colors as in G. (**I-N**) Another example conjunctive grid × head direction-modulated cell (I as in C, J as in D, K as in E, L as in F, M as in G, and N as in H). (**O-Q**) Grid cells in an MEC example mouse (Mouse #97045). Grid cells were plotted in (**O**)-(**Q**) in sequence of grid scores. Upper rows show grid cells with the highest grid scores. (**O**) Plots showing spatial location of calcium events (without animal’s trajectory) for 310 grid cells (same cells, as the total grid cells in Figure 5). Size of spots indicates amplitude of individual calcium event. (**P**) Spatial tuning maps for the same cells as in O. Color code in each map is scaled to peak calcium activity. (**Q**) Color-coded autocorrelation of the spatial tuning maps for the same cells as in (**O**) and (**P**).

**Figure S6.**
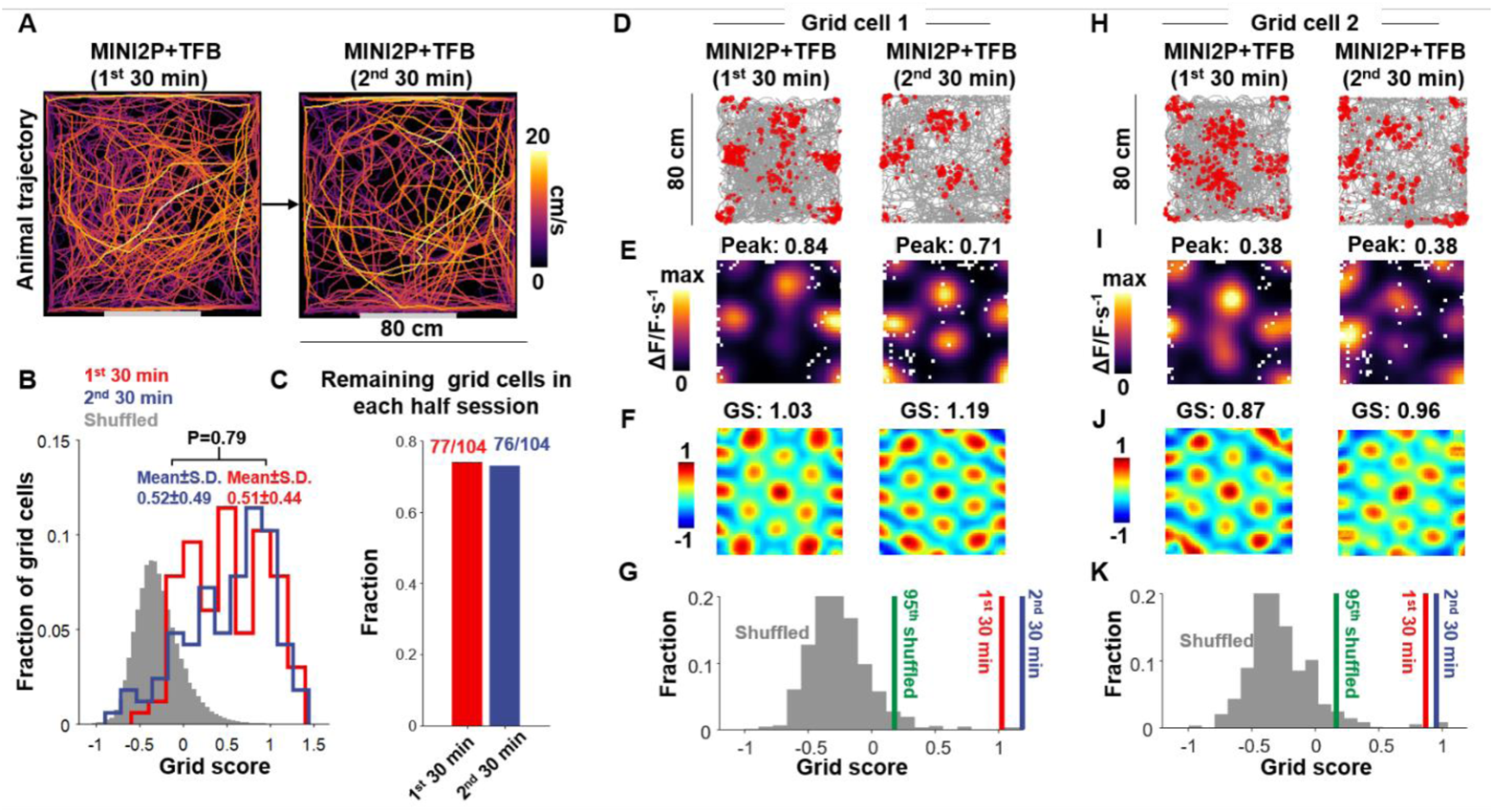
Same analysis as in Figure 6 but on two consecutive 30 min control recordings with the 3.0 g MINI2P microscope and the TFB (no change in weight or cable). Related to Figure 6. See also Table S1 Part 6.

**Figure S7.**
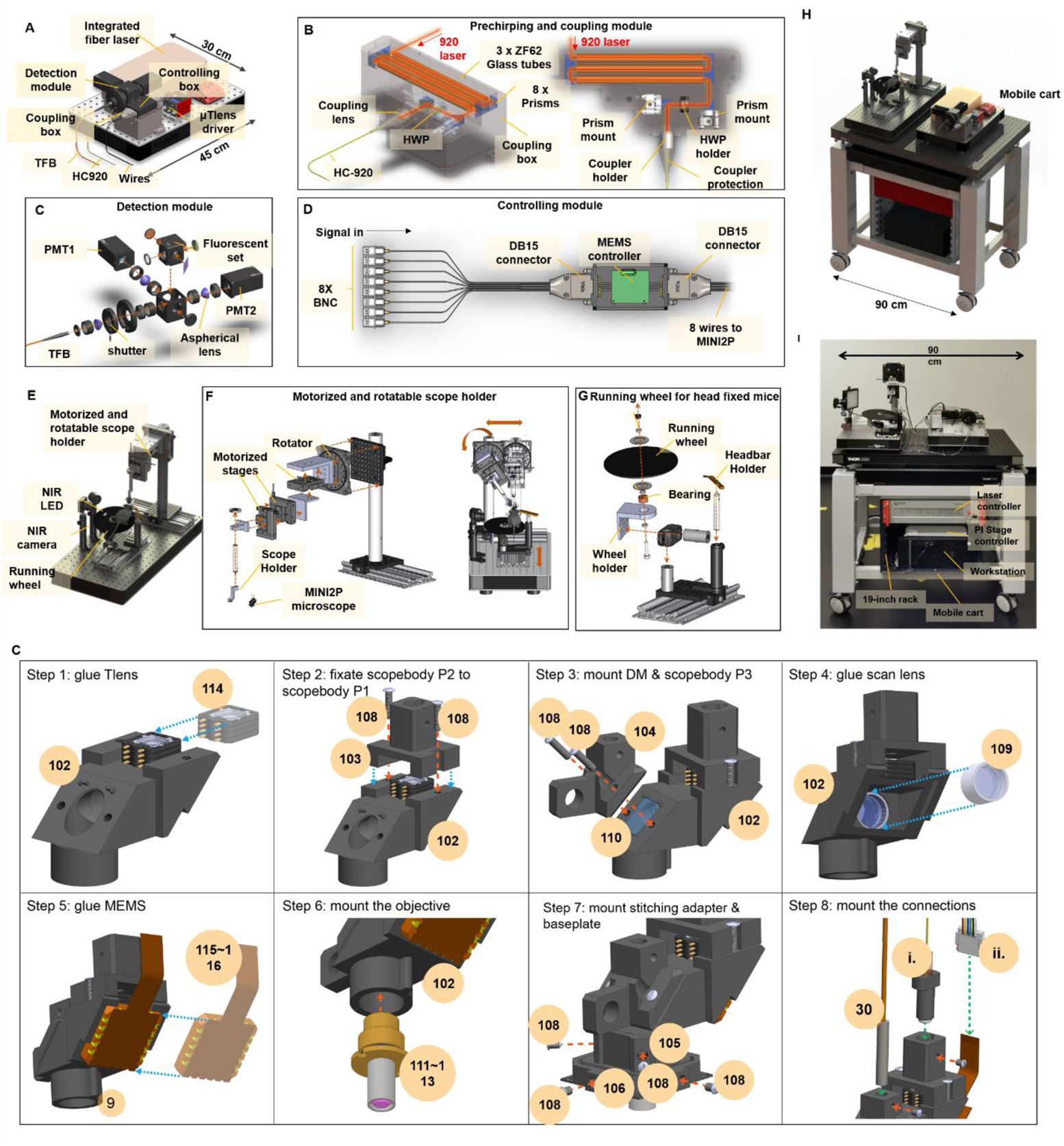
Detailed design and building illustration of MINI2P system. Related to STAR Methods. (**A-D**) The core optics module. (**A**) Overview. All parts can be integrated on a 45 cm x 30 cm optical breadboard. (**B**) Assembling schematic of the prechirping and coupling module. 3 x 15cm (diameter: 10 mm, 45 cm long in total) ZH62 glass tubes were used to compensate the positive dispersion of 2.5-meter-long HC-920 hollow-core photonic-crystal-fiber. Different lengths and numbers of glass tubes can be chosen according for different lengths of HC-920. A half wave plate (HWP) was used to eliminate double-pulse effects in the fiber (see STAR Methods). (**C**) Assembly schematic of two-channel detection module. (**D**) Assembling schematic of controlling module. (**E-G**) Scope-mounting module. (**E**) Overview. NIR LED and NIR camera were used to monitor position of the MINI2P microscope and the animal during the microscope mounting. The animal was head-fixed and running on the wheel during microscope mounting. (**F**) Assembling schematic of the motorized and rotatable scope holder. The motorized stages moved the MINI2P microscope in xy- and z- axes with 1-µm resolution. The rotator allowed the microscope mounting angle to be adapted for different brain regions. Right: procedure for adjustment of microscope mounting angle. (**G**) Assembling schematic of running wheel for head fixation. (**H**) Overview of the complete MINI2P system. (**I**) Photo of a MINI2P system. (**J**) Step-by-step assembly illustration. Complete assembling protocol, and material list with all 2D drawings and 3D models are provided (see key resources table). Numbers in yellow circles indicate item ID in the material list.

**Table S1.**
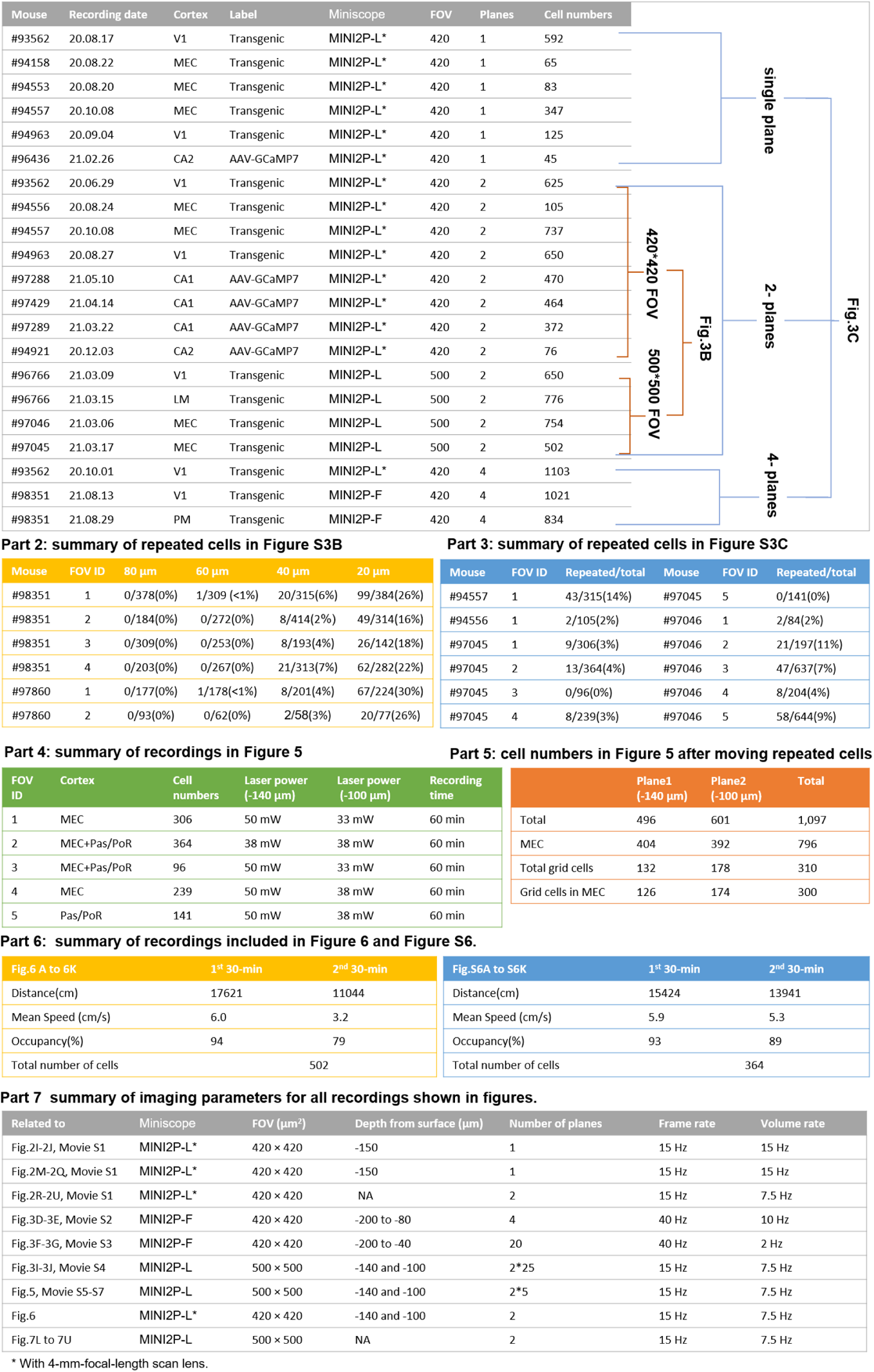
related to all imaging data. Part 1: Settings and data for all MINI2P recordings included in Figure 3B and 3C, across animals and brain regions (includes V1, LM, PM, CA1, CA2, MEC). The number of cells is the original number extracted from Suite2P without applying any filtering. If multiple recordings with the same parameter settings (420×420 μm^2^ FOV, 500×500 μm^2^ FOV, single plane, multiple planes) were found in the same animal, we only included the recording with highest number of cells. Currently we only have 2-plane 500×500 µm^2^-FOV imaging data for VC and MEC. For a fair comparison, we only used 2-plane 420×420 µm^2^-FOV imaging data in Figure 3B. Part 2: Summary of the number and fraction of repeated cells in VC recordings included in Figure S3B, with different plane intervals. Part 3: Summary of the number and fraction of repeated cells in MEC recordings included in Figure S3C, with a constant plane interval of 40 μm. Part 4: Imaging parameters for the experiment in Figure 5. Part 5: Number of cells after removing repeated cells in Mouse #97045. Part 6: Summary of recordings included in Figure 6 and S6 shows effect of consecutive recording sessions on behavioral performance as well as detection of grid cells. Part 7: Summary of imaging parameters for all recordings shown in figures.

## KEY RESOURCES TABLE

| REAGENT or RESOURCE | SOURCE | IDENTIFIER |
| --- | --- | --- |
| **Antibodies** |  |  |
| Chicken polyclonal to GFP (primary antibody) | Abcam | AB_13970 |
| Goat anti-chicken IgY (H+L), Alexa Fluor 488 (secondary antibody) | Thermo Fisher Scientific | Catalog #A-11039, RRID AB_2534096 |
| **Bacterial and Virus Strains** |  |  |
| AAV1-syn-GCaMP7f | Addgene,(Dana et al., 2019) | RRID:Addgene_ 104488 |
| AAVretro.CAG.tdTomato | Addgene, (Tervo et al., 2016) | RRID:Addgene_59462 |
| **Experimental Models: Organisms/Strains** |  |  |
| Mouse wild type C57BL/6JBomTac | Taconic | Cat#B6JBOM; RRID: IMSR_TAC:b6jbom |
| Mouse transgenic B6;DBA-Tg(tetO-GCaMP6s)2Niell/J | (Wekselblatt et al., 2016) | RRID:IMSR_JAX:024742 |
| Mouse transgenic B6.Cg-Tg(Camk2a-tTA)1Mmay/DboJ | (Wekselblatt et al., 2016) | RRID:IMSR_JAX:007004 |
| **Software and Algorithms** |  |  |
| Labview 2020 | national instruments | http://www.ni.com/labview/<br>RRID:SCR_014325 |
| Matlab 2020a | MathWorks | https://se.mathworks.com/products/matlab.html<br>RRID: SCR_001622 |
| Solidworks 2020 | Dassault Systèmes | https://www.solidworks.com/#_ga=2.204840921.81351093.1631434699-79998290-e304-11eb-a0b7-f50a6ae89b52 |
| Zemax 2019 | Zemax | https://www.zemax.com/ |
| Fiji | Fiji | http://fiji.sc/ ; RRID: SCR_002285 |
| ScanImage 2021 (matlab) | Vidrio Technologies (Pologruto et al., 2003) | https://vidriotechnologies.com/scanimage-v2021/ |
| Suite2p (python) | (Pachitariu et al., 2017) | https://github.com/MouseLand/suite2p |
| DeepLabCut (python) | (Mathis et al., 2018)<br>(Nath et al., 2019) | https://github.com/DeepLabCut/DeepLabCut |
| AnimalTracker (labview program) | This paper | https://github.com/kavli-ntnu/MINI2P |
| DistortionCleaner (matlab application) | This paper | https://github.com/kavli-ntnu/MINI2P |
| NATEX (matlab application) | This paper | https://github.com/kavli-ntnu/MINI2P |
| StitchingChecker (matlab application) | This paper | https://github.com/kavli-ntnu/MINI2P |
| Spatial tuning analysis code (matlab) | This paper | https://github.com/kavli-ntnu/MINI2P |
| **System construction materials** |  |  |
| 3D models and 2D drawings | This paper | https://github.com/kavli-ntnu/MINI2P |
| Building illustration | This paper | Supplementary Document, Section 8-10 |
| Shopping & Machining List | This paper | Supplementary Document, Section 11 |

## *STAR* METHODS

### RESOURCE AVAILABILITY

#### Lead contact

Further information and requests for resources and data should be directed to and will be fulfilled by the lead contact, Edvard I. Moser.

#### Corresponding authors

Weijian Zong; Edvard I. Moser;

#### Data and code availability

The data for each figure in this manuscript will be available as of the date of publication in a standardized format accessible in Matlab. DOIs will be listed in the Key Resources Table.

Any additional information required to reanalyze the data reported in this paper is available upon request.

### EXPERIMENTAL MODEL DETAILS

#### Animals

Experiments were performed according to the Norwegian Animal Welfare Act and the European Convention for the Protection of Vertebrate Animals used for Experimental and Other Scientific Purposes, permit numbers 18013, 6021 and 7163. The animals were housed under conditions free from specific pathogens according to the recommendations set by the Federation of European Laboratory Animal Science Associations (Mahler et al., 2014). We used wild-type (C57BL/6JBomTac) as well as transgenic mice, which express GCaMP6s ubiquitously under control of the CaMKII promoter (Camk2a-tTA; tetO-G6s, (Wekselblatt et al., 2016).

Behavioral experiments in Figure 1 and Figure S1 are from 3 male and 2 female transgenic mice, and 5 male wild-type mice. Statistics of cells number in Figure 3B and 3C are from 7 male and 3 female transgenic mice, and 3 male and 2 female wild-type mice. VC data in Figure 2I-L, Figure 3D-G, and Figure S3A and B are from three male transgenic mice and VC data in Figure 3H-M and the data in Figure 4 was recorded in another male transgenic mouse. MEC data in Figure 2M-Q are from one male transgenic mouse. MEC data in Figure 5 and Figure 6 are from another male transgenic mouse. CA1 imaging data in Figure 2R-U and Figure 7 were obtained from two male wild-type mice pre-injected with AAV1-syn-GCaMP7f virus. 10 additional wild-type mice were used in the behavior experiments in Figure S1F-K to show the impact of the connection cable on the animals’ behavior.

All mice were 12-24 weeks old at the time of surgery and 16-28 weeks at the start of behavioral pre-training. Before implantation, the mice were group-housed with up to three littermates (cage size: 32 × 17 cm^2^, 15 cm height). Before surgery and testing, the mice were handled and pre-trained in the recording environment until they were familiar with the environment and foraged freely in the box. After implantation, the mice were housed individually in transparent Plexiglass cages (36 × 24 cm^2^, 26 cm height). The mice were maintained on a 12 hr light/12 hr dark schedule and tested in the dark phase. The mice were never put on food or water restriction. Mouse health was checked daily.

### METHOD DETAILS

#### Visual cortex (VC) surgery

Anesthesia was induced by placing the animal in a closed glass box filled with 5% isoflurane flowing at a rate of 1.2 L/min. The mice were subsequently moved rapidly into a stereotaxic frame, which had a mask connected to an isoflurane pump. Airflow was kept at 1.2 L/min with 0.5–2% isoflurane as determined by physiological monitoring. The mice were then given two pain killers (Temgesic and Metacam) underneath the skin on the back, plus a local anesthetic (Marcain) underneath the skin over the durotomy site. Mannitol (150mg/ml) was also administered by intraperitoneal injection 30–60 min before the durotomy to reduce intracranial pressure. Then the scalp was resected to expose the entire dorsal surface of the skull. The periosteum was removed, but the bone was left intact. After exposing the skull, a circular craniotomy with a diameter slightly greater than 4.6 mm was made over the left hemisphere (centered 2.4 mm lateral to the midline and 2.8 mm posterior from bregma). The dura was kept intact. To increase the anchoring strength, a thin layer of Optibond (Kerr, CA, USA) was applied to the skull, and a thicker layer of Charisma (Kulzer, Hanau, Germany) was applied on top of Optibond. Both materials were cured with UV light (Kulzer, Hanau, Germany). Then a 4.6-mm-diameter customized coverglass (Sunlight, Fuzhou, China) and a custom titanium headbar, glued together with optical adhesive (NOA61, Norland, NJ, USA), were inserted into the craniotomy by using a custom-made headbar holder. During the insertion, the headbar holder was connected to a stereotactic micromanipulator (1760, Kopf, CA, USA) and tilted 20 degrees laterally to ensure that the cover glass would be approximately parallel to the dorsal surface of the VC cortex. The manipulator drove the headbar to go down until the glass touched the brain. Then the gap between the headbar and the skull was sealed with Venus (Kulzer, Hanau, Germany). Next, all other gaps between the headbar and the skull were filled with dental cement (Metabond, Parkell, NY, USA). After the dental cement had cured, the headbar was released from the holder. After the implantation, the chronic window was covered in Kwik-Cast (World Precision Instruments, FL, USA).

#### Medial entorhinal cortex (MEC) surgery

The surgical protocol was developed and optimized based on previous work (Low et al., 2014). Anesthesia was induced by placing the animal in a closed glass box filled with 5% isoflurane flowing at a rate of 1.2 L/min. Afterwards, the mice were rapidly moved into a stereotactic frame, which had a mask connected to an isoflurane pump. Airflow was as for VC surgeries, as were analgesics and local anesthetic. Mannitol (150mg/ml) was also administered by intraperitoneal injection 30–60 min before the durotomy to reduce intracranial pressure. The scalp was resected to expose the entire dorsal surface of the skull. The periosteum was removed, but the bone was left intact. After exposing the skull, a circular craniotomy with a diameter of 5 mm was made over the left hemisphere (centered 3.8 mm lateral to the midline and with the center of the transverse sinus). Because of the large craniotomy size, the left edge of the craniotomy window might fall out of the squamosal bone, so the muscle there was carefully released from the skull. The dura over the cerebellum was removed to allow micro prism insertion into the transverse fissure. The dura over the cortex was kept intact. The pitch of the mouse’s head was adjusted to make the anterior edge and posterior edge of the cranial window parallel to the horizontal plane. The prism assembly, made by a titanium cannula (5-mm diameter, 1-mm depth, custom), a 5-mm diameter coverglass (CS-5R, Warner, MA, USA), and a 1.3×1.3×1.6 mm^3^ prism (Figure 2M, custom, Sunlight, Fuzhou, China), glued together with optical adhesive (NOA61, Norland, NJ, USA), was held and implanted by a custom-made holder, with the prism inserted into the subdural space within the transverse fissure (Figure 2N and Figure 5A). During the implantation, the cannula holder was connected to the stereotactic micromanipulator (1760, Kopf, CA, USA) and tilted 20 degrees laterally to make sure that the cover glass was approximately parallel to the dorsal surface of the brain at the implantation site. The prism assembly was moved down slowly (less than 100 μm/s) until the cover glass was flush against the brain. Then the gap between the cannula and the skull was sealed with Venus (Kulzer, Hanau, Germany). Next, we added a custom headbar holder on top of the cannula, and all other gaps between the headbar and the skull were filled with dental cement (Metabond, Parkell, NY, USA). After the implantation, the chronic window was covered with Kwik-Cast (World Precision Instruments, FL, USA).

During the MEC surgery, we injected a retrogradely transported AAV into the dentate gyrus (DG) and CA3 to label MEC layer 2/3 cells with projections to these two regions (Tervo et al., 2016). After exposing the skull with a scalpel blade, a 0.5 mm craniotomy was made 1.7 mm posterior to bregma and 1.0 mm lateral to the midline for targeting the DG. The other 0.5 mm craniotomy was made 1.7 mm posterior to bregma and 2.1 mm lateral to the midline for targeting the CA3. 200 nL of rAAV2-CAG-tdTomato (Addgene, plasmid # 59462; RRID: Addgene_59462, no dilution) was injected at a rate of 20 nL/min, with a glass capillary (Nanoject III), 1.8 mm ventral to the surface of the brain in the first injection site (DG), and 1.7 mm ventral to the surface of the brain in the second injection site (CA3). In both injections, the glass capillary was left in place for 10 minutes before retraction. After retraction, the craniotomy was covered with an absorbable hemostatic gelatin sponge (Spongostan) and a biocompatible silicone sealant (Kwik-Cast). The injections were made before craniotomy was made in MEC.

#### Hippocampus surgery

The CA1 surgical protocol was developed and optimized based on previous work (Barretto et al., 2011). Mice were placed in a closed glass chamber filled with 4% isoflurane, flowing at a rate of 1.2 L/min. After induction of anesthesia, they received an intraperitoneal injection of Mannitol (150 mg/ml) and were transferred to a stereotactic frame where isoflurane was delivered through a nose cone. Air flow was then kept at 0.4 L/min with 0.5-1.5% isoflurane as determined by monitoring physiological parameters throughout the surgery. Subcutaneous injections of Temgesic (0.03 mg/ml), Metacam (1mg/ml), Saline (9 mg/ml) and Marcain (0.5 mg/ml) were given prior to head fixing the animal in the stereotactic frame. After exposing the skull with a scalpel blade, a 0.5 mm craniotomy was made 2.0 mm posterior to bregma and 1.5 mm lateral to the midline. 1 μL of AAV1-syn-GCaMP7f (Addgene, plasmid # 104488; RRID: Addgene_ 104488, diluted 1:9 with dPBS) was injected at a rate of 50 nL/min, with a glass capillary (Nanoject III) 1.2 mm ventral to the surface of the brain, and was left in place for 10 minutes before retraction (Dana et al., 2019). After retraction, the craniotomy was covered with an absorbable hemostatic gelatin sponge (Spongostan) and a biocompatible silicone sealant (Kwik-Cast), and the skin was closed with sutures.

2 weeks after injection, the mice were implanted with a medical-grade stainless steel cannula (1.8 mm outer diameter, 3.1 mm length) with a coverslip glued to the end. A new craniotomy was made by using a 1.8 mm diameter trephine drill and centering it at the previously made injection craniotomy. Overlying cortex was carefully aspirated under continuous irrigation with cold saline until white-matter fibers became visible. White-matter fibers running in the mediolateral direction were gently peeled away, leaving the alveus intact. After aspiration, the cavity was intermittently irrigated and filled with Spongostan until all bleedings had stopped. The guide cannula was then lowered into the cavity until the coverslip touched the brain, approximately 1.1 mm DV from the skull. The tiny gap between the cannula and the skull was sealed with topical skin adhesive (Histoacryl) and flowable composite (Venus Diamond Flow). The whole skull was then covered with dental adhesive (Optibond) and composite (Charisma Diamond). A custom-made titanium head-bar was sealed to the skull and cannula with dental cement (Meliodent) mixed with carbon powder. The cannula was covered with Kwik-Cast to protect any light and dust from entering it.

One week after the surgery, animals were head-fixed on a running wheel, and a GRIN lens (1 mm diameter, 4.3 mm length, Grintech, Germany) was inserted into the cannula and glued onto the coverslip with optical adhesive (NOV61, Norland, USA).

#### Histology

After finishing experiments, the animals were first anesthetized with isoflurane and then euthanized with an overdose of pentobarbital after which they were perfused, first with 0.9% physiological saline followed by 4% formalin or paraformaldehyde (PFA). The implantation sites were carefully dissected, with a small fraction of skull remaining attached, where the prism was located, to allow fixation of the cortex and cerebellum with the prism still in place. After fixation in 4% formalin/PFA for at least one week the brains were dislodged from the skull, carefully as to maintain the connection between the cortex and cerebellum. The brains were then sliced in either 30 µm sagittal sections (VC), 50 µm horizontal sections (MEC), or 50 µm coronal sections (CA1) with a cryostat (Thermo Scientific Microm HM 560). The slices were stained in two series with a standard cresyl violet (Nissl) protocol and a fluorescent GFP staining. To stain for GFP, free-floating slices in PBS were washed in 3% BSA and 0.3% TX100 in PBS for 2 x 5 min. They were then incubated in the primary antibody (anti-GFP chicken, Abcam, RRID: ab13970, diluted 1:1000 with PBS) in 3% Bovine Serum Albumin (BSA) and 0.3% TX100 in PBS on a shaker in a cold room overnight. The slices were then washed in the BSA/TX100/PBS solution another 2 x 5 min before incubation of the secondary antibody (goat anti-chicken IgY (H+L) secondary antibody, Alexa Fluor 488, Thermo Fisher Scientific, catalog # A-11039, RRID AB_2534096., diluted 1-800-1:1000 with PBS) in the aforementioned BSA solution for one hour in room temperature on a shaker. Finally, the slices were washed 2 x 5 min in PBS (on a shaker in room temperature) before being mounted with ProLong™ Gold Antifade Mountant with DAPI (Invitrogen, catalog # D1306). The position of the prism and visualization of the staining were acquired with an Axio Scan Z1 microscope and Axio Vision software (Carl Zeiss, Jena, Germany).

#### Open-field task

Recording began after complete recovery from surgery. All mice were habituated to handling and the behavioral task gradually to a satisfactory level prior to any experimentation, and each day the animals were placed in the recording room during final preparations in order to allow for familiarization to the surroundings. The task was implemented in a dark room, where the sole light source was a warm-light LED strip in the ceiling to promote exploration. A session consisted of a single recording lasting either 30 min (each of the 22 FOVs of VC imaging in Figure 3H-M and Figure 4, CA1 imaging in Figure 7, and the behavior experiments shown in Figure 1), or 40 min (each of the 3 FOVs of VC imaging in Figure 3H-M and Figure 4), or 1 hour (MEC imaging in Figure 5); a pair of recordings in Figure 6 lasted 30 min each, with a 10 min break in between. Neural activity was recorded as the animal freely explored an 80×80 cm^2^ square black box (VC and MEC recording in Figures 4–6) or a 55×55 cm^2^ square black box (CA1 recording in Figure 7), in both cases with a single sheet of white laminated A4 paper fixed to one wall of the box, serving as a polarizing cue. The animal was placed into the box and allowed to move around freely while the experimenter periodically threw crumbs of cookies into the environment. An NIR camera was used to record the position of the animal from above.

#### Connection cable assembly in open field task

The “thin” connection cable assembly consisted of a HC-920, a 6-core electric wire, and a 0.7-mm-diameter fiber bundle of 100 to 200 0.05-mm-diameter glass fibers. The “thick” connection cable assembly was made by adding a 1.3-mm fiber bundle with the same glass fibers to the “thin” connection cable assembly. The 0.7-mm-diameter fiber bundle and 1.3-mm fiber bundle together mimicked the 1.5-mm-diameter fiber bundle used in the previous version of 2P miniscope (Zong et al., 2021; Zong et al., 2017). The connection cable assembly was held in place over the center of the open field with a thin nylon strain connected to a bearing on the ceiling of the room and connected to a counterweight at the other end. The length from the holding part of the cable to the MINI2P microscope was about 1 meter, long enough for the animals to run freely into all corners of the box. No commutator was used. The flexibility of the connection cable assembly allowed for more than 10 circles of twisting without restricting the animaĺs behavior. In most experiments, animals ran for over 30 mins without obvious tangling of the cable. In rare cases where the cable restricted the animals’ behavior, we stopped the experiment, moved the animal out of the box, de-twisted the cable, put the animal back, and continued the experiment.

#### Tracking of animal behavior

The behavior of the animals in the 80×80 cm^2^ (or 55×55 cm^2^) black-floor square box was monitored by a monochrome camera (acA2040-90umNIR, Basler, Ahrensburg, Germany) mounted on the top of the box with a 16-mm lens (C11-1620-12M-P f16mm, Basler, Ahrensburg, Germany). The environment and the floor were illuminated uniformly by 8-12 850 nm LED lights mounted at different angles and regular positions around the box. An 875-nm short-pass filter (#86-114, Edmund, NJ, USA) was mounted in front of the camera to filter out 920 nm laser light leaking from the miniature microscope. The tracking video was recorded in external mode with either the camera company’s software (Pylon, Basler, Ahrensburg, Germany) or a custom LabView program (*AnimalTracker*, see Key Resources Table). At the start of each frame of the 2P imaging, a TTL pulse (0-5V) was generated from the vDAQ card (see STAR Methods, “MINI2P system” section), which triggered one frame recording of the animal tracking camera. In this way, the animal tracking video acquisition was in sync with the 2P frame acquisition at all times.

#### Behavioral testing of miniscope weight and cable flexibility

To investigate the effect of miniscope weight and the flexibility of the connection cable on the behavior of the animals, two custom-made dummy miniscopes (3g and 5g) were used together with two different connection cable assemblies (“thin” and “thick”). The dummies simulated the shape and weight distribution of the miniscope and could be mounted directly onto implanted headbars using a single screw. 10 animals were used in total, including 5 transgenic mice with real implants (3 in VC and 2 in MEC), and 5 wild-type animals which had received sham implants (no cranial window, only headbar implantation).

The purpose of these sets of experiments was to compare the overall effect of miniscope versions on behavior and ascertain the extent to which each component (scope weight and connection cable) results in divergence from naturalistic, freely moving behavior. The main experiment was designed to compare the combined impact of weight and cable, with reference to the previous version of the 2P miniscope (Zong et al., 2021). Hence, Figure 1 shows data collected successively (in the same animals) with the 3g dummy microscope and the thin cable assembly, simulating the current MINI2P, and the 5g dummy microscope with the thick cable assembly, simulating the previous version of 2P miniscopes (Zong et al., 2021). A control session with no miniscope and no cable was also conducted. To better understand which combination of components impacts the behavior the most, additional experiments were subsequently designed to compare fiber and scope weight independently (Figure S1). In the comparison of connection cable assemblies, we kept the scope weight constant (3g) and used either the thin cable (3g-t) or the thick cable (3g-T), while in the scope weight comparisons, we used the same thin cable, with either 3g dummy (3g-t) or 5g dummy (5g-t) miniscopes. The comparison of connection cable assembly also included 10 additional sessions in each group, obtained from recordings with earlier versions in the lab. The recording order in each animal was pseudorandomized. Before recording started, the animals were fitted with the miniscope dummy (which was pushed onto the implanted head bar and secured without head fixation) and then gently placed into the open field arena, where they were left to adjust to the box and the darkness for a few minutes. Then we started the 30-min free-foraging open-field task with the same behavior recording setup as used in other experiments. After the session was over, the animals were gently scooped up from the open field in a familiar tube, the dummy scope was removed, and the animals were let back into their home cage. All testing started at the same time every day, and a maximum of two animals were tested per day to control for natural fluctuations of activity level throughout the day.

To inspect the behavior of the animal in each recording, 6 types of analyses were conducted, comparing animals in respect to the accumulated running distance over time, total running distance, average speed, peak running speed, coverage in the center of the box, and tortuosity of the animaĺs running path. For this, the *x* and *y* head position, *X*(*t_i_*) and *Y*(*t_i_*), were extracted from the tracking data processing pipeline (see below) and smoothed with a 0.5 s moving average. *t_i_* denotes the ith frame (time point) in the whole recording. Then the inter-frame running distance *d*(*t_i_*) between successive time points (frame interval ΔT = 0.07s) at time point *t_i_* was calculated: *d*(t_i_)=sqrt([X(t_i_)-X(t_i-1_)]^2^+[Y(t_i_)-Y(t_i-1_)]^2^). The momentary speed *S*(t_i_) was calculated with the inter-frame running distance and the frame interval as *S*(*t_i_*)= *d*(*t_i_*)/ ΔT. The accumulated distance over time was calculated as the cumulative sum of all inter-frame distances within a session 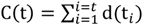, where *i* is the time point number from the start (i=1) to the current time point *t* of the recording (i=t). The total running distance was calculated as the integral over all inter-frame running distances. Average and median speed estimates were derived over the whole recording, and the top speed was defined as the 90^th^ percentile of the momentary speed in the whole recording. Then the total running distance and time spent within ±15 cm in the *x* direction and ±15cm in the *y* direction around the center of the box was calculated (“total time in center” and “total distance in center”).

#### Tortuosity

The animals’ momentary tortuosity was defined as the ratio of the length of the path (**L**) to the straight-line distance between its ends (**D**), with a ±1.25-second sliding window for each time point of the recording. Initially we tried several lengths of sliding windows (1s, 2s, 2.5s, 5, 7.5s, 10s) and plotted the momentary tortuosity on animal trajectories. We found that the 2.5s sliding window gave the best resolution of turning behavior when the animals were running. The *x* and *y* head positions *X*(*t*) and *Y*(*t*) were smoothed with a 0.5 s smoothing average. The length of the path, *L*(*t*), between ends of a ±1.25s sliding window was computed as the integration of inter-frame running distance 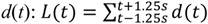. Then we computed the straight-line distance between the ends of each sliding window, *D*(*t*), as the Euclidean distance from the start frame (t-1.25s) to the end frame (t+1.25s). The momentary tortuosity at timepoint *t* was then calculated as *T*(*t*)=*L*(t)/*D*(t). The momentary tortuosity was assessed only when the animals were running, we selected the speed threshold (7.3 cm/s) as the average of the 75^th^ percentile of speed in all control experiment recordings (10 animals); 50^th^ and 75^th^ percentiles of the tortuosity distribution were then extracted only from frames with speeds higher than this threshold. The mean turning speed was calculated from frames with tortuosity higher than the tortuosity threshold (a value of 1.96, equivalent to the average of the 75^th^ percentile of tortuosity in all control experiment recordings (10 animals)), for which the animal moved faster than 2.5 cm/s. We finally identified the 50^th^ and 75^th^ percentiles of running speed of the frames with tortuosity higher than this threshold for each recording.

#### MINI2P System

The MINI2P system included a core optics module, a scope-mounting module, and a controlling module (Figure S7A-I). We have provided the complete shipping/machining list, detailed building protocol of the MINI2P system and all 3D models and 2D drawings for custom components (see Key Resources Table).

The core optics module comprised all the optical components other than the miniature microscope (Figure S7A). The laser source was a compact, single-wavelength, fiber-based femtosecond laser (FemtoFiber ultra 920, Toptica, Munich, Germany; 920 nm +/− 10 nm, 100 fs, 80 MHz, 1.2 W). Group-delay dispersion compensation (−40000 to +1000 fs²) and fast analog power modulation (>1 MHz bandwidth) were integrated. Compared to tunable Ti-sapphire lasers, single-wavelength fiber lasers have benefits such as a smaller footprint, a lower price, much lower operation noise, fewer maintenance requirements, and the wavelength matches the narrow bandwidth of the hollow-core photonic crystal fiber (HC-920, K50-060-00, NKT, Copenhagen, Denmark; bandwidth: 900-950 nm). In the coupling box (Figure S7B), the laser beam is reflected by 8 right-angle prisms (MRA12-P01, Thorlabs, NJ, USA) and goes through three 15-cm-long ZF-62 glass tubes (GLA-10×150-AR800-1100, Sunlight, Fuzhou, China). The ZF-63 glasses gave a constant positive dispersion compensation, and the laser’s internal dispersion compensator minimized the residual dispersion. The number and length of the ZF-62 glass tube can be changed according to the different lengths of HC-920 (Zong et al., 2017). Due to the terminal reflection, birefringence, and other nonlinear effects during the transmission in the HC-920, a double-pulse effect might occur during the laser transmission in the fiber, which decreases the peak power and the two-photon effect. This double-pulse effect can be observed by using an autocorrelator (PulseCheck, APE, Berlin, Germany) as two small side peaks beside the center autocorrelation signal. By rotating the laser’s polarization using a half-wave plate (HWP, WPHSM05-915, Thorlabs, NJ, USA) before the laser was coupled into the photonic crystal fiber (HC-920, NKT, Denmark), the second pulse could be eliminated. The laser was then coupled into the termination-processed HC-920 fiber assembly (Supplementary Document, Section 8) with a coupling lens (SM1T2, Thorlabs, NJ, USA). The coupling efficiency was monitored by measuring the laser power output from the other end of the HC-920 assembly with a power meter (S121C+PM100D, Thorlabs, NJ, USA). By iteratively tuning two mirror mounts (POLARIS-K05S2, Thorlabs, NJ, USA) and adjusting the distance between the HC-920 assembly’s termination (in the glass flange) and the coupling lens (in the coupling holder), the coupling efficiency can usually reach above 75% (Supplementary Document, Section 8). After the coupling efficiency reached a maximum, the fiber end, glass flange, and the coupling holder were glued together with UV glue (NOA61, Norland, NJ, USA). The HC-920 fiber should stay on the holding stage for at least 24 hours until the UV glue stops shrinking. The optical alignment was stable with the tight coupling design of a minimized number of optics components. No active vibration-isolating equipment was required. The output power decrease was less than 1% over three months, so frequent re-alignment during the daily operation was unnecessary. The mirror mount’s adjustment was accessible without opening the coupling box. In the detection module (Figure S7C), an aspheric condenser lens (ACL25416U-A, Thorlabs, NJ, USA) first collimated the TFB output. Then a filter set (FESH0750/DMLP567R/MF525-39/MF630-69, Thorlabs, NJ, USA) split the light into a short-wavelength component (500-550 nm) and a long-wavelength component (595-665 nm). The two different light paths were then focused on GaAsP photomultiplier tubes (PMT, PMT2101/M, Thorlabs, NJ, USA) by aspheric condenser lenses (ACL25416U-A, Thorlabs, NJ, USA). The MEMS driver (for BDQ PicoAmp 5.4 T180, Mirrorcle, CA, USA) was in the controlling box with two DB-15 connectors (Figure S7D). The TLens driver (Thorlabs, NJ, USA) generated high voltage for the µTlens. A mechanical shutter (SHB1, Thorlabs, NJ, USA) was put in front of the filter set to protect the PMTs from unexpected exposure.

The scope-mounting module served to mount the miniature microscope on the animal’s head (Figure S7E). We used a scope holder consisting of three motorized linear stages (M-112.2DG1, Physik Instrumente, Karlsruhe, Germany) and a manual rotation stage (RP03/M, Thorlabs, NJ, USA, Figure S7F). A hardboard (TB4, Thorlabs, NJ, USA) was cut into a 17-cm-diameter circle and sprayed with rubber spray to be used as a running wheel (Figure S7G). A 6-mm hole was drilled in the wheel’s center to mount spacers, a bearing (626-2Z, SKF, Gothenburg, Sweden), an M6 screw, and a custom-made wheel holder. We used one near-infrared (NIR) light-emitting diode (LED, LIU850A, Thorlabs, NJ, USA) and one NIR camera (CS165MU/M, Thorlabs, NJ, USA) with a lens (MVL8M23, Thorlabs, NJ, USA) to monitor the microscope and the animal’s behavior. A headbar holder held the headbar from a single side and left free space for mounting the microscope and monitoring the animal’s behavior on the other side.

The controlling module was placed in a 19-inch rack (9U, Schroff, BW, Germany) containing a workstation computer (7080, Dell, TX, USA), the laser controller, and the PI-stage driver (C-884.4DC, Physik Instrumente, Karlsruhe, Germany) (Figure S7H and I). The FPGA card, vDAQ (Vidrio Technologies, VA, USA), plugged into the workstation, took over all hardware control and data acquisition. Finally, the scope mounting module, the core optics module, and the controlling module were integrated into a mobile cart (POC001, Thorlabs, NJ, USA) with a breadboard (PBG52506, Thorlabs, NJ, USA).

#### µTlens

Individual Tlenses were constructed by a layer of glass membrane attached to a circle MEMS piezo actuator (thickness: 0.05 mm), a layer of transparent polymer (thickness: 0.1 mm), and a layer of thin glass (thickness: 0.03 mm) (Figure 2C). Depending on the voltage applied, the MEMS piezo bends the glass membrane to generate a divergent lens, a flat lens, or a convergent lens. For MINI2P, four Tlenses were centered and stacked together, and the alignment accuracy was determined by measuring the wavefront error under an interferometer. Five µTlenses produced in different batches had identical optical power responses (Figure S2C), a similar wavefront error, and high transmission in the near-infrared wavelength band (>95% from 900-1000 nm, Table 1, Part 1).

To enable precise multi-layer and 3D imaging with the µTlens, we first calibrated the change of focus by matching the focus with a known movement distance using the scope-holding stage. The calibration curve (Figure 2D) was then saved in the ScanImage machine data file and automatically applied to adjust the µTlens control voltage when ScanImage was running.

#### Resolution and FOV measurement

We measured the point spread function (PSF) of the MINI2P microscope by using 1-micron fluorescence beads (F8776, ThermoFisher Scientific Inc., MA, USA, Supplementary Document, Section 2 and 3). Cross-sections along the z-axis centered on each bead were used to calculate the axial FWHMs. We took reflection images of 50-µm-grid test samples (R1L3S3P, Thorlabs, NJ, USA) to measure the FOV of the MINI2P microscope in different focusing planes (0 μm to 240 μm) of the µTlens.

#### Tapered fiber bundle (TFB)

The supple fiber bundle (SFB) used in the previous two versions of the 2P miniscope (Zong et al., 2021; Zong et al., 2017) had a diameter of 1.5-mm with 200-500 single fiber cores (diameter: 30-50 µm). An SFB with a smaller diameter was not efficient enough for 2P imaging (Figure S2D and S2E). To address this constraint, we designed and fabricated a new type of tapered fiber bundle (TFB) consisting of a tapered glass rod and a 0.7-mm-diameter fiber bundle (Figure 2B). Scattered fluorescence photons were collected by the larger end of the tapered glass (diameter: 1.5 mm) and, after multiple total internal reflections in the glass rod, the photons were coupled into the fiber bundle from the tapered end (diameter: 0.6 mm). Because the tapered glass also has a focusing effect, no collecting lens was required. The reflection index and the outline curvature of the tapered glass rod were carefully designed to match the numerical aperture (NA) of the objective (NA 0.15 at imaging plane) and the fiber bundle (NA 0.5), bringing the collection efficiency of the TFB to a level nearly identical to that of the 1.5 mm SFB and much higher than the single 0.7 mm SFB (Figure S2D and Figure S2E).

#### HC-920 assembly

To protect the HC-920 fiber and stabilize the coupling, the coupling lens, the HC-920 fiber, and the collimating lens were assembled as an “HC-920 assembly” (complete assembling illustration is in Supplementary Document, Section 8). If any part of the HC-920 assembly is damaged or a different length HC-920 is required, the whole assembly can be replaced. First, a 2.3-m Hytrel jacket (FT900Y, Thorlabs, NJ, USA) was put on 2.52-m HC-920 to protect the fiber. Then about 11 cm pure 920 fiber was exposed on both ends for termination processing. About 6 centimeters of plastic coating of the fiber on both ends was removed using a fiber stripping tool (T08S13, Thorlabs, NJ, USA). A 1-cm end of the fiber was cleaved using a fiber cleaver (XL411, Thorlabs, NJ, USA). The end of the fiber should be flat without any chipping, scratches, dirt, or pits under the microscope. After the end faces are processed, the fiber end was carefully inserted into a glass flange (TUB-1.8×6.5-0.127, Sunlight, Fuzhou, China) until the end of the plastic coating reached the smallest part of the inner taper. Then the glass flange’s inner taper was filled with optical adhesive (NOA61, Norland, NJ, USA), and it was cured with UV light. After both ends of the HC-920 fiber were processed and glued to the glass flanges, one flange is connected to the coupling holder to do the laser coupling. Next, the other glass flange was inserted into the collimator holder with a pre-mounted collimating lens (84-127, Edmund, NJ, USA) to make the. The distance between the glass flange and the collimator holder was adjusted using assistance tools (MBT602/M, Thorlabs, NJ, USA; collimator holding tool, custom-made) to collimate the output beam. After the output beam was collimated, the fiber end, the glass flange, and the collimator holder were glued together with optical adhesive (NOA61, Norland, NJ, USA). The HC-920 stayed on the assistance tools for at least 24 hours until the shrinking of the glue had stopped.

#### MINI2P miniscope

The output of the HC-920 assembly first went through the µTlens and then was reflected by the MEMS scanner (Figure 2A, complete optics drawing of MINI2P miniscope is in Supplementary document, Section 1). The position of the Tlens was designed to be as close to the MEMS scanner (2.34 mm) as possible to minimize the magnification change (difference <10%) across the whole scanning range. After the MEMS scanner, the laser beam passed the scan lens (Focal length: 5mm, D0166, Domilight, Nanjing, China), the dichroic mirror (MIR-4×5×0.2-HT450-650&HR800-1100, Sunlight Ltd., Fuzhou, Fujian, China), and then entered the objective (D0213/D0277/D0254, Domilight Ltd., Nanjing, China). All three objectives share the same threads (M5-0.5) and have an equal distance between the imaging plane and the mounting referent (the end of the threads), so that they are interchangeable. The emitted fluorescence goes through the dichroic mirror and is collected by the TFB (C54706, Schott, Germany). The microscope shells (Scopebody P1, P2, and P3 in Figure S7J) are manufactured from carbon-fiber-enhanced plastic (PEEK-CF30). The MEMS, the scan lens, and the dichroic mirror are glued directly to Scopebody P1. The HC-920 assembly, the TFB, and the electrical wires are plugged on Scopebody P2 and P3 and fixed by several M1.6 screws. The objectives are screwed on Scopebody P1. The delicate design of MINI2P improves matching accuracy so that the alignment and mounting of each component is very simple (Supplementary Document, Section 10). Assembling one MINI2P microscope can be broken down into 8 discrete steps, which altogether take about 1 hour of work (Figure 7J).

#### Control and acquisition

ScanImage (version 2020, Vidrio Technologies, VA, USA) fully supports the hardware control and data acquisition of MINI2P. Following the wiring illustration and the operation manual in Supplementary Document, Section 9, the system can be run directly without further modification. Two analog outputs of vDAQ send the MEMS scanning control signal (fast axial and slow axial) to the control box. The fast-axial frequency is 5600 Hz for the MEMS-F (A7M10.2-1000AL, Mirrorcle, CA, USA) and 2000 Hz for the MEMS-L (A3I12.2-1200AL, Mirrorcle, CA, USA).

The maximum peak to peak voltage (V_peak-peak_) should not exceed 8.5 V. The slow axial frequency for both MEMSs is set automatically by the software according to the frame rate. A third analog output sends the control signal to the µTlens driver (KPZ101, Thorlabs, NJ, USA) that provided 0-60V high-voltage for driving the µTlens. A fourth analog output sends the laser power control signal to the laser controller. Two digital outputs generate the MEMS driver’s lowpass filter modulation (check the MEMS driver’s manual). Another digital output provides the frame clock (D3.7 in vDAQ) to synchronize the animal tracking camera. The signals from two-channel PMTs are connected to two high-speed (120 MHz) analog inputs of the vDAQ card. A maximum of 4 channels can be acquired simultaneously. One high-speed analog input is reserved for registering external voltages (e.g. optogenetics stimulation, visual stimulation, treadmill speed, etc.) to the 2P imaging. The output of the controlling box is connected to the microscope wire assembly with a DB15 connector. The other end of the wire assembly is connected to the FPC on the microscope with a micro-FPC connector. The FPC is connected to the MEMS by direct surface soldering and to the µTlens by two wires soldered to the pins on the back of FPC. The details of system timing are shown in Supplementary Document, Section 9.

#### Optimization of the MEMS scanning

We established a mathematical description of the MEMS scanning based on geometrical optics. The geometry of the MEMS scanning is shown in Figure. S2H. In the static state (no driving force), the normal of the MEMS mirror (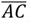) is placed at an angle of 45 degrees to the incident beam (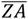) and the surface of the scan lens (named the “scanning plane”). The first scanning axis, the “slow axis”, is parallel to the scanning plane, and the second orthogonal scanning axis, the “fast axis”, is 45 degrees to the scanning plane. According to geometrical optics, the numerical solution of the laser spot position according to the scanning angles of the slow axis and the fast axis can be described as:

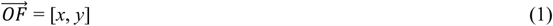

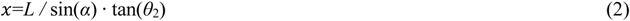

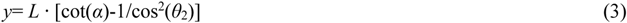

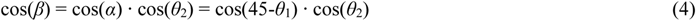

Details of the derivation can be provided upon request. *x,y* are the lateral and axial shifts of the scanned laser spot (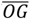 and 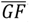) in relation to the static laser spot *O*. *L* is the length of the vertical line to the scanning plane (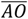) from the incident spot on MEMS (**A**). 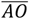 is equal to the reflection beam when the MEMS is in the static state. *θ*_1_ and *θ*_2_ are the scanning angles of the slow axis and the fast axis, respectively. *α* is the beam incident angle to the normal of the MEMS if the fast axis is in a static state and only the slow axis is scanning (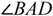). *β* is the beam incident angle to the normal of the MEMS (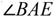), and is equal to the reflection angle based on the reflection law (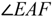). Here we have:

1. the scanning area is symmetric according to the 1^st^ axis but asymmetric according to the 2^nd^ fast axis, which generates a proximate trapezoid scanning area with the side length decreased according to *θ*_2_;
2. the waist of the trapezoid is approximately 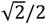 of the height when *θ*_2a_*= θ*_1a_; and approximately equal when 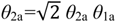 (see also Supplementary Document, Section 5);
3. the scanning line generated by the fast-axial scanning from −*θ*_2a_ to *+θ*_2a_ when the slow axis is at a certain angle *θ*_1_ is not straight but curved;

Conclusion 1) indicates that the fast-axial scanning range needs to be at least 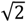 times of the slow-axial scanning range 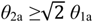 to achieve a relevant square field of view. 2) and 3) show that the final image has a systematic distortion consisting of the scanning field bending and size variation. Also, when one axis was running in a particular frequency region, the scanning angle was amplified (Figure S2G). In this region, the axis did not reach the resonant condition, so the scanning angle was still controllable, referred to as “quasi-resonant frequency.” This effect introduced a frequency-dependent nonlinearity of the MEMS mechanism. We found that we could use this effect to increase the scanning angle of the fast axis. The basic idea is to control the fast axis scanning at a working frequency (*F_w_*) in the “quasi-resonant region” where the amplification factor 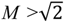, so that the scanning area will be increased by 30% compared to the static condition. Here we analyzed and tested two types of MEMS scanners: both MEMSs have a similar shape of the frequency response function: the magnitude is flat (magnitude~1) in the low-frequency domain, increases to the peak at a certain frequency (the first resonant frequency F_1st_), and then decreases monotonically in the high-frequency domain (MEMS-L, A3I12.2-1200AL, max scanner angle: +/− 5.2 degree, F_1st_=2000 Hz; and MEMS-F, A7M10.2-1000AL, max scanner angle: +/− 4.5 degree, F_1st_=4800 Hz). The typical frequency response (amplitude) of MEMS-L (orange curve) and MEMS-F (blue curve), derived from the Fourier transform of the full-range step response, are shown in Figure S2G and summarized in Table 1, Part 2. MEMS-L has a high Q-factor (the slope to the peak frequency), and MEMS-F has a low Q-factor. The magnitude is equal to the scanning angle’s amplification if the scanning is run with a sine wave (single frequency component). Thus, we limited the fast axis of the MEMS running in the frequency region with the magnitude 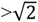 (the shallow pink area, referred to as “amplification allowed”, above the yellow dashed line in Figure S2G). For MEMS-L and MEMS-F, this gave: MEMS-L:

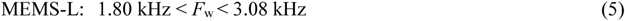

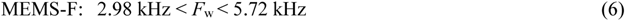

Working at the +/−20% range of the *F*_1st_ (the purple shallow, named “unstable region” in Figure S2G) was not stable, because the MEMS was easy to be permanently broken. Thus, the MEMS must work outside of the following ranges:

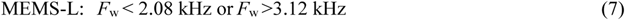

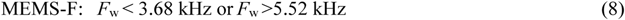

Combining (5), (6) and (7), (8), there was only one working region for MEMS-L (red shaded region, referred to as “MEMS-L working region” in Figure S2G):

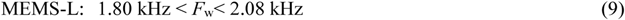

but for MEMS-F, there were two working regions (blue shaded region, referred to as “MEMS-F working region” in Figure S2G):

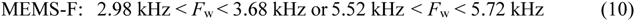

As a result, we set the *F_w_* to 2 kHz for MEMS-L, which gave a frame rate of 15 Hz for 256×256, and we set the *F*_w_ to 5.6 kHz for MEMS-F, which gave a frame rate of 40 Hz for 256×256. Both frequencies allowed stable scanning with full use of the maximum scanning angle of the MEMS.

#### Distortion correction

We developed an imaging-based calibration and distortion correction method to correct scanning field bending and magnification variation (Figure S3D). The key idea is to generate a series of transformation matrices in the different zoom, frame-size and µTlens focus positions, between the distorted image (raw image) and the corrected image (target image) in the calibration step. After each recording, the matrix for the same imaging parameter (zoom, frame size, focus) is used for correcting the distortion. To do this, we first imaged a series of standard 50 μm-grid distortion-test target (R1L3S3P, Thorlabs, NJ, USA) with different zoom (×1, ×2, ×3, etc.), frame-size (512^2^ or 256^2^) and focus points (0 to 240 um). Without distortion, the grid pattern should have horizontal and vertical lines in parallel (Figure S3D step1). The unparallel appearance of these lines indicates the distortion (Figure S3D step2). We made use of the crosses of the horizontal and vertical lines of the grids as landmarks (named “anchor points”) for generating the transformation matrix. First, several original anchor points (AP) were generated with a ground-truth grid pattern (blue dots in Figure S3D step1). Then we superimposed the original anchor points on the raw image. At this stage, most of the anchor points were mismatched with the crosses on the grid image (blue dots in Figure S3D step2). Then we dragged these points manually in the software to the positions of the correct crosses (Figure S3D step3). After we assigned each anchor point to one grid cross, a 2D piecewise linear transformation matrix was generated between the original anchor points (blue dots in Figure S3D step3) and the moved anchor points (red dots in Figure S3D step3). This transformation matrix (Matlab: “PiecewiseLinearTransformation2D”) controls pixels in the image by breaking up the image into local piecewise-linear regions. A different affine transformation map controls pixels in each local region. Finally, we applied this matrix to the imaging data (Figure S3D step 4) either before feeding the raw data into suite2p or afterwards (i.e. on motion corrected image stack, projections, and ROI data exported by suite2p). The corrected image is distortion-free so that all anchor points overlap with the crosses of the grids (Figure S3D step 4). The whole algorithm is included in a custom software *DistortionCleaner* (see Key Resources Table). All the FOV and PSF measurements in the paper were made after the aberration was corrected.

#### Removing repeated cells in multiplane imaging

Three parameters were used to decide if cells from different planes were the same cells: the center distance (center of cell defined as the mean value of all pixel positions belonging to the cell as extracted from Suite2P), the overlap ratio (described below in “Multi-FOV stitching” section), and the Pearson’s correlation of the calcium signals (ΔF/F) between cells in adjacent planes. We manually changed the thresholds for these three parameters. We decided on a threshold of 15 μm for center distance, 50% for overlap ratio and 0.5 for signal correlation for most of the multi-plane data in this paper; this minimized the number of false negatives (repeated cells that were not identified), with an acceptable false positive rate (falsely identified repeated cells). If a cell was detected in multiple planes, only the one with the higher average ΔF/F value was kept. All spatial tuning analyses were performed after repeated cells were removed. Software for removing repeated cells is included in *NATEX* (see Key Resources Table).

#### Multi-FOV stitching

Successful FOV stitching relies on several critical conditions: 1) each individual FOV should be larger than the shifting step so that enough overlapping area is present in each pair of images to perform alignment by matching landmarks; 2) imaging should be at high resolution and distortion-free so that landmarks could be clearly identified in all images and aligned; 3) the stitching adapters should have high precision and reliability, so each shift step covers the same distance; 4) the imaging should have a large enough *z*-scanning range, so that, in each FOV, the same depth from the surface can be calibrated and fixed; and 5) (optionally) clear wide-field imaging of the blood vessel distribution should be available to confirm the accuracy of alignment. MINI2P fulfills these criteria. The FOV of MINI2P is about 500 × 500 µm^2^, larger than the shifting step (400 µm in *x* and *y*), so about 10% (50×500 µm^2^) of adjacent FOVs overlapped and could be used for alignment. We identified single neurons or small blood vessels on both FOVs of overlapping pairs for precise alignment. Each stitching adapter was carefully measured and tested for matching with the microscope. The actual shifting steps were 400 µm±20 µm, and always maintained the required overlap. The *z*-scanning range of MINI2P is 240 µm. We placed the brain’s surface at a focus of 40 µm, i.e. 40 µm above and 200 µm below the brain’s surface were accessible. The two imaging planes in our data, −100 µm and −140 µm, were in the middle of the *z*-scanning range, making it easy to stay at the same focal depth across different FOVs, compensating for varying distances between the objective and the cover glass. Finally, a high-resolution wide-field blood vessel imaging was acquired to achieve precise FOV stitching.

The standard FOV stitching protocol includes three steps (Video S4, Part II, and Supplementary Document, Section 6). First, the neighboring FOVs (after aberration correction) were aligned manually by matching the landmarks either in the overlapping region of neighboring FOVs, or to a wide-field imaging through manual rigid-body shift (rotation and translation). Second, the cross-correlation between the neighboring FOVs in the overlapping region was used to determine the best alignment. The cross-correlation was limited within ±5 pixels in the *x* and *y* directions to avoid divergence. Third, we plotted all cells extracted from Suite2p and checked the overlap of repeated cells. If obviously repeating cells did not overlap well, we re-did steps 1 and 2. If no obvious overlapping cells were found, we skipped step3 to avoid false-positive overlaps. No scaling or shearing was applied to the data. After we confirmed the stitching position of each FOV, the overlap ratio between two cells was defined as:

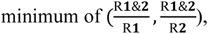

where R_1_ refers to the pixels belonging to cell 1, R_2_ to the pixels belonging to cell 2, and R_1&2_ to the overlapping pixels belonging to both cell1 and cell2 (Figure S3E). The ratio was calculated for each cell with reference to all other cells. Repeated cell pairs were defined as cells with overlap ratios larger than 0.75 (note that this threshold is larger, i.e. more conservative, than the value we used for identifying repeated cells in different planes (0.5), reflecting the fact that the same cells should appear more “similar” in morphology, if viewed in different FOV but still in the same plane, than when they are imaged across different planes). Only one of the two cells in a repeated cell pair – the one with the higher average ΔF/F value - was kept (Figure S3F). The edges of the FOVs were smoothed and merged for better visualization in Figure 3I and Figure 5C and 5D. All spatial tuning analyses were performed after removal of repeated cells. Software for multi-FOV stitching and removal of repeated cells after stitching is included in *StitchingChecker* (see Key Resources Table)

#### Retinotopic mapping

Retinotopic mapping was based on wide-field calcium imaging. Given that retinotopic mapping is an established method, we adapted the system and codes from previous work with minimal modifications (Zhuang et al., 2017). Widefield fluorescence images were acquired with a custom-made microscope (Supplementary Document, Section 7) using a 4x objective (TL4X-SAP, Thorlabs, NJ, USA) and a 100-mm tube lens (AC254-100-A, Thorlabs, NJ, USA). Illumination was provided by a blue LED (M470, NJ, Thorlabs), via a bandpass filter (MF469-35, Thorlabs, NJ, USA), and fluorescence was detected by a CMOS camera (AcA2040-90um, Basler, Ahrensburg, Germany) via a dichroic and bandpass filter (MD499 and MF525-39, Thorlabs, NJ, USA). Parts were mounted on a stereotactic micromanipulator (1760, Kopf, CA, USA). Its optical axis was tilted 20 degrees in the coronal plane such that the optical axis was perpendicular to the cranial window. The microscope’s focal plane was first positioned on the surface of the cortex for blood vessel imaging, then the focal plane was moved 500 µm deeper, defocusing the surface vasculature during the retinotopic mapping. Illumination and image acquisition were controlled with software written in LabVIEW.

During imaging, the animal’s head was restrained by the implanted head bar, eyes were positioned in the horizontal plane, and the body was free to run on a 17 cm diameter slightly tilted disk. Visual stimuli were displayed on a 27” display (Dell U2717D), placed 15 cm from the right eye. The mouse was oriented with its midline at ~30° to the monitor’s plane. We calculated the visual coordinates to the midline (azimuth coordinates) and the horizontal plane through the eyes (altitude coordinates). The monitor covered approximately 0 to 120 degrees in azimuth and −30 to 40 degrees in altitude. Retinotopic maps were generated by sweeping a bar of flickering black-and-white checkerboard stimuli across the monitor (Zhuang et al., 2017). The pattern covered 20 degrees in the direction of propagation and filled the monitor in the perpendicular dimension. The checkerboard square size was 25 degrees. A spherical correction was applied to the checkerboard pattern (Zhuang et al., 2017). Each square alternated between black and white at 6 Hz. To generate a map, we swept the bar across the screen twenty times from down to up and twenty times from left to right, moving at nine degrees per second. A gap of 5 s was inserted between neighboring sweeps, resulting in repetition of the stimulus at 0.048 Hz for vertically-moving stimuli and 0.043 Hz for horizontally-moving stimuli, as descripted by (Zhuang et al., 2017).

During mapping, fluorescence images were acquired at 10 Hz with 4 × binning of the full pixel size of the camera, resulting in an effective pixel size of 11 µm. The camera covered a 5.6×5.6 mm^2^ FOV, which is larger than the chronic window (~4.6 mm). We did not observe any decline in fluorescence during the mapping. All image analyses were performed in a Matlab programming environment. Our retinotopic mapping analysis followed methods described by (Zhuang et al., 2017). The azimuth and altitude position maps from fluorescence versus time data for each pixel were then combined to generate a visual field sign map, which was spatially filtered with a Gaussian kernel (3-pixel width). The visual field sign at each pixel is the sine of the angle between the local gradients in azimuth and altitude. The border of each cortex was drawn manually in software *StitchingChecker* (Video S4, Part II, and Key Resources Table) according to the visual field sign map, which revealed sharp boundaries between cortical regions.

### QUANTIFICATION AND STATISTICAL ANALYSIS

#### Standardized core data processing pipeline

The neural activity and tracking data processing pipeline is packaged in software *NATEX* (see Key Resources Table). It includes three blocks, related to **neuronal activity processing**, **processing of tracking data**, and **alignment and combination of neuronal activity and tracking data**.

#### Neuronal activity processing

The image stack recorded in ScanImage was loaded into Suite2P for motion correction, region of interest (ROI) detection, calcium signal extraction, and deconvolution. The output from Suite2P included two major pieces of information: 1) activity of individual cells and 2) anatomical information about these cells. For precise anatomical alignment of adjacent FOVs (i.e. for FOV stitching) and the quantifications of FOV size and PSFs, the imaging data was unwrapped with a pre-calibrated transformation matrix generated from DistortionCleaner (see also details of this procedure under “Distortion correction”).” The neuronal activity output includes three traces for each detected ROI (putative cell somata), consisting of the raw fluorescence signal *F*_cell_(*t*), the neuropil fluorescence signal F_np_(t), and the deconvolved calcium activity E(t). *F*_cell_(*t*) and F_np_(t) are the weighted averages of activity within and surrounding an extracted ROI respectively, excluding all pixels that overlap with other ROIs. For each cell, the neuropil signal was subtracted from the raw calcium signal using a fixed coefficient (as suggested in (Pachitariu et al., 2018)), yielding the neuropil corrected signal: F_corr_(t) = F_cell_(t)-0.7* F_np_(t). The deconvolved calcium activity E(t) was computed via non-negative deconvolution of F_corr_ with a constant exponential kernel of the decay timescale that matches the value reported in the literature for the calcium sensor we used (1.5s for GcaMP6s) (Friedrich et al., 2017; Pachitariu et al., 2017). The deconvolution increases temporal precision and can denoise the recorded fluorescence signal.

To further diminish the effect that slow fluctuations like photobleaching can have on the fluorescence signal F_corr_, we extracted a running baseline (F_0_(t)). F_0_(t) was estimated as a sum of two components F_0_(t) = F_s_(t) + m as described previously (Low et al., 2014): F_s_(t) denotes the eighth percentile of F_corr_(t) within a ±15-s moving window centered at time *t* and *m* is a constant value added to F_s_(t), such that F_0_(t) is centered around zero in periods without calcium activity (baseline points). These baseline points were extracted as time points for which the local standard deviation (std) of the signal (±15s moving window) did not exceed a cutoff of std_min_+0.1*(std_max_-std_min_), where std_min_ and std_max_ denote the minimal and maximal standard deviations over all data points.

Then calcium activity, expressed as the fractional change in fluorescence with respect to baseline (ΔF/F), was calculated as

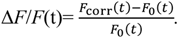

To distinguish calcium transients corresponding to neuronal activity from artifactual fluctuations such as electrical noise and motion artifacts, “significant transients” were identified as described previously (Low et al., 2014). These are transients that occur in periods in which ΔF/F exceeds 2 times the local standard deviation of the baseline fluorescence for more than 0.75s.

Time points spanning significant transients were used to filter both Δ*F*/*F*(t), yielding Δ*F*/*F*(t)_clear_, as well as the deconvolved calcium activity E(t), yielding E(t)_clear_, with signal outside transients set to zero. We normalized the deconvolved and filtered calcium activity E(t)_clear_ by scaling its maximum amplitude to the maximum of ΔF/F_clear_(t) and referred to non-zero incidences of deconvolved calcium activity as “calcium events”.

We next calculated each cell’s signal to noise ratio (SNR, signal/noise), where “signal” is defined as the mean amplitude over all 90^th^ percentiles of ΔF/F(t) in significant transients and “noise” denotes the noise level of ΔF/F(t), calculated as the mean of differences of ΔF/F(t) in periods outside significant transients. A threshold of 3 was set for SNR, and only cells passing this threshold were selected for spatial tuning analysis.

The anatomical information output includes data such as the *x, y* position indicated by pixel number, and *z* position indicated by plane ID, as well as ROI sizes (in pixels) of each individual cell. If data was acquired from multiple planes, we extracted the distance (in *x/y*) between the center of each cell in one plane to each other cell in adjacent planes, as well as their spatial overlap ratio, and the correlation of their calcium signal. Cell pairs with center distances smaller than 15 µm, with more than 50% overlapping pixels, or with signal correlations larger than 0.5, were classified as the same cell (repeated cells). If a second, structural channel was recorded (like tdTomato in MEC data in this study), we used Suite2p’s built-in measurement of probability that the ROI is a cell in second channel (a probability value ranged between 0 and 1) to assess if cells were double labeled (probability close to 1) or not (probability close to 0). In this study we set the cutoff to 0.5, above which a cell was considered double labeled. We repackaged this anatomical information into a dataset named *NeuronInformation.mat* for FOV stitching and anatomical studies. The SNR information of each cell were also storied in *NeuronInformation.mat* for downstream analysis.

#### Tracking data processing

The raw tracking video was first analyzed in DeepLabCut (DLC) (Mathis et al., 2018), which yielded the position of body parts of the animal in each frame. In total, four body parts were extracted: left ear, right ear, body center, and tail base. The DLC model was trained with at least 200 frames from more than ten manually labeled datasets, involving different mice, illumination conditions, and box sizes, to provide robustness against environment variations. The output of DLC includes these body part positions (in pixels) in each frame and a vector with the same length of the video frames ranging from 0 to 1, indicating the confidence of the position estimate (likelikood). We then replaced frames with likelihood < 0.5 with the mean value of one frame before and after. The position of the head was defined as the center of the left ear and right ear position and expressed in *x-y* coordinate: X_head_(t) and Y_head_(t). The traces X_head_(t) and Y_head_(t) were smoothed by computing a linear regression in each window of 0.2 s (3 frames) independently. The momentary moving speed S_head_(t) was then calculated by dividing the moving distance of the head position in two adjacent frames by the frame interval (1/frame rate) (unit: cm/s), and head direction of each frame D_head_(t) is defined as 90 degrees anticlockwise to the direction from the left ear to the right ear.

#### Alignment and combination of neuronal activity and tracking data

Since the 2P imaging and the animal tracking camera are synchronized in hardware, the neuronal activity and animal tracking can be aligned in time. For single-plane imaging, ΔF/F(t), E(t) and other imaging related signals, have the same number of frames as X_head_(t), Y_head_(t) and other tracking related signals, so that each timepoint is pre-aligned. For multiplane imaging (number of planes = X), the neuronal activity data of each cell contains 1/X the number of frames in the tracking data. Thus, the alignment between the neuronal activity and the animal tracking data depends on which imaging plane the cell is from. For example, the neuronal activity of a cell in plane 1 is aligned with every X^th^ tracking datapoint starting with the 1st tracking timepoint, a cell in plane 2 with the X^th^ tracking datapoint starting with the 2^nd^ tracking timepoint.

#### Spatial tuning maps, and head direction tuning maps

Spatial tuning maps were generated to display the neuron’s calcium activity normalized by occupancy. Head direction tuning maps and speed tuning curves were constructed with similar procedures. The value of each bin in the tuning map was obtained as the ratio between the sum of the amplitudes of the deconvolved calcium activity and the time spent in the bin (bin size: 2.5×2.5 cm^2^ for spatial maps, 3° for head-direction tuning functions, and 1.5 cm/s for speed tuning curves), smoothed by a Gaussian filter (standard deviation: 3 cm for spatial heatmaps, 6° for head-direction tuning functions, and 3 cm/s for speed tuning curves). The color code in the spatial tuning map is scaled from zero to the value of the bin with the maximum amplitude of the calcium activity, with ΔF/F·s^-1^ as the unit. To rule out effects of changes in behavioral state (foraging versus sitting still), we dismissed all data at running speeds lower than 2.5 cm/s in all calculations of the observed and shuffled data.

#### Analysis of spatial modulation and extraction of place-modulated cells (PCs) in VC and CA1

For each cell with more than 100 calcium events in each session and SNR>3, we first calculated spatial information based on methods described previously (Skaggs et al., 1996). Spatial information content in bits per ΔF/F·s^-1^ was calculated as

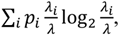

where λ_i_ is the mean amplitude of calcium activity in the *i*-th bin, λ is the overall mean amplitude of calcium activity, and p*_i_* is the probability of the animal’s in the *i*-th bin. A cell was included as a candidate PC if its spatial information exceeded chance level determined by repeated shuffling of the experimental data. Shuffling was performed for each cell individually, with 200 permutations per cell for VC, and 1,000 permutations for CA1. For each permutation, the entire sequence of deconvolved calcium activity of the cell was time-shifted along the animal’s path by a random interval between on one side 30 s and on the other side 30 s less than the length of the session, with the end of the session wrapped to the beginning. Spatial information was calculated for each permutation. If the spatial information from the recorded data was larger than the 95th percentile for spatial information in the distribution from the 200 shuffled data sets in VC and 1,000 shuffled data sets in CA1 of the same cell, the cell was included as a PC candidate. Then we examined the intra-trial place-modulation stability of the PC candidate, defined as the Pearson’s correlation between the spatial tuning map of the calcium activity in the first half-session (0-15 min for 30-min sessions, 0-20 min for 40-min sessions) and in the second half-session (15-30 min for 30-min sessions, 20-40 min for 40-min sessions. If the spatial correlation between tuning maps for the first and second half-session passed the 95th percentile threshold for the same measure in the shuffled versions of the same data, it was kept as a candidate and was passed on to the next filtering step, in which we tested the presence of place fields for each cell. A place field was defined as a connected area (bins) in each spatial tuning map for which the amplitude of mean calcium activity exceeded 20% of the peak calcium activity. Fields were accepted as place fields if their area was larger than 7.5 × 7.5 cm (3 × 3 = 9 bins) and smaller than 62.5 cm × 62.5 cm (or 25 × 25 = 625 bins), and their mean calcium activity exceeded 0.1 ΔF/F·s^-1^ for VC and 0.02 ΔF/F·s^-1^ for CA1. PCs were required to have at least one place field.

#### Analysis of head direction-modulated cells (HD cells) in VC

For each cell with more than 100 calcium events in each session, the length of the mean resultant vector (mean vector length, MVL) was calculated from the head direction tuning curve as described previously (Sargolini et al., 2006). A cell was included as a candidate head direction-modulated cell if the MVL of its directional tuning curve exceeded a chance level determined by repeated shuffling of the experimental data. Shuffling was performed for each cell individually, with 200 permutations per cell. For each permutation, the entire sequence of deconvolved calcium activity of the cell was time-shifted along the animal’s path by a random interval between on one side 30 s and on the other side 30 s less than the length of the session, with the end of the session wrapped to the beginning. An MVL value was calculated for each permutation. If the MVL from the recorded data was larger than the 95th percentile of MVLs in the distribution from the 200 permutations of shuffled data from the same cell, the cell was included as an HD cell candidate. We also examined the intra-trial tuning stability of the HD cell candidate, defined as the Pearson’s correlation between the head direction tuning curve in the first half-session (0-15 min for 30-min sessions, 0-20 min for 40-min sessions) and in the second half-session (15-30 min for 30-min sessions, 20-40 min for 40-min sessions). For an HD cell candidate to be identified as HD cell, its intrasession head direction tuning correlation had to pass the 95th percentile threshold for correlation in the shuffled versions of the same data. The shuffling was performed with the same method as for MVL. The Pearson’s correlation between the head direction tuning curve in the first half-session and in the second half-session was calculated for each shuffling permutation.

#### Comparing the population ratio of functional cell types to chance level

To compare the population ratio of functional cells in our data to the population ratio expected to pass all criteria by chance, we shuffled the deconvolved calcium events of each identified functional cell (in the VC recording in Figure 4, it refers to 293 PCs, or 559 HDCs; in the MEC recording in Figure 5, it refers to 300 MEC grid cells; and in the CA1 recording in Figure 7, it refers to 122 PCs). The procedure was repeated several times per cell (16 or 17 times for PCs in VC, 8 or 9 times for HDCs in VC, 2 or 3 times for grid cells in MEC, and 2 or 3 times for PCs in CA1), until the total number of permutations reached the total number of cells in the respective brain region (4,786 in VC, 796 in MEC, and 254 in CA1). Subsequent to each iteration of shuffling (for all cells), we determined how many cells passed all criteria. This procedure was then repeated 200 times in VC and 1,000 time in MEC and CA1 to obtain a distribution of chance levels (see also Figure S4A).

#### Grid cells in MEC

The structure of the heatmaps of calcium activity was evaluated for all cells with more than 100 calcium events in each session by calculating the spatial autocorrelation for each spatial tuning map (Sargolini et al., 2006). For each cell, a grid score was determined by taking a central circular sample of the autocorrelogram, with the central peak excluded, and comparing rotated versions of this sample. For each sample, we calculated the Pearson correlation of the ring with its rotation in α degrees first for angles of 60° and 120° (group-1) and then for angles of 30°, 90° and 150° (group-2). We then defined the minimum difference between any of the elements in the first group (60° and 120°) and any of the elements in the second (30°, 90°, and 150°). The cell’s grid score was defined as the highest minimum difference between group-1 and group-2 rotations in the entire set of successive circular samples. A cell was defined as a grid cell if its grid score exceeded a chance level determined by the repeated shuffling of the experimental data. Shuffling was performed for each cell individually, with 1,000 permutations per cell. For each permutation, the entire sequence of deconvolved calcium activity by the cell was time-shifted along the animal’s path by a random interval between on one side 30 s and on the other side 30 s less than the length of the session, with the end of the session wrapped to the beginning. The grid score of each shuffled version of the data was calculated based on previously published methods (Langston et al., 2010). If the grid score from the recorded data was larger than the 95th percentile of grid scores in the distribution from the 1,000 permutations of shuffled data of the same cell, the cell was defined as a grid cell. We also examined the cell’s grid spacing and grid orientation to minimize the number of false positives, and only grid cells passing further criteria for spacing and orientation were included (see next section for grid spacing and grid orientation).

#### Grid spacing and grid orientation offset

For fine-grained analysis of the geometric features of the grid cells, we analyzed grid spacing and orientation in the following manner. From the spatial autocorrelograms, we defined individual fields as neighboring bins above a correlation criterion (0-0.5, depending on scale). We then computed the distances between the center of mass of the six inner fields (closest to the autocorrelogram center) with reference to the autocorrelogram center. The grid spacing was defined as the mean distance of these six fields. As spatial autocorrelograms are symmetric, we defined 3 axes for further analysis. We first determined which of the 6 inner fields was closest in absolute angular distance to a fixed horizontal reference (aligned with one of the walls of the recording box) and labeled this Axis 1. The two axes that displayed minimal absolute angular distance to Axis 1 were labeled Axis 2 (most positive) and Axis 3 (most negative). Grid orientation was expressed as the arithmetic angular mean of Axes 1-3. Grid orientation offset was defined as the smallest angle of Axes 1-3 to the north-south wall of the box. To ensure that the number of false positives in the sample of grid cells was minimal, only grid cells with (i) a minimal interaxis angle of >30 degrees, (ii) a maximum interaxis angle of <90 degrees, and (iii) distances of six inner fields did not differ substantially (a ratio between 0.5 and 2 for all fields) were kept for downstream analysis.

#### Statistical tests

All statistical tests were two-sided. We used the Friedman test (a non-parametric alternative to repeated measure ANOVA) with a Tukey post hoc test for all the statistical tests in Figure 1 and Figure S1. For testing the influence of microscope weight and the cable flexibility on the animals’ behavior, the sequence of the experiments was randomized to eliminate learning effects. We used paired Student *t*-tests for group comparisons in Figure 6B and Figure S6B.

