## Supplementary Documents for "Large-scale two-photon calcium imaging in freely moving mice"

**This file includes:**

Supplementary Document

#### Content:

1.Optics of MINI2P miniscope (page 2)

2.Objective drawings and resolution test (page 3)

3. Resolution and FOV measurement in different focal planes (page 4)

4 Stability of µTlens compared to EL-3-10 (page 5-page 6)

5.Simulation of distorted scanning field (page 7)

6.Three-step protocol for accurate FOV alignment. (page 8)

7.Retinotopic mapping: hardware and data. (page 9)

8. Design and construction of HC-920 fiber assembly. (page 10)

9. System wiring and control. (page 11)

10. Materials for assembly of a MINI2P miniscope. (page 12)

11. Shopping & Machining List (page 13- page 24)

#### 1.Optics of MINI2P miniscope

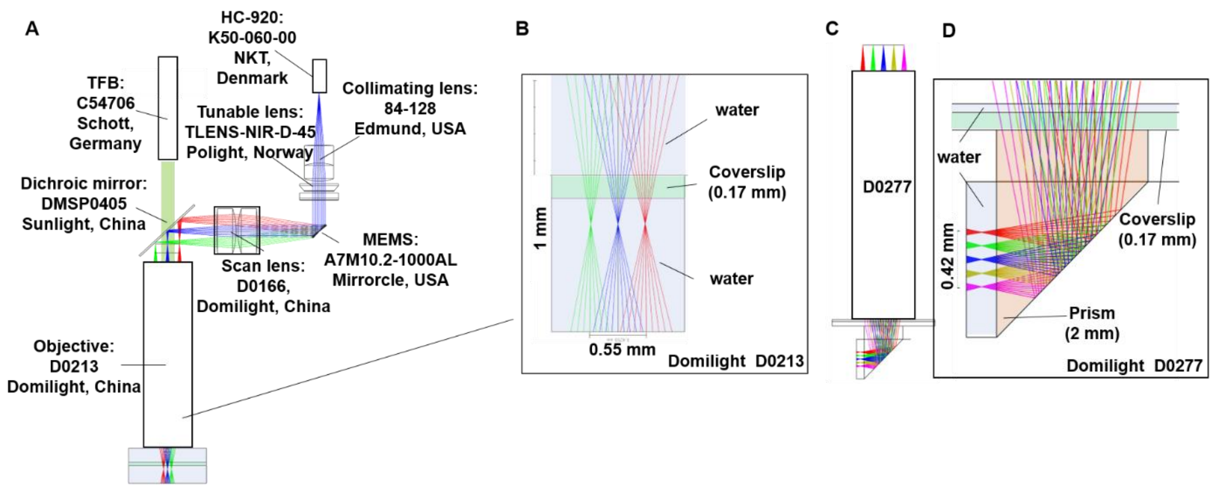

(A) Optical simulation from HC-920 fiber output to sample plane. All optical components shown in A are commercially available. (B) Objective D0213 (water-immersed) was used for illustration. (C and D) Zemax simulation of objective D0277 with 170-µm coverslip and 2-mm prism (material: BK9).

#### 2.Objective drawings and resolution test

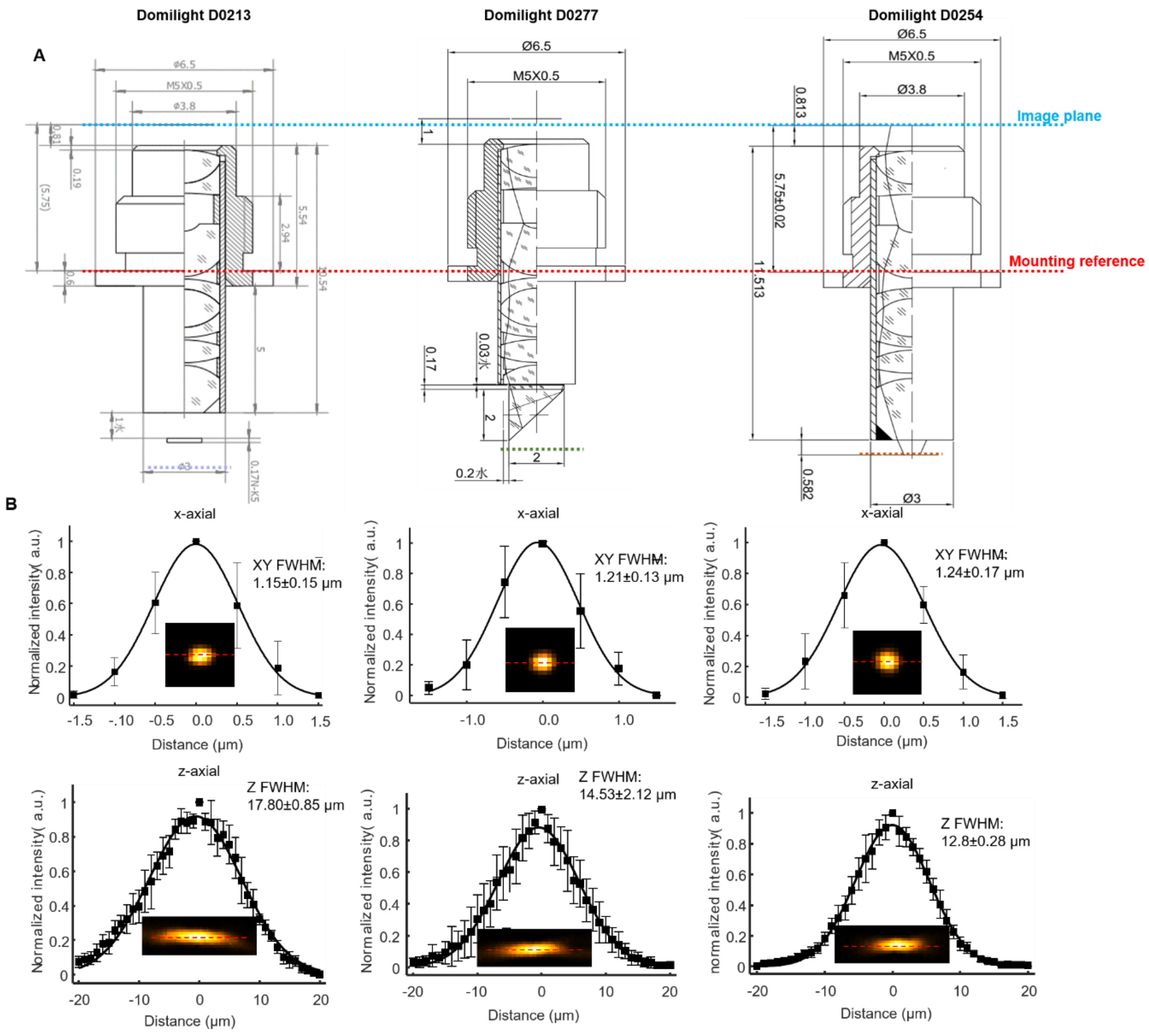

(A) Optical and mechanical design of three objectives. Identical threads and similar distance between mounting reference (red dashed line) and imaging plane (blue dashed line) ensure that the three objectives are interchangeable. (B) Resolution test using MINI2P-L with three objectives. 3D imaging of 1-µm fluorescence beads was used to calculate 3D point-spread-function (PSF) of the microscope. Top: intensity of cross-section along x-axis centered at peak intensity position of beads image (example: dashed line on inserted image). Filled squares indicate recorded data; curve indicates Gaussian fit. Error bars indicate standard deviation of 6 beads data randomly selected from about 400×400 µm$^2$ in the center of FOV. XY FWHM indicates full width at half maximum of the Gaussian fitting. Inserted image: average image of 6 beads in the *xy* plane that the peak intensity located. Bottom: intensity of cross-section along *z*-axis centered at peak intensity position of the beads image (example: dashed line on the inserted image). Z FWHM indicates full width at half maximum of the Gaussian fitting. Inserted image: average image of 6 beads in the *xz* plane that the peak intensity located. *xy* pixel sizes: 780 nm, Stack interval: 1 µm.

#### 3. Resolution and FOV measurement in different focal planes

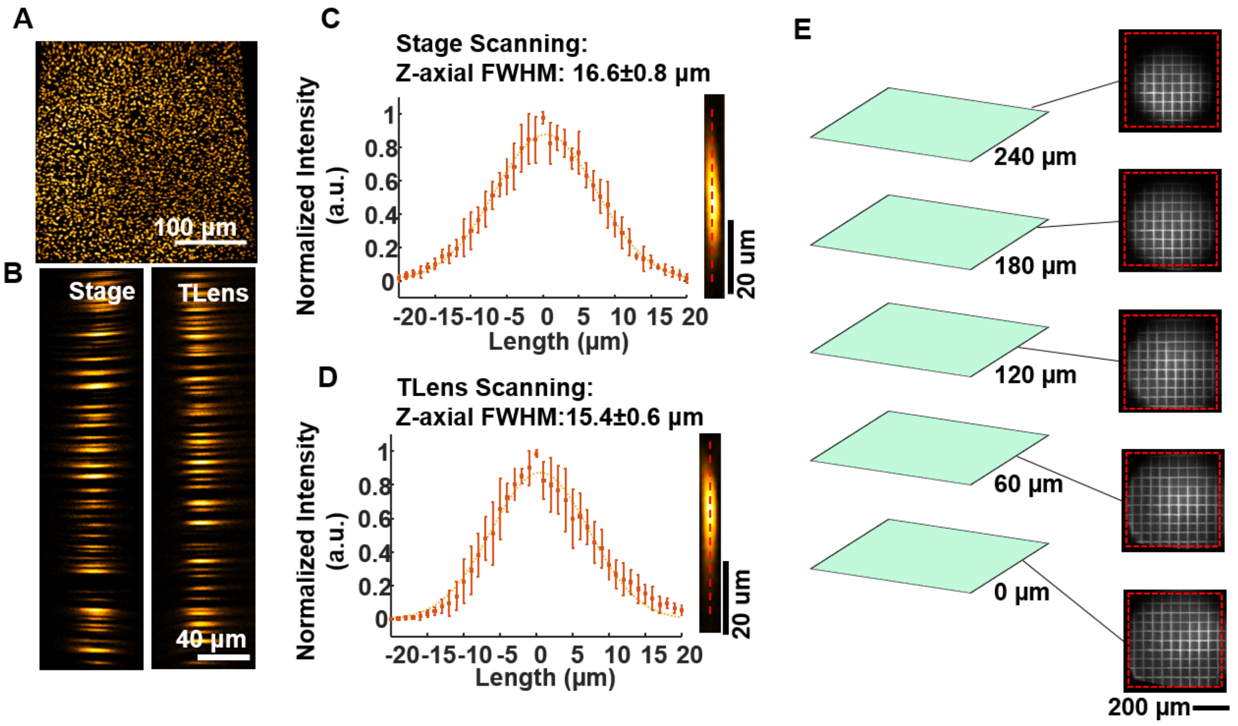

(A to D) Axial resolution of the MINI2P microscope after scanning with quartet μTlens.
(A and B) Imaging of 1-μm fluorescence beads taken by either moving the motorized stage which held the microscope (left in B, labelled as "stage") or changing the focus of the quartet μTlens (right in B, titled as "Tlens").
(A) XY projection of beads image (with μTlens scanning).
(B) XZ projection.
(C and D) Cross-section along *z*-axis centered at the peak intensity position of each bead (dashed line on the right image) was used to calculate the axial full width at half maximum (FWHM) by stage scanning (C) or μTlens scanning (D). Dots indicate recorded data; curve indicates Gaussian fit. Error bars show standard deviation from 10 beads randomly selected over a FOV of 300×300 μm$^2$. *z*-axis FWMHs were extracted from the Gaussian fitting. Right image: average *xz*-projected image of 10 beads.
(E) Usable FOVs of the MINI2P-L in different focus planes. Images of a 50-μm-grid test sample in 5 different focus planes (0 μm to 240 μm) with μTlens scanning. Field distortion has been corrected. Red dashed box indicates a 500×500 μm$^2$ area.

#### 4.Stability of μTlens compared to EL-3-10

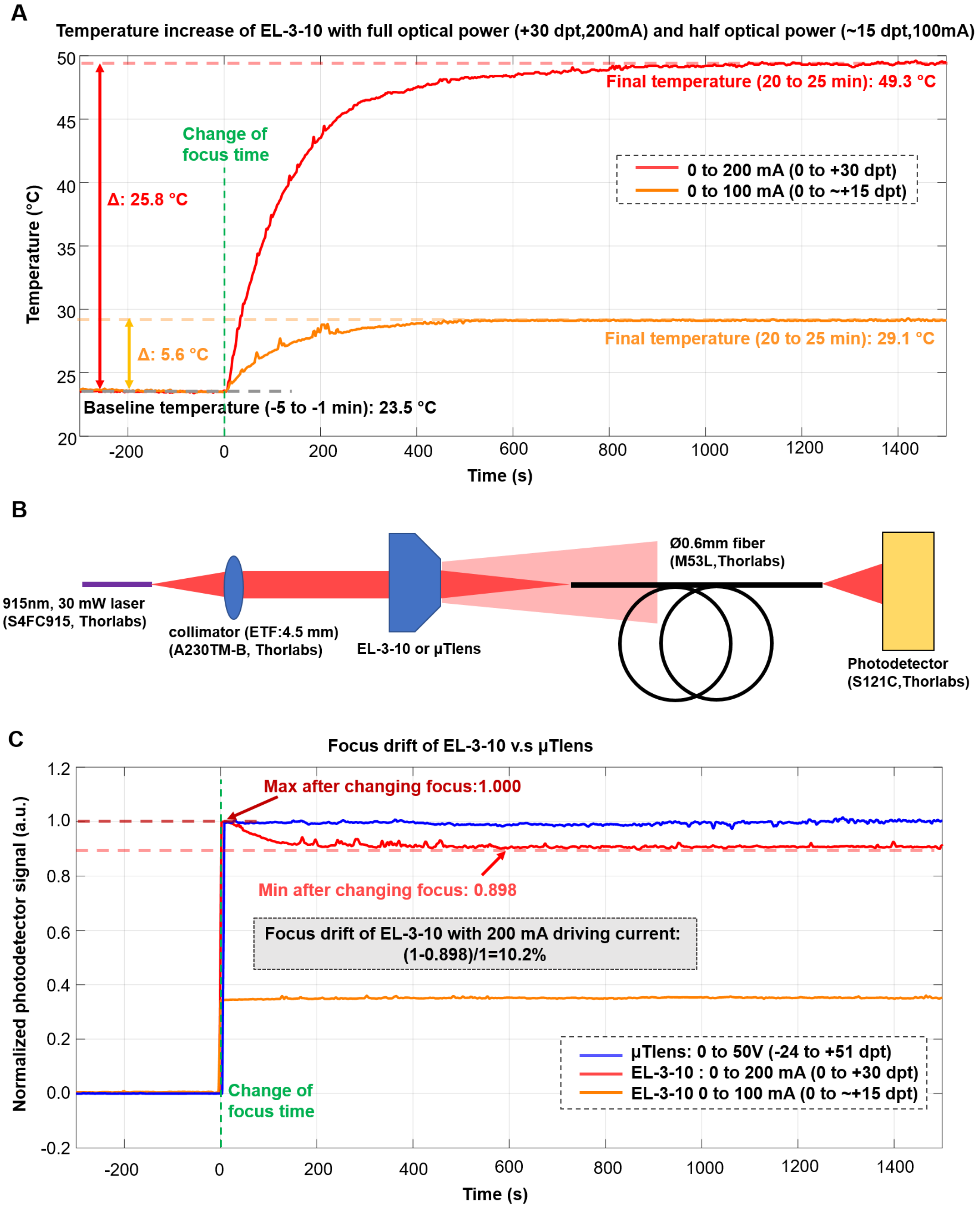

(A) By driving the EL-3-10 with full optical power (+30dpt, 200mA), the temperature increased more than 24 °C compared to the baseline temperature (0 dpt, 0 mA) when no driving current was given to the lens, reaching 48.0 °C within 10 minutes, when it stabilized to 49.3 °C (increase of 25.8 °C) at 20 to 25 minutes. By driving the EL-3-10 with half optical power (about +15dpt,

100mA), a final temperature of 29.1°C was reached after 10 to 15 minutes, thus increasing only 5.6 °C. The temperature of the ETL (EL-3-10) was measured by attaching a 10 kΩ thermistor (TH10K, Thorlabs, NJ, USA) on the shell of the lens. The baseline temperature was calculated by averaging temperature measurements from 5 min to 1 min before the focus was changed (green line, from 0 mA to 200 mA or 100 mA). The final temperature was calculated by averaging temperature measurements from 20 min to 25 min after the focus was changed (green line, from 0mA to 200 mA or 100 mA). Temperature was measured every 0.5 s.
(B) System for measuring the relative focus of EL-3-10 and µTlens (quartet). The system is identical to that of Zong et al. (2021).
(C) Relative focus drift, defined as the normalized difference between the maximum and minimum photodetector signal after the focus is changed (green line), was 10.2% for EL-3-10 with full optical power (+30dpt, 200mA), 1.4% for EL-3-10 with half optical power (~+15dpt, 100mA), and 2% for µTlens(quartet) with full optical power (75dpt, 50V). Grey box shows calculation of the relative drift of EL-3-10 with full optical power. The relative drift of EL-3-10 with half optical power, and that of the µTlens(quartet) with full optical power, were calculated in the same way. Note substantially larger focus drift of EL-3-10 compared to µTlens(quartet) with full optical power.

#### 5.Simulation of distorted scanning field

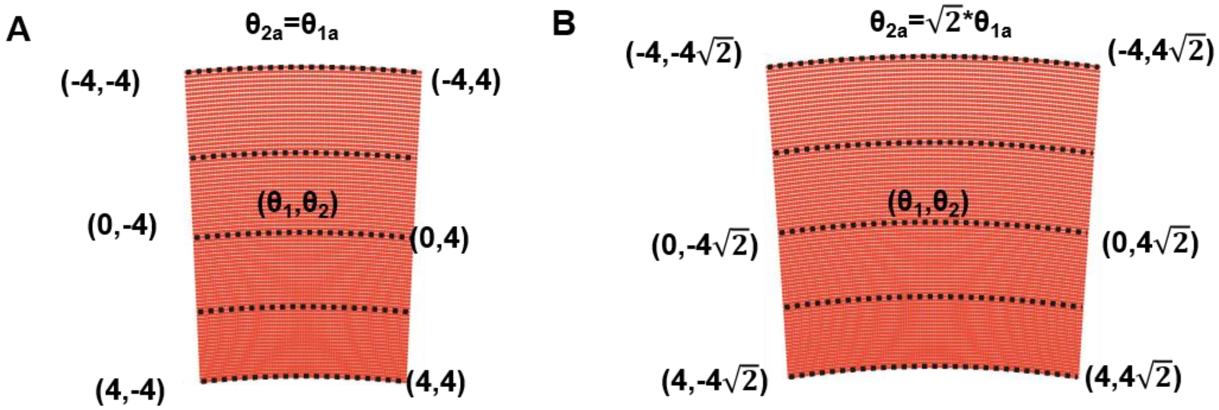

Scanning fields without distortion correction.

(A) Scanning field when fast axis and slow axis have the same scanning angle.

(B) Scanning field when scanning angle of the fast axis is √2 of the slow axis.

#### 6.Three-step protocol for accurate FOV alignment.

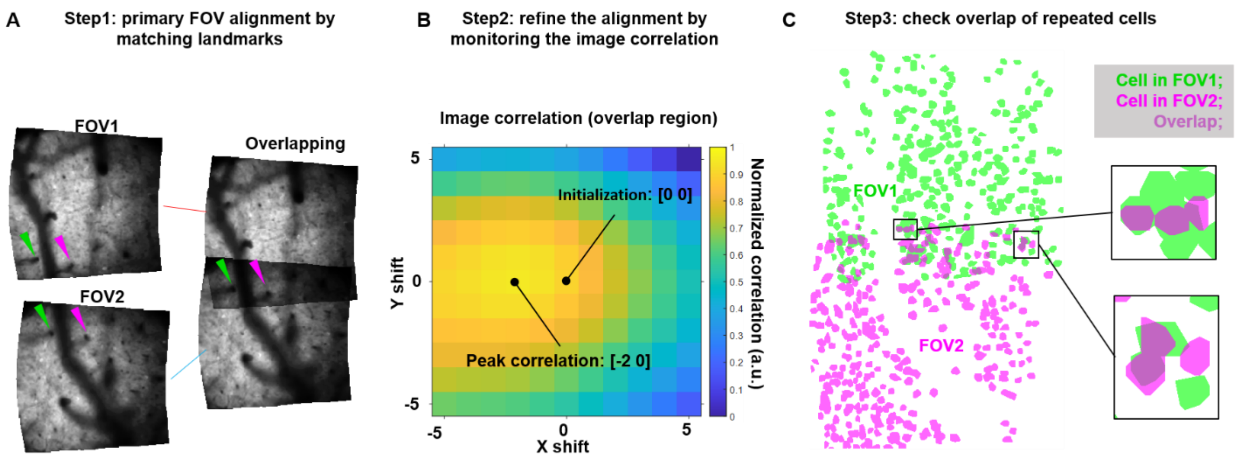

(A) Step1: primary FOV alignment based on overlapping landmarks. If wide-field imaging was not available, neighboring FOVs were first manually aligned by identifying overlapping landmarks on the averaged images. Left: green and purple arrows show two landmarks on both FOVs that were used for registration. Right: Overlap of the same two FOVs after stitching.

(B) Step2: refinement of the alignment by monitoring the image correlation: a small shift was made in the *x* and *y* directions (-5 pixels to 5 pixels) and the position of peak image correlation was identified and used to refine the alignment.

(C) Step3: Final check of alignment by assessing overlap between repeated cells (cells present in both FOVs): Suite2P extracted cells ROI are colored according to the FOV they belonged to and overlaid in the same image (green: FOV1, purple: FOV2, merged color: overlapping pixels). Repeated cells across neighboring FOVs should have large overlap values if the alignment is precise. If this was not the case, Step1 and Step2 would be reinitiated.

#### 7.Retinotopic mapping: hardware and data.

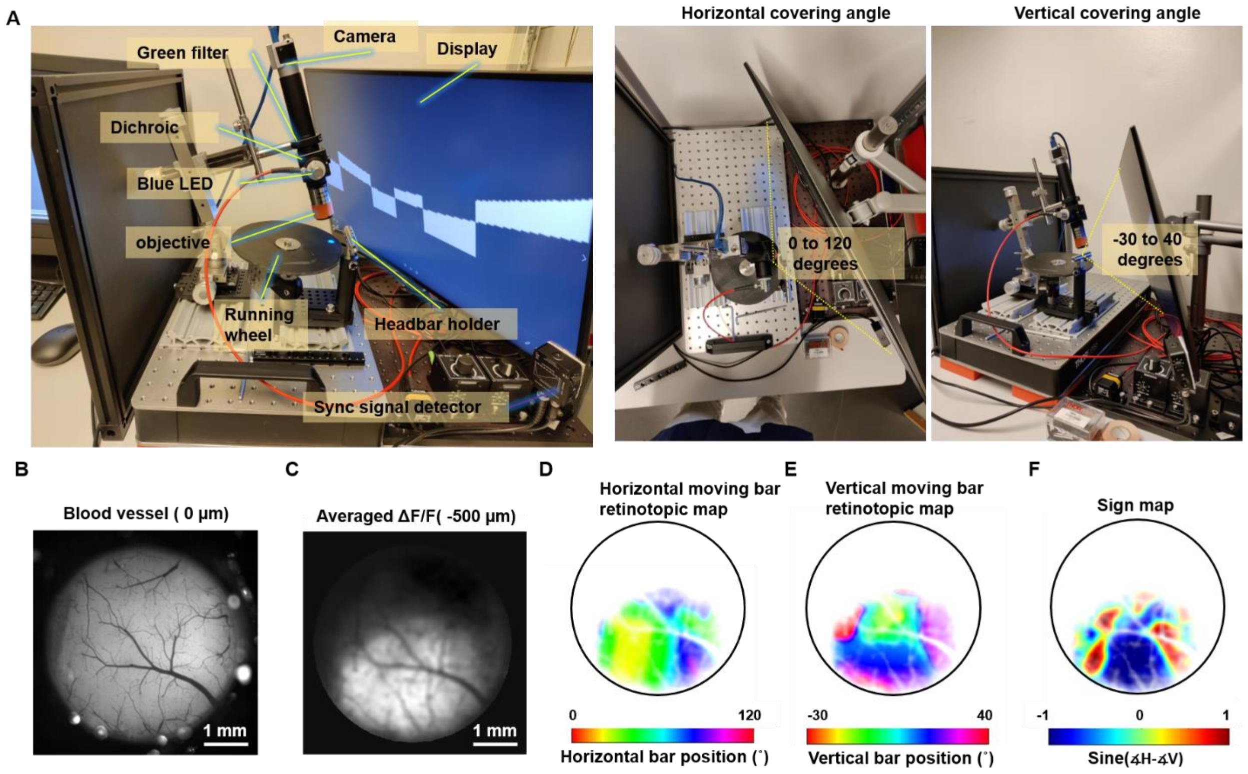

(A) Pictures of the visual stimulation system for retinotopic mapping on visual cortices.

(B) Wide-field image of blood vessels on entire chronic window (4.6 mm).

(C) Average intensity of all images highlights visual cortices as the area with maximum response to the visual stimulation.

(D) Extracted horizontal retinotopic map.

(E) Extracted vertical retinotopic map.

(F) Sign map calculated from (D) and (E) for determining borders of visual cortices. Color indicates the sine value of the angle between vertical (V) and horizontal (H) retinotopic mapping gradients (red:1; blue: -1). See Methods and Materials for details.

#### 8. Design and construction of HC-920 fiber assembly.

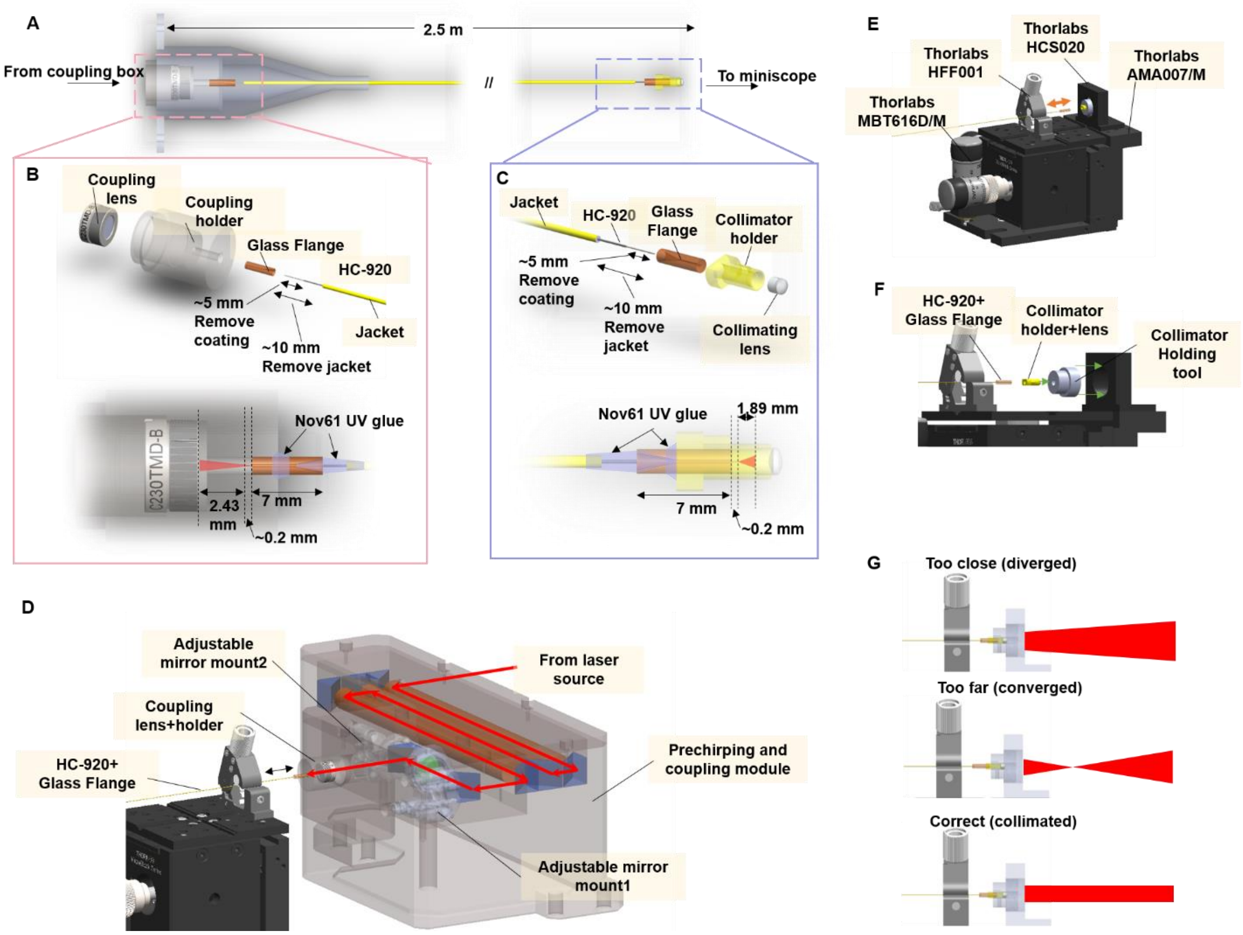

(A-C) Schematic of the HC-920 fiber assembly.

(A) The whole assembly consists of (B) a fiber coupler, a HC-920 fiber and (C) a fiber collimator.

(B) Components of the fiber coupler.

(C) Components of the fiber collimator.

(D) Illustration showing the fiber alignment stage (the black device on left) for making of fiber coupler and coupling laser into HC-920.

(E-G) Illustration showing making of the fiber collimator.

(E-F) Tools for aligning and collimating the fiber collimator.

(G) Adjusting distance between HC-920 fiber and collimating lens until the output beam is collimated.

All 2D drawings and 3D models are in Github: https://github.com/kavli-ntnu/MINI2P

### 9. System wiring and control.

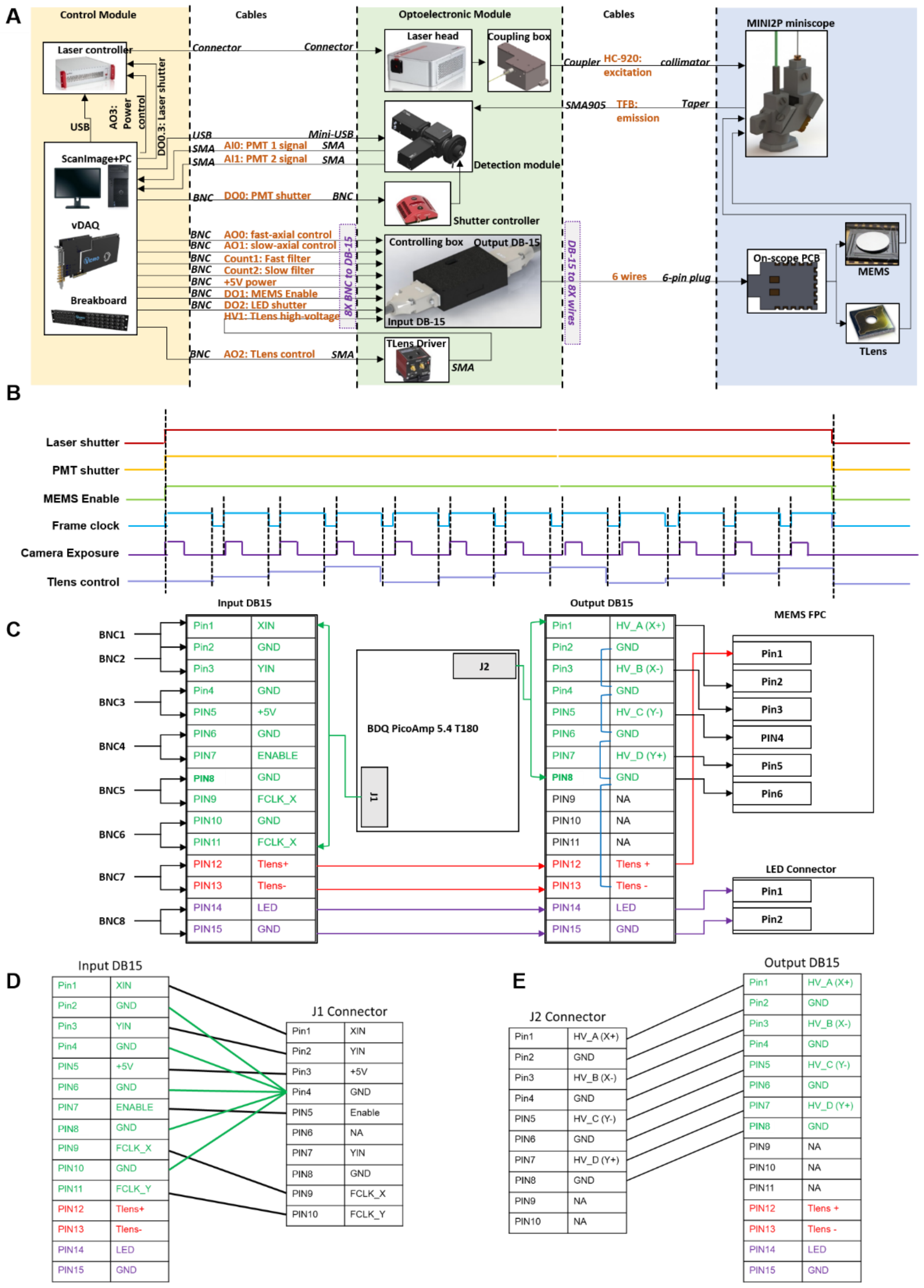

(A) Schematic of electrical hardware and wiring.

(B) Timing of control signals.

(C-E) Details of wiring of the controlling box, and the connection from the control box to the microscope.

#### 10. Materials for assembly of a MINI2P miniscope.

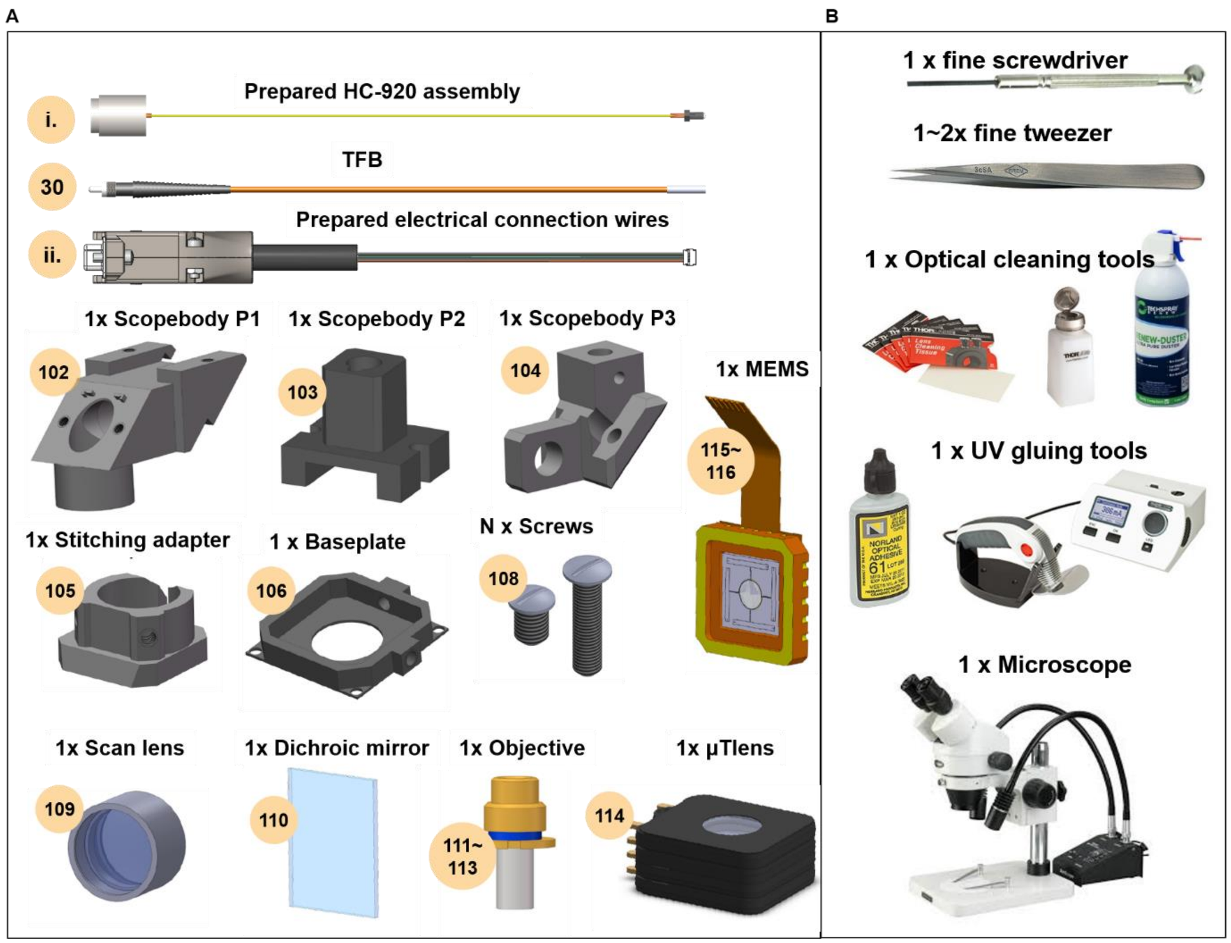

(A) All required components for assembling one MINI2P miniscope.

(B) All required equipment and tools for assembling one MINI2P miniscope (see Section 11 for details).

#### 11. Shopping & Machining List

| Item ID | Source | Access | Description | Link | Image | Quantity |
|---|---|---|---|---|---|---|
| | | | | | | **General tools** |
| a. | Thorlabs | purchase | Spanner Wrench 1 | SPW30 | 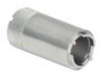 | 1 |
| b. | Thorlabs | Purchase | Fiber stripping tool | FTS4 | 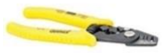 | 1 |
| c. | Thorlabs | purchase | Fiber Cleaver | XL411 | 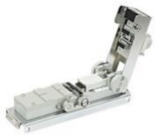 | 1 |
| d. | Thorlabs | purchase | Spanner Wrench 2 | SPW602 | 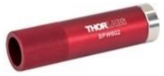 | 1 |
| e. | Thorlabs | purchase | UV Curing LED System | CD20K2 | 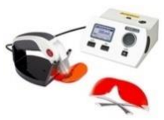 | 1 |
| f. | Thorlabs | purchase | Handheld laser source (630nm) | HLS635 | 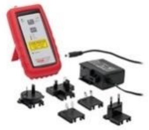 | 1 |
| g. | Thorlabs | purchase | Power and Energy meter | PM100D | 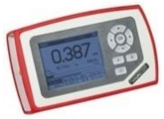 | 1 |

| | | | | | | |
|---|---|---|---|---|---|---|
| h. | Thorlabs | purchase | Green LED | M530F2 | 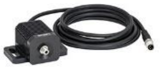 | 1 |
| i. | Thorlabs | purchase | NIR Detector Card | VRC4 | 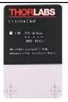 | 1 |
| **Core optics module** | | | | | | |
| **Mechanical components** | | | | | | |
| 1 | Thorlabs | purchase | Nexus Breadboard | B3045L | 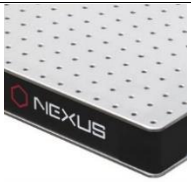  | 1 |
| 2 | Thorlabs | purchase | Sorbothane Feet | AV6/M | 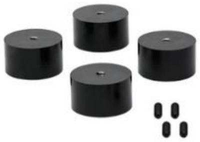 | 1 |
| 3 | Thorlabs | purchase | Fiber Adapter Plate | SM1SMA | 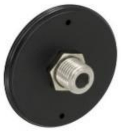 | 1 |
| 4 | Thorlabs | purchase | Adjustable Mirror Mount | POLARIS-K05S2 | 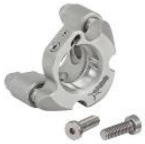 | 2 |
| 5 | Thorlabs | purchase | Rotation Mount | MRM05/M | 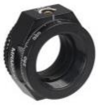 | 1 |
| 6 | Thorlabs | purchase | M6 cap screws kit | HW-KIT2/M | 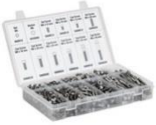 | >20 |
| 7 | Thorlabs | purchase | Ø900 µm Hytrel Furcation Tubing | FT900Y | 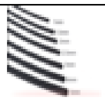 | 3 |
| 8 | Thorlabs | purchase | 3 axis Microblock stage | MBT616D/M | 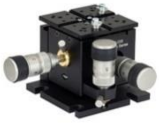 | 1 |

| | | | | | | |
|---|---|---|---|---|---|---|
| 9 | Thorlabs | Purchase | 3-Axis MicroBlock Stage | MBT602/M | 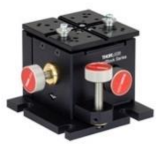 | 1 |
| 10 | Thorlabs | Purchase | Compatible Flexure Stage Mount | HCS020 | 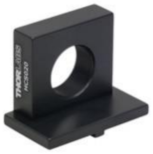 | 1 |
| 11 | Thorlabs | Purchase | Fiber Clamp Holds fiber | HFF001 | 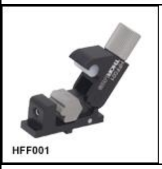  | 3 |
| 12 | Thorlabs | purchase | Fixed mounting bracket | AMA007/M | 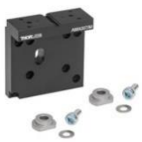 | 1 |
| 13 | Thorlabs | purchase | SM05 Threaded Adapter to laser source | AD1109F | 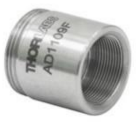 | 1 |
| 14 | Thorlabs | purchase | Kinematic Cage cube | DFM1T3 | 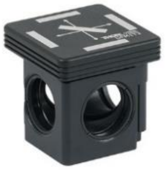 | 1 |
| 15 | Thorlabs | purchase | Kinematic cage cube base | DFM1B | 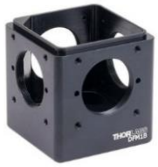 | 1 |
| 16 | Thorlabs | purchase | Lens Tube | SM1L05 | 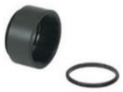 | 3 |
| 17 | Thorlabs | purchase | End Cap for Machining | SM1CP2M | 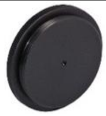 | 1 |
| 18 | Thorlabs | purchase | Coupler | SM1T2 | 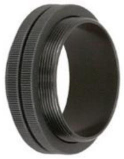 | 3 |
| 19 | Thorlabs | purchase | Optical Beam shutter with controller for PMT | SHB1 | 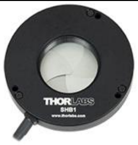 | 1 |

| | | | | | | |
|---|---|---|---|---|---|---|
| 20 | Thorlabs | purchase | Post mounting adapter | SHM1/M | 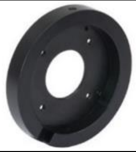 | 1 |
| 21 | Kavli NTNU | Self-made | Coupling box | Github(Kavli-ntnu, 2021) | 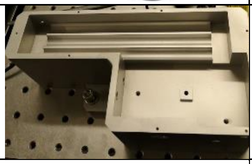 | 1 |
| 22 | Kavli NTNU | Self-made | Box cup | Github(Kavli-ntnu, 2021) | 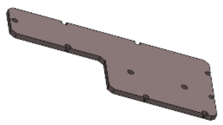 | 1 |
| 23 | Kavli NTNU | Self-made | Coupling holder | Github(Kavli-ntnu, 2021) | 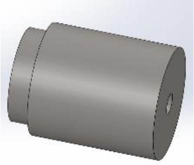 | 1 |
| 24 | Kavli NTNU | Self-made | Coupling protector | Github(Kavli-ntnu, 2021) | 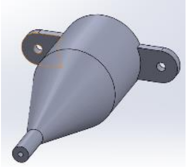 | 1 |
| 25 | Kavli NTNU | Self-made | Collimator assembly tool | Github(Kavli-ntnu, 2021) | 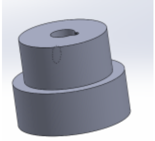 | 1 |
| 26 | Kavli NTNU | Self-made | Collimator Holder | Github(Kavli-ntnu, 2021) | | 1 |
| 27 | Kavli NTNU | Self-Made | Control box shell | Github(Kavli-ntnu, 2021) | 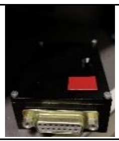 | 1 |
| 28 | Kavli NTNU | Self-Made | Control box cup | Github(Kavli-ntnu, 2021) | | 1 |
| **Optical components** | | | | | | |
| 29 | Toptica | purchase | Laser source | FemtoFiber ultra 920 | 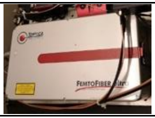 | 1 |
| 30 | Schott | purchase | Tapered fiber bundle (TFB) | C54706 | NA | 1 |
| 31 | NKT Photonics | purchase | Hollow-core PCF, HC-920 | K50-060-00 | NA | 1 |

| | | | | | | |
|---|---|---|---|---|---|---|
| 32 | Sunlight | purchase | Glass rods (ZF62) | GLA-10x150-AR800-1100 | NA | 3 |
| 33 | Sunlight | purchase | Glass flange $\emptyset 0.126x6.5mm$ | TUB-1.8x6.5-0.127 | NA | 2 |
| 34 | Thorlabs | purchase | Prisms direct light from the seed laser up to the HC-920 fiber. | MRA12-P01 | 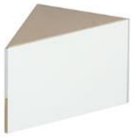 | 8 |
| 35 | Thorlabs | purchase | Emission filter 525 nm green channel | MF525-39 | 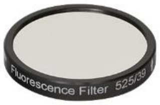 | 1 |
| 36 | Thorlabs | purchase | Emission filter 630 nm red channel | MF630-69 | 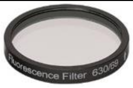 | 1 |
| 37 | Thorlabs | purchase | Shortpass Filter | FESH0750 | 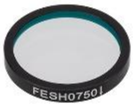  | 2 |
| 38 | Thorlabs | purchase | Dichroic Mirror | DMLP567R | 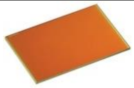 | 1 |
| 39 | Thorlabs | purchase | Aspheric Condenser Lens | ACL25416U-A | 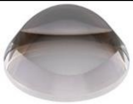 | 3 |
| 40 | Thorlabs | purchase | Coupling lens | C230TMD-B | 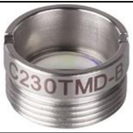 | 1 |
| 41 | Thorlabs | purchase | Half-Wave Plate | WPHSM05-915 | 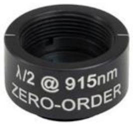  | 1 |
| 42 | Thorlabs | purchase | Protected Silver Mirrors | PF05-03-P01 | 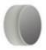 | 2 |
| 43 | Edmunds | Purchase | Collimating Lens | #84-128 | 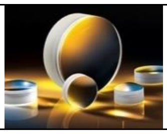 | 1 |
| **Electrical components** | | | | | | |

| | | | | | | |
|---|---|---|---|---|---|---|
| 44 | Mirrorcle Technologies | Purchase | MEMS driver (controller) BDQ PicoAmp 5.4 B160 | DR-11-054-00 | 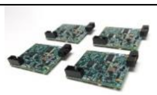 | 1 |
| 45 | Digikey | Purchase | BNC cables | CCBNS-MM-RG174-36 | 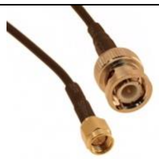 | 8m |
| 46 | RS | purchase | DSUB15 connector plug (male) | 472-859 | 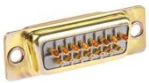 | 2 |
| 47 | RS | purchase | Backshell | 765-9448 | 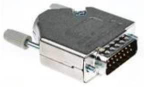 | 2 |
| 48 | RS | | DSUB15 Connector socket (female) | 472-865 | 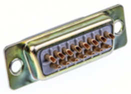 | 2 |
| 49 | Digikey | purchase | 6-pin connector for MEMS | FH19C-6S-0.5SH(10) | 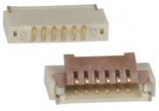 | 1 |
| 50 | Industrifil | purchase | Single wire cables | link | 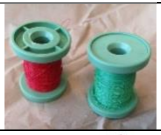 | 6 |
| 51 | Thorlabs | purchase | K-Cube Piezo Controller | KPZ101 | 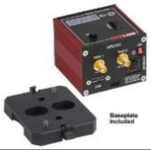 | 1 |
| 52 | Thorlabs | purchase | PMTs | PMT2101/M | 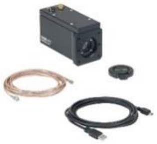 | 2 |
| 53 | Thorlabs | purchase | Controller for shutter | SHB1T | 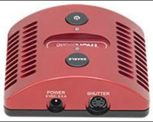 | 1 |
| **Scope mounting module** | | | | | | |
| **Mechanical components** | | | | | | |
| 54 | Thorlabs | Purchase | Aluminum optical breadboard | B3060A | 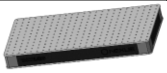 | 1 |

| | | | | | | |
|---|---|---|---|---|---|---|
| 55 | Thorlabs | Purchase | Sorbothane Feet Ø45.0 mm, Internal M6 Mounting Thread, 4 Pieces to support previous breadboard. | AV6/M | 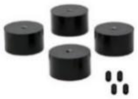 | 1 |
| 56 | Thorlabs | Purchase | One-sided Construction Rail, Clear Anodized, 95x500 mm | XT95SP-500 | 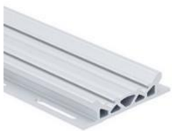 | 2 |
| 57 | Thorlabs | purchase | Rail carriage suitable for sliding onto previous rail and with M6 tapped holes | XT95RC4/M | 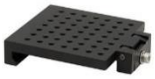 | 2 |
| 58 | Thorlabs | Purchase | Mounting Post Ø1.5" L=350mm | P350/M | 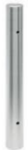 | 1 |
| 59 | Thorlabs | Purchase | Post Mounting Clamp | C1545/M | 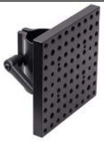 | 1 |
| 60 | Thorlabs | purchase | Manual Rotation Stage | RP03/M | 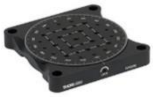 | 1 |
| 61 | Thorlabs | Purchase | Studded Pedestal Base Adapter | BE1/M | 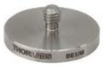 | 1 |
| 62 | Thorlabs | purchase | Optical Post | TR75/M | 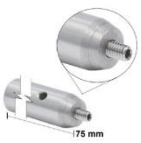  | 2 |
| 63 | Thorlabs | Modified from product | Running wheel hardboard | Github, TB4 | 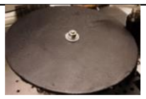 | 1/4 |
| 64 | Thorlabs | purchase | M6 spacers & washers | NA | 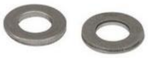 | >4 |

| | | | | | | |
|---|---|---|---|---|---|---|
| 65 | Thorlabs | purchase | Spacer on both sides of wheel | PS1M | 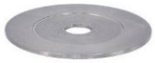 | 2 |
| 66 | SKF | purchase | Bearing OD 19mm ID 6mm | 626-2Z | 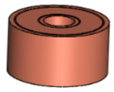 | 1 |
| 67 | Thorlabs | purchase | Right-angle Clamp | RSA90/M | 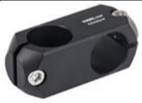 | 1 |
| 68 | Thorlabs | purchase | Pillar posts | RS50/M | 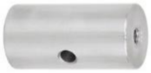 | 2 |
| 69 | Thorlabs | purchase | Universal Post holder | UPH100/M | 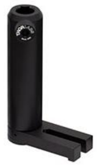 | 3 |
| 70 | Thorlabs | purchase | Locking Ball and Socket Mount; to support LEDs near Tracking Camera | TRB1/M | 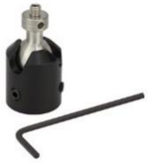 | 8 |
| 71 | Thorlabs | purchase | Adapter camera-lens | SM1A10Z | 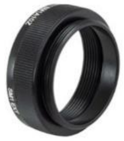 | 2 |
| 72 | Kavli NTNU | Self-made | MINI2P holder P1 | Github(Kavli-ntnu, 2021) | 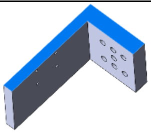 | 1 |
| 73 | Kavli NTNU | Self-made | MINI2P holder P2 | Github(Kavli-ntnu, 2021) | 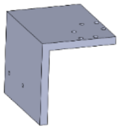 | 1 |
| 74 | Kavli NTNU | Self-made | MINI2P holder P3 | Github(Kavli-ntnu, 2021) | 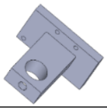 | 1 |
| 75 | Kavli NTNU | Self-made | MINI2P holder P4 | Github(Kavli-ntnu, 2021) | 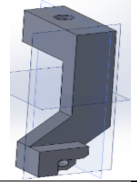 | 1 |
| 76 | Kavli NTNU | Self-made | Wheel holder | Github | 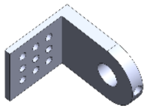 | 1 |

| | | | | | | |
|---|---|---|---|---|---|---|
| 77 | Kavli NTNU | Self-made | Headbar holder | Github | 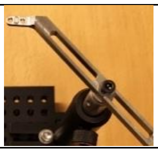 | 1 |
| **Optical and electrical components** | | | | | | |
| 78 | Thorlabs | purchase | 850nm IR LED Array Light Source | LIU850A | 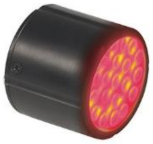 | 1 |
| 79 | Thorlabs | purchase | CMOS Sensor camera | CS165MU/M | 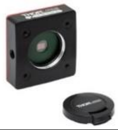 | 1 |
| 80 | Edmund Optics | purchase | Lens focal length 4.5mm Or 8.5mm for lateral and frontal camera to control head fixation | F1.3 f8.5 2/3” | 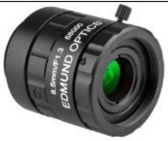 | 1 |
| 81 | Edmund Optics | purchase | Basler camera for Animal Tracking (top or bottom view) | acA2040-90um | 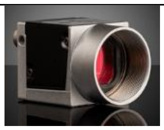 | 1 |
| 82 | Physik Instrumente (PI) | purchase | DC Motors | M-112.2DG | 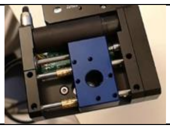 | 3 |
| **Mobile cart and controlling system** | | | | | | |
| **Mechanical and electrical components** | | | | | | |
| 83 | Thorlabs | Purchase | Mobile Cart | POC001 | 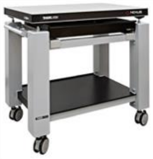 | 1 |
| 84 | Thorlabs | Purchase | Optical breadboard 600x900x55mm | PBG52506 | 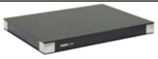 | 1 |
| 85 | Thorlabs | Purchase | Optional drawer | POD001 | 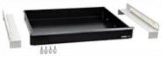 | 1 |
| 86 | Schroff | Purchase | 19-inch rack | Ref 721-2708 | 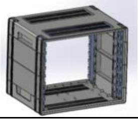 | 1 |

| | | | | | | |
|---|---|---|---|---|---|---|
| 87 | McMASTE R-CARR | Purchase | Span-in Nuts | 90680A729 | 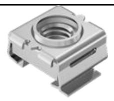 | 10 |
| 88 | Dell | Purchase | Workstation is an Intel core i9 with operating windows 10 Pro | 7080 | 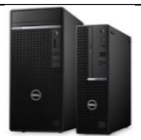 | 1 |
| 89 | Dell | Purchase | Monitors | NA | NA | 1 |
| 90 | Vidrio Technologie s LLC | Purchase | vDAQ card provides data acquisition and control of Laser, TLens, shutter, among others | V-vDAQ.R1 | 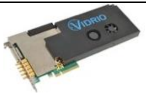 | 1 |
| 91 | Vidrio Technologie s LLC | Purchase | vDAQ breadboard | V-vDAQ.R1 | 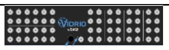 | 1 |
| 92 | Physik Instrumente (PI) | Purchase | Motion Controller for PI motors | C-884.4DC | 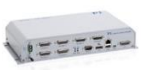 | 1 |
| 93 | Toptica | Purchase | Controller for 920nm laser | Toptica | 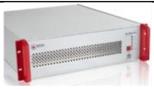 | 1 |
| 94 | Dustin | Purchase | USB-hub ports | Deltaco | 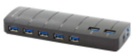 | 1 |
| 95 | Thorlabs | Purchase | BNC Male to BNC Male | CA3136 | 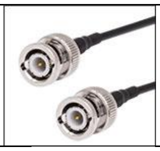 | 8 |
| 96 | Thorlabs | Purchase | BNC to SMA Male Connector | CA2848 | 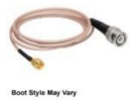  | 2 |
| 97 | Thorlabs | Purchase | SMC connector | PAA101 | 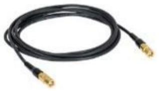 | 2 |
| 98 | Thorlabs | Purchase | SMA-to-SMA cable | CA2912 |  | 2 |

| | | | | | | |
|---|---|---|---|---|---|---|
| 99 | Thorlabs | Purchase | Power supply for Tlens driver | TPS002 | 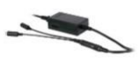 | 1 |
| **Software** | | | | | | |
| 100 | Vidrio Technologies LLC | Purchase | Open-source software for the whole system | ScanImage 2020 | NA | 1 |
| 101 | Others | Free or purchase | See Document S4 | NA | NA | 1 |
| **MINI2P miniscope** | | | | | | |
| **Mechanical components** | | | | | | |
| 102 | Kavli NTNU | Self-made | Scope Body P1 | GitHub | | 1 |
| 103 | Kavli NTNU | Self-made | Scope Body P2 | GitHub | | 1 |
| 104 | Kavli NTNU | Self-made | Scope Body P3 | GitHub | | 1 |
| 105 | Kavli NTNU | Self-made | Stitching adapter | GitHub | | |
| 106 | Kavli NTNU | Self-made | Baseplate | GitHub | | 10 |
| 107 | Kavli NTNU | Self-made | Alignment tool | GitHub | 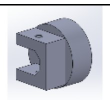 | 1 |
| 108 | TR fastenings | Purchase | Screws for MINI2P, M1.4; M1.6; M2.0; | NA | | >10 |
| **Optical components** | | | | | | |
| 109 | Domilight | Purchase | Scan lens | D0166 | | |
| 110 | Sunlight | Purchase | Dichroic mirror | DMSP0405 | | |
| 111 | Domilight | Purchase | Objective1 | D0213-3X | | 1 |

| | | | | | | |
|---|---|---|---|---|---|---|
| 112 | Domilight | Purchase | Objective2 | D0254-3X | | 1 |
| 113 | Domilight | Purchase | Objective3 | D0277-3X | | 1 |
| 114 | Polight | Purchase | µTLENS | TLENS-NIR-D-45 | | 1 |
| **Electrical components** | | | | | | |
| 115 | Mirrorcle | Purchase | MEMS “fast” 5600 Hz | Fast - A7M10.2-1000AL | 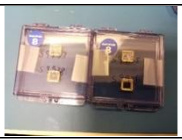 | 1 |
| 116 | Mirrorcle | Purchase | MEMS “slow” 2000 Hz | A3I12.2-1200AL-LCC20-TW | | 1 |
| 117 | Kavli NTNU | Purchase | MEMS PCB | | 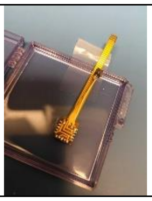 | 1 |
